# Mast fruiting in a large tropical African legume tree species and the mineral nutrient limitation hypothesis

**DOI:** 10.1101/2023.04.03.535348

**Authors:** David M. Newbery, Sarah Schwan, George B. Chuyong, Godlove A. Neba, Culbertson E. Etta, Julian M. Norghauer, Martin Worbes

## Abstract

The large grove-forming ectomycorrhizal tropical tree *Microberlinia bisulcata* displays strong mast fruiting on poor soils. Evidence is presented that masting is probably determined primarily by mineral nutrients and secondarily by radiation. To test the nutrient resource limitation hypothesis, phenological recordings were matched with climate variables and modelled using logistic time-series regression. A strong predictor was average daily rainfall in the dry season, low in the current year of masting and high in the year prior. Less strong was the increase in dry season radiation between prior and mast years. Masting events showed no relationship to annual stem increment. Masting intensity was reduced after two caterpillar attacks. Fallen-leaf phosphorus concentration increased in the inter-mast interval and that for potassium peaked mid-interval. A drier dry season prior to masting may have increased tree carbon, a wetter one increased nutrient uptake by roots and mycorrhizas. A rooting–fruiting trade-off in carbon is suggested. The long-term stochastic driver appears to be dry-season rainfall. Synchrony across masting trees may be enhanced by equilibration of nutrients across the mycorrhizal-root network. Life history strategy determined by mineral nutrient resource dynamics provides the best explanation and advanced model of the observed pattern of events.

## INTRODUCTION

By fruiting and seeding intensively at intervals as opposed to continuously over the years, seed and seedling survival may be enhanced for plants species with a masting iteroparous type of life-history as an evolutionary stable strategy [1–3]. Trees provide by far the majority of the documented cases. To explain how a species’ fitness can be maintained or increased, a so-called ‘economy of scale’ concept has often been invoked [4]. Three general processes have been proposed to achieve this type of economy, usually thought of, teleonomically, as the ultimate causes of mast fruiting and seeding: (1) the efficacy of pollination may be increased by mass flowering, (2) seed predators may be more readily satiated by mass seeding, and (3) the chances of successful dispersal and establishment may be improved by larger reproductive output [5–8]. Much of the reported research into masting, however, has been in the ecological and evolutionary direction with a strong temperate-biome bias (see reviews aforementioned). Surprisingly, there have been comparatively few studies on masting in the tropics [e.g., 9, 10-12].

A much more critical issue for understanding masting in trees is the present lack of evidence for, and hypotheses about, the physiological mechanisms which *enable* mast fruiting and seeding, for both temperate and tropical ecosystems. These mechanisms are commonly viewed by ecologists as being proximate causes. Recent discussions about whether a strict distinction exists between ultimate and proximate causes [13, 14] leave it more open that factors, such as nutrient supply, directly affecting physiological processes might also qualify as ultimate causes of mast fruiting [15, 16], although it has been contested whether they can constitute an ‘economy of scale’ [17].

For flower, fruit and seed production to be allocated efficiently over the life-time of the adult tree as masting years, a set of physiological resource-mechanisms, probably in serial form, must be in place [18]. Resource budget models, initiated by Isagi *et al.* [19], have simulated how a resource, usually carbon (C), accumulates in storage to an internal plant threshold which then triggers flowering. Almost all such models have involved the coupled wind pollination of neighbouring individuals [20–25]. They have often involved the Moran effect whereby masting is synchronized over large distances by driving climate factors. The triggering is thought to be brought about by a resource level being crossed in the plant which affects plant hormone concentrations that then cue a leafing axil to change to a flowering one, and finally enable fruit maturation [see e.g., 26]. In this respect, an individual tree can be modeled as an integrated and dynamic system which controls the timing and intensity of reproductive allocation to masting. Details of the involved physiological mechanisms can be expected to differ across tree species and sites, each achieving a stable yet flexible masting strategy in a different way.

The evidence for C as the main resource directly involved in masting is, however, quite weak. This conclusion comes from direct measurements on whole trees and observations of allocation and experimental manipulation of branches; and from overall C budgets and studies on the dynamics of non-structural carbohydrates (NSCs) [27–39]. A few studies have suggested that mineral nutrients may be the key resources underlying budget mechanisms, but the evidence remains scarce because the topic has been so little researched [40]. So far, Han *et al.* [41, 42] have shown that nitrogen (N) storage and mobilization is important for triggering flowering in *Fagus*. Ferandez-Martinez *et al.* [43] and Le Roncé *et al.* [44] have indicated that phosphorus (P) and zinc (Zn), and N and Zn, respectively are likely involved in mast fruiting of temperate trees more widely. Carbon should not be laid aside completely as a co-resource though. It is needed with N, P, potassium (K), and other macronutrients, not just to build fruits and seeds, but also indirectly to support growth and maintenance of organs such as fine roots and mycorrhizas which are essential for nutrient acquisition.

If macronutrients do underpin the mechanisms of mast fruiting, it could be expected that the element most important in controlling reproduction might be the one which is at the lowest level of availability at the site relative to the others, that is the one that is in most critical supply overall for the trees [15, 45]. This element will need to be accumulated to support successful reproduction, because seed filling depends on minimally attaining the necessary stoichiometric ratios for all elements required and doing so without undue costs to other tree growth and maintenance demands in the short to medium term. Further, extra-stoichiometric requirements may need to be met where the same element’s availability is very low in the soil: a rise in elemental concentration in the seeds would likely be advantageous for seedling establishment.

In this way natural selection would be expected to evolve a cueing-threshold-triggering system using the critical nutrient in the form of a cell storage product. Other elements in not quite so low supply as the first will also be important, which leads to the idea of nested systems of stores and thresholds. The search, then, is for those key limiting elements that most determine tree growth rates and nutrient cycling, which can also be shown to determine frequency and intensity of reproduction. Whilst the elements involved are probably different for different species, habitats and ecosystems, in principle, the same fundamental process overall can be postulated to operate.

Artificial selection of trees for fruiting under controlled nursery conditions has provided a major insight into tree physiology with regard to fruiting and what may lie as the basis to masting [46–49]. Cultivated trees often follow an ‘alternating bearing’ fruiting schedule, with years of ‘on’ and ‘off’ harvests over a regular 2-year period. Several temperate tree species in the Fagaceae show such an alternating pattern in natural forest, which leads to the idea that, theoretically, the minimal-intrinsic model across all masting species is perhaps alternating years of reproduction [11], but in naturally non-fertilized situations the interval between fruiting becomes extended beyond 1 year to allow time to accumulate critical resources and thus allow masting to effect maximize fitness. (‘Interval’ is here defined as the years of non-masting between masting events.) The fundamental question then becomes, how is the tree system coordinated to synchronize mast fruiting? Nutrient cycles have been strongly implicated in several fruit tree studies [e.g. 50, 51].

Lowering of minimum temperature has been suggested as a cue for mass, or ‘general’, flowering, and hence mast fruiting, in many species of Dipterocarpaceae dominating the aseasonal rain forests of S. E. Asian [52–57]. Much uncertainty remains, though, whether such drops in temperature are simply associated with the strong drought conditions that normally precede flowering events (low rainfall coming mostly with the El Nino Southern Oscillation) or they are acting as causal factors of floral induction per se. Dipterocarps are considered to have originated in the Indian subcontinent where the climate was seasonal [52]. In Thailand with a distinctly seasonal climate today, dipterocarp forests show annual phenological patterns which correspond to drought and lowered minimum temperature [58]. For Africa, eight principal tree species at Lopé, Gabon, had fruiting abundances which were strongly negatively correlated with mean minimum temperature in the main dry season (July-August) [59]. At Kibale, Uganda, community-level fruiting was also found to be negatively correlated with minimum temperature in the previous dry season [60].

The main hypothesis of the present paper is that limiting mineral nutrients can determine mast fruiting at the mechanistic, tree-physiological and proximate/ultimate level, particularly for those species that are ectomycorrhizal. This limitation is expected to be strongest where soil mineral nutrients at a site are in critically low supply and ectomycorrhizas are essential to nutrient uptake. Both the tropical Fabaceae subfamily Detarioideae and the Dipterocarpaceae are often ectomycorrhizal [61–63] and both commonly do show mast fruiting. Among the temperate families (Fagaceae and Pinaceae especially) there is also a strong association between masting and having this symbiotic mode of nutrient uptake [64]. Apart from the review of Corrales *et al.* [65], the potential role of ectomycorrhizas in masting appears to have largely gone uncommented [e.g. 8].

Two associated processes could potentially underpin how mast fruiting might be controlled by mineral nutrient limitation. They are not fully exclusive to one another. (i) Ectomycorrhizas might achieve a form of economy of scale due to their putatively extensive shared hyphal networks between the roots of large adult trees [66], possibly also via root grafts in dense stands [67], which could equilibrate levels of the limiting nutrient resource [68], and thereby lead to trees reaching similar internal thresholds simultaneously. (ii) Year-to-year stochasticity in climate [7, 69], especially seasonally-related rainfall, might cause soil nutrient availability and uptake to increase in the interval years between masting events, thus enabling stores to increase across all trees synchronously. For that no economy of scale per se is necessary. Furthermore, these two processes might interact positively, entraining common years of masting because they both depend on ectomycorrhizas and rooting activity.

An ecosystem well suited for testing the nutrient resource limitation hypothesis would be high-stature primary forest, for which C is in abundant supply, on soils very low in one or two elements that are macronutrients. Such an opportunity was provided by our long-term work at Korup National Park (SW Cameroon) since the 1970’s. In testing the main mineral nutrient hypothesis, it is also necessary to simultaneously test the most likely alternative, which is that C limitation, driven by radiation levels, determines mast fruiting. This latter aim is supported by an analysis of fruit production in a nearby oil-palm plantation, and a new analysis of previously collected tree ring growth data in relation to masting. The role of minimum temperature cueing, or possibly controlling, phenology is re-evaluated. Supporting knowledge of tree growth, forest dynamics, nutrient cycling, soils and climate variation at Korup are laid out in the following Background section. This information is essential for fully understanding the framework of the hypotheses being tested and the importance of the two associated processes.

The present paper builds on the study of mast fruiting in *Microberlinia bisulcata* A. Chev. between 1989 and 2004 [11] and extends it to 2017 (29 years): it is an update. For the parts concerning nutrients, some previously published values (as cited) are brought in with the newly reported ones to allow a complete five-element masting interval analysis. In Methods and Materials, and in Results, old and new values are clearly distinguished from one another. In the Discussion section, a synthesis of the new updated results, an improved main hypothesis and schematic model are developed and integrated with a review of the literature. These form a strong basis for new predictions, tests and analyses of mast fruiting in the African detarioids.

## BACKGROUND

### Forest ecosystem

Korup National Park, in the South-West Region of Cameroon, is covered by primary rain forest of the Atlantic Coastal type [70], and lies within the Guinea-Congolean refugium [71]. Local and regional maps are found in Gartlan *et al.* [72] and Newbery *et al.* [73]: further details of the site are given in Newbery and Gartlan[74] and Newbery *et al.* [11, 75].

*Microberlinia bisulcata* dominates large patches or ‘groves’, 1–3 km across, in the primary rain forest of southern Korup. Trees ≥ 50 cm stem diameter composed 17.8% of the overall density and 32.6% of total basal area in 2005, and together with co-dominants *Tetraberlinia bisulcata* and *T. korupensis* correspondingly 46.5% and 61.3% [75, 76]. All three species are strongly ectomycorrhizal as are several other co-occurring tree species of the subfamily Detarioideae in the Fabaceae, [formerly Caesalpiniaceae-Leguminosae, 73]. The taxonomy follows the revision of LPWG [77] and de la Estrella *et al.* [78]. They are not N-fixing trees. *Microberlinia bisulcata* trees attain considerable stature in the main canopy, characterized by wide dome-shaped crowns, and very extensive laterally spreading buttress systems [75, 79]. The species is a leaf-exchanger and therefore semi-deciduous: new leaves at the start of the dry season (December to February) push off the old leaves, flowering follows within a month [11, 68]. Flowers are outcrossed mainly by small bees and pollen seems unlikely to be limiting: bees are highly abundant in the canopy during the dry season [68]. It remains untested, however, whether insect pollination could form an economy-of-scale factor: in this connection presumably the density of trees plays a crucial role. Into the wet season (March to October) pods mature from soft green to hard fibrous brown structures. Seeds are dispersed ballistically in August-September [80].

The Korup-Ndian area along with Douala at the coast is climatologically special for western Central Africa because it has one clear dry season each year as opposed to the two less-distinct ones found elsewhere in the region. Mean annual rainfall for 1984–2016 was 5159 mm (range 4023 to 6531 mm). The typical seasonal course in rainfall is shown in Newbery *et al.* [81], with almost none in January and February but reaching 900–1000 mm in August or September. Site elevation of the site in southern Korup National Park (KNP) is ∼120 m a.s.l. (range of 40 to 200 m from south to north inland), slightly varying between swampy stream-fed lower areas up to local slopes, in places strewn with quartzite rocks. The soils are very sandy and free-draining, acidic and very low in available P and K [72], meaning that here high-stature tropical forest is growing on one of the most nutrient-poor sites in tropical Africa. Within groves at Korup, soil organic matter is largely confined to the top 2 cm of the profile, forming a surface mat that is densely occupied by fine roots and ectomycorrhizal hyphae [73, 81]. Larger vertical roots can tap deeper water reserves. Nutrient cycling in the groves shows a ‘fast-forward’ modus: leaf litter decay is fast, there is no permanent leaf litter layer, and most cycling via the very efficient mat happens in the early wet season [82, 83]. At this time throughfall nutrients help to prime decomposition and uptake [84]. For the rest of the year, heavy rains leach the forest, and this results in a tight closed ecosystem in which ectomycorrhizas play a central role in P acquisition and uptake [81]. In addition, during each year’s dry season, Harmattan dust brings some P to the forest via aerial deposition. A critical field experiment showed that growth and recruitment of *M. bisulcata* is not P-limited [85], possibly because of its highly evolved adaptations to the low P site or another element like K was limiting.

Recruitment of *M. bisulcata* over at least the last six decades has been poor due to very low seedling and sapling survival [68, 86–89]. Seed production in mast years is copious [11] but this species’ seedlings are shade-intolerant [90, 91]. The groves demonstrate transient dominance: the stands now are predominantly made up of large to very large trees, with extremely few small ones present [75]. The dynamics is thought to be one of patches arising and then decaying, being replaced, *in situ* or *ex situ*, within a mosaic of species-richer forest [74, 75]. The species is classified as a long-lived light-demanding tree [see 88]. Part of the expected life-history strategy of long-lived and light responsive species such as *M. bisulcata* would be to delay size at the onset of maturity (SOM) in order that greater seed output later offsets the relatively low probability of persistence in the forest understorey. Two further coevolved traits are the ectomycorrhizal habit mentioned and mast fruiting. Ectomycorrhizas not only provide advantages in resource acquisition for juvenile and young adult trees but are also thought to be a means to store resources which enable mast fruiting in the older adult ones [11, 68]. A key sub-hypothesis has been that these links between adults synchronize the timing of masts by equilibrating nutrient sinks across groves [68, 81]. There is mounting evidence from other forests that ectomycorrhizas build intricate below-ground hyphal networks between individual trees [e.g. 92, 93], although the definite evidence for these networks providing a means of C or mineral nutrient transfer is inconsistent, at least for adults to seedlings [94]. However, in the context of mast fruiting of adults, the hypothesized transfers are between very large trees occurring in dense stands (the groves at Korup).

### Mast fruiting

Mast fruiting in *M. bisulcata* was first realized from early phenological recording in 1988 to 1995, with prominent seed crops in 1989, 1992 and 1995 yet barely any seed fall in the intervening years [68]. Flowering occurs to some extent in most years (a few fail completely); though not always does a heavy seed crop depend on heavy flowering [11]. The two *Tetraberlinia* species tend to mast in the same years as *M. bisulcata* but not both together [68, and unpublished data]. After a propitious start with a potential 3-year cycle for masting (i.e., a 2-year interval between mast years), this cycle turned out to be shorter between 1999 and 2005. Because masting was associated with peaks in mean daily radiation in the dry season prior to fruiting this suggested that C was at a premium, with some threshold needed to be crossed before masting occurred. Detailed nutrient cycling studies in plots dominated by *M. bisulcata* showed, for the first time, that leaf litter falling after masting in 1989 was low in P concentration compared with the dry season of the years before and after (1988, 1990) , and this matched a decline in available P in the soil, especially in the surface organic layer [81]. This last result suggested that P was being shunted into the seed crop and less was remaining to be shed in litter and, given the low soil P at the site and the presence of ectomycorrhizas, P was also likely limiting mast fruiting. Tree C and P levels might be operating a double-threshold control system and this led to the phosphorus and climate proposed ectomycorrhizal response (PACER) hypothesis [81].

The climate at Korup varies stochastically from year to year, particularly regarding the start, duration and intensity of the dry season. The dates defining the actual season each year are based on 30-day running rainfall total (*rft*) falling < 100 mm [11]. This season plus the first 6 weeks of the wet season very largely determine tree physiological processes for the whole year. Fine-tuned sensing of the environment to optimize resource gain and allocation within a life history strategy is central to survival. Under- or over-allocation at the wrong time could lead to large fitness losses, hence the presumed operation of thresholds that safely trigger phenological change. That C and P might be dual constraints within a broader nutrient-limitation hypothesis as an explanation of mast fruiting for many tree species requires a detailed budget to determine how much C and P (also N and K) are allocated to fruits and seeds in a main masting event. Censuses of seed and pod counts, along with their masses and nutrient amounts, were obtained for the 1995 masting [95], showing that *M. bisulcata* produced on average 26K seeds per tree or 83.5K seeds/ha. Seeds are small (0.64 g each), and with an average of 2.0 seeds per pod, total pod mass was 25 g (see images in the Supplementary Material of Norghauer and Newbery [80]; for the 2007 and 2010 events, 2.5 seeds per pod were estimated. Masting resulted in an investment of 1034 kg /ha dry mass that year, and compared with leaf litter production in previous non-mast years [82], was 52% of it by comparison, or in a mast year 74%, i.e. a substantial allocation of resources [95]. Seeds and pods together had N and P amounts 13 and 21% of that in annual leaf litter production, indicating an important use of P during masting. Unfortunately, C, P, N and K investments in flowers are unknown but if included they would increase the reproductive allocation estimates slightly. Flowers of *Microberlinia* are very small in mass compared with seeds and fruits. Masting in *M. bisulcata* is highly synchronous, and it is unusual for a spatially isolated tree to fruit outside of an event [11, 68].

Using a detailed 150-tree phenological study in the main P-plot at Korup of 1995–2000, and incorporating circumstantial reliable observations in the field from previous and later years up to 2004, enabled a better picture of how dry season variability might control mast fruiting in *M. bisulcata* [11]. From daily rainfall and radiation data (recorded locally at Bulu, Ndian; 1984–2017), a clearer model was developed. Masting was tending to occur when the prior dry season was *drier* than average (more intense, low mean daily rainfall), plus the dry season in the year prior was *wetter* than average (less intense, high mean daily rainfall). After 1993, radiation did not show the same trend as in 1989–1992, mainly because after 1995 it climbed steeply as part of an inferred 11-year solar cycle [11]. The relative change in radiation between successive years was important, and not the absolute levels. There was no clear evidence of mast fruiting depending on minimum temperature in the dry season with data until 2004 [11, 68, 96].

The argument developed was that the current year’s dry season dryness reflected also to some degree increased radiation, but the wetter dry season the year prior was possibly indicating better conditions promoting tree nutrient uptake at the most effective time. The detailed phenology data showed that masting was strongest on trees on upper slopes (i.e. sites with relatively drier surface soils in years with a ‘dry’ dry season), and that in the moderately ‘wet’ dry season of 2000, those trees that masted were also on the drier sites. This tentatively pointed to surface soil moisture in the dry season being a key determinant to what was happening. The intensity of the dry season is unlikely to have particularly affected the water balance of adult trees themselves because they would have had access to deeper water sources: after all, trees flushed, leaves and branches grew, and flowering took place, in the dry season. Taken together, the preliminary evidence pointed to processes in the organic mat regarding fine roots and ectomycorrhizas, coupled to nutrient transfers within the trees and ectomycorrhizal network [11, 64]. With the masting time series extended to 29 years –almost twice its previous length, plus information collated from several nutrient studies at the site, further tree phenology censuses, a more detailed statistical modelling of fruiting on rainfall and radiation, and the effects of an unexpected herbivore attack – this paper assesses the extent to which C, P and K, might be determining the masting cycle in *M. bisulcata* and, in extension, presents a more refined hypothesis and model of the phenomenon.

## METHODS AND MATERIALS

### Climate records

Climate data continued to be available from the PAMOL Bulu Station, Ndian [97]. The station lies 12 km to the SE of the study site. Daily radiation, rainfall and temperature were recorded since 1 Jan. 1988 until 27 May 2017. Radiation was estimated using a Gunn-Bellani (BG) radiometer, with volume evaporated per day (V, ml) convertible into radiation (R, W m^−2^). Two periods, however, lacked radiation data because of instrument failure in March –May 1993 and April 2009–May 2011. Improved new radiometer calibration equations were developed using several shorter series of pyrometer measurements and these applied to the complete time series to obtain mean monthly radiation (Electronic Supplementary Material [hereafter ESM], Appendix 1). The short break in radiometer readings had been previously linearly interpolated, but the radiometer data for the much longer break 2009–2011 required more refined statistical methods. These are outlined in the analysis in ESM, Appendix 1. Since radiometers are rarely still in use today, this step in methodology is original. It also meant that the standard BG instrument calibration curve [used in 11] could be replaced with the now much better site-specific parameters. The equations were applied to the whole series (1989–2017), with differences to the first half of the time series referenced in ESM, Appendix 1. World Climate data bases unfortunately proved inadequate for estimating these missing values due to the sparseness of regional stations in western Central Africa and the special topography around Mt. Cameroon (SSE of Korup), with the few data records available for the last decades being incomplete (see ESM, Appendix 1).

A dry season occurred when the 30-day running rainfall total (*rft*) at Ndian (Korup) was < 100 mm, and this gave a defined start date and duration (the ‘drought-defined dry season’ of Newbery *et al.* [11]. This dry season typically moved one month back or forward around the modal months of December to February, coming earlier or later in different years, and often being shorter or longer. Hence, in what follows, ‘dry season 1991’ for example means December 1990 to February 1991 (modally). For a more general, less strict, annual matching of phenology to climate, a 12-month year starting on 1 December one year and ending on 30 November the next was adopted, with quarter 1 being December to February, and so on. Part-years were taken as November to March (5 months, dry) versus April to October (7 months, wet) to encompass the weeks before and after the dry season proper. Since its use in Newbery *et al.* [11], the definition of drought dry season was slightly relaxed here, in that within the outer defining limits a very few days where *rft* was at < 103 mm were included as drought days provided that the total returned to below 100 mm for the rest of the season. This affected four of the 16 years slightly in their dry season durations and mean daily rainfall values (Table 1).

**Table 1.**
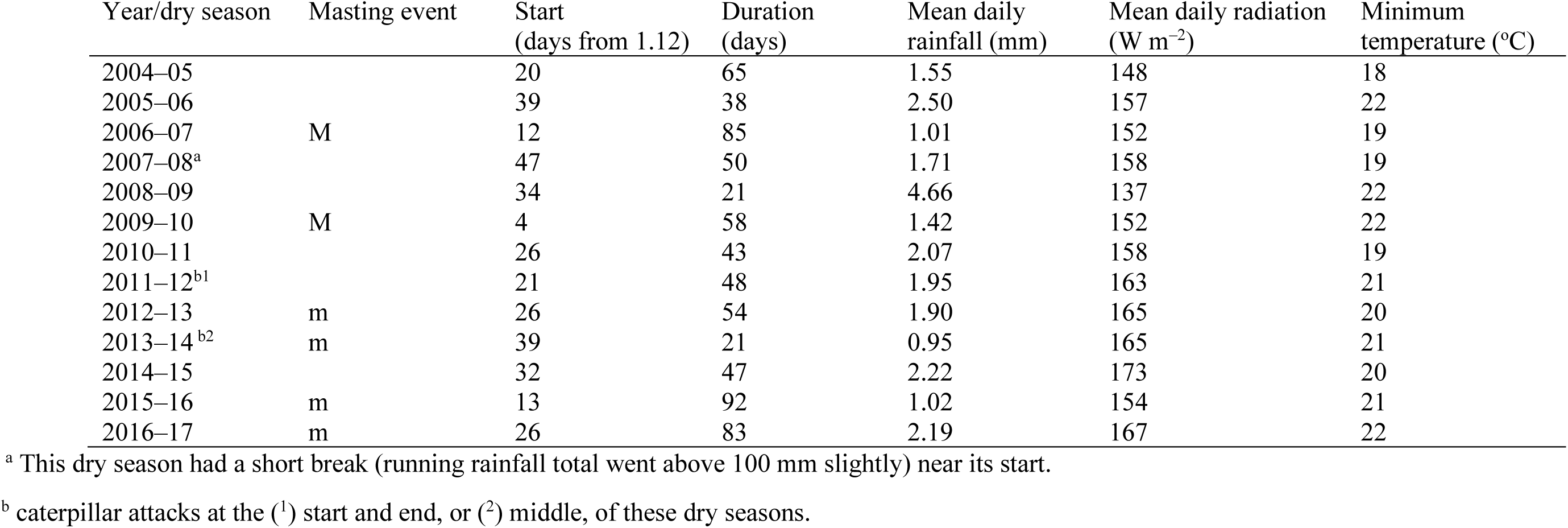
Incidence of mast fruiting in *Microberlinia bisulcata* at Korup for the years 2005 through 2017, and dry-season climate variables (i.e., last dry season before the event). ‘Year/dry season’ denotes, for example, the dry season from end 2004 into early 2005, coming before the masting in 2005. ‘M’ indicates a strong pronounced masting, ‘m’ a modest but still significant one. This table is a direct continuation of Table 1 in Newbery *et al.* [11] for 1988–89 to 2003–04. The exception to note is the updated radiometer calibration equation applied (ESM, Appendix 1) which changed the scale (linearly) for radiation. The new radiation values for 1989–2004 are also in ESM, Appendix 1.

| Year/dry season | Masting event | Start<br>(days from 1.12) | Duration<br>(days) | Mean daily<br>rainfall (mm) | Mean daily radiation<br>(W m<sup>-2</sup>) | Minimum<br>temperature (°C) |
| --- | --- | --- | --- | --- | --- | --- |
| 2004–05 |  | 20 | 65 | 1.55 | 148 | 18 |
| 2005–06 |  | 39 | 38 | 2.50 | 157 | 22 |
| 2006–07 | M | 12 | 85 | 1.01 | 152 | 19 |
| 2007–08<sup>a</sup> |  | 47 | 50 | 1.71 | 158 | 19 |
| 2008–09 |  | 34 | 21 | 4.66 | 137 | 22 |
| 2009–10 | M | 4 | 58 | 1.42 | 152 | 22 |
| 2010–11 |  | 26 | 43 | 2.07 | 158 | 19 |
| 2011–12<sup>b1</sup> |  | 21 | 48 | 1.95 | 163 | 21 |
| 2012–13 | m | 26 | 54 | 1.90 | 165 | 20 |
| 2013–14<sup>b2</sup> | m | 39 | 21 | 0.95 | 165 | 21 |
| 2014–15 |  | 32 | 47 | 2.22 | 173 | 20 |
| 2015–16 | m | 13 | 92 | 1.02 | 154 | 21 |
| 2016–17 | m | 26 | 83 | 2.19 | 167 | 22 |

<sup>a</sup> This dry season had a short break (running rainfall total went above 100 mm slightly) near its start.
<sup>b</sup> caterpillar attacks at the (<sup>1</sup>) start and end, or (<sup>2</sup>) middle, of these dry seasons.

Minimum temperature at heights of c. 15 and 35 m in the canopies of each of two large *M. bisulcata* trees in the P-plot were logged between 28 January and 16 February 2000 [96]. A highly significant linear relationship between mean canopy temperature (*minT_C_*) and the Bulu station temperature (*minT_B_*) was determined: minT_C_ = 4.8 + 0.8 · minT_B_, meaning that canopy temperatures were slightly higher (17.6 to 20.8) than at Bulu for the critical range of interest 16 to 20 deg. C. [96].

### Phenological recording

The main ‘P-plot’ is located in the southern part of the Park (5°10’ N, 8°70’ E; global coordinates 0476438/ 0553538) at an elevation of 96 m a.s.l.: refer to Newbery *et al.* [75, 76] for further details. Mast fruiting from 1989 to 2004 was reported and analysed in Newbery *et al.* [11]. Recordings were then continued from 2005 to 2017 for a subset of trees, and it is this full time series of 29 yr that is presented, with an extended analysis, in the paper here. Following pod maturity to completion was not possible in 2017 because of security restrictions preventing further access to KNP. The situation remains unresolved as of early 2025.

Evidence for masting rested on a combination of intensive phenological scoring of marked trees in two periods, quantitative trap-based estimates of seeding in two specific studies, from which masting fruiting is inferred; circumstantial records linked to seedling demography and seedling experiments in several years, and general yet objective assessments of the whole forest in other years during the course of large-plot census work, dendroband recording times, and seedling herbivore exclusion and nutrient lysimeter studies. (ESM, Appendix 2, table S1). Barely a year passed without adequate observation and reporting because of the continuing project workflow and frequent field presence of the Stirling-Bern research group since 1988.

The two periods of intensive recording were achieved in the following manner. To increase the frequency of recording per annum between 1995 and 2000, 150 *M. bisulcata* trees in the main 82.5 ha ‘P-plot’ [out of 294 in total with stem diameter ≥ 50 cm; 75] had been scored on 61 occasions (mostly at monthly intervals), for intensity of leaf fall, leaf flush, flowering and immature and mature pods: scale 0, none; 1, ∼25; 2, ∼50; and 3, ∼75 and up to 100% [11]. Later, within an 8-ha sub-area of the 82.5-ha P-plot (1000–1200 m E, and 100–500 N, with respect to SW origin; see Fig. 2a of [75], 61 of the 65 trees surviving since the last enumeration in 2005 (stem diameter, mean ± SE = 112 ± 4 cm, range = 54–174 cm), were scored between 2009 and 2013 in a similar manner as the first period. In 2014, fallen pod density on the ground was taken as a proxy for mature pod density (ESM, Appendix 2, table S1).

Masting was categorized as ‘high’ (labelled ‘M’) when all, or almost all (> ∼85%), of the recorded trees had many pods (scale = c. 3), and ‘moderate’ (‘m’) when fewer trees (∼45–85%) had many pods, or numbers of pods on relatively more trees were significantly lower. These masting events were clearly differentiated from those with ‘no’ or ‘very low’ pod production, and from those ‘low’ ones with just a few to several (∼ < 20%) trees masting (ESM, Appendix 2, table S1: the Evidence Table). The masting scale was therefore semi-parametric: except for the last 2 years the scale was quasi-binomial. In the Results section, three categories of event are use: ‘M’ and ‘m’ as masting; and the set “none, very low and low” as non-masting. Direct estimates of seed output, as a quantitative measure of mast fruiting, or even qualitative crown scoring, for all trees in all years was simply not feasible at this remote and difficult field location. The reduced and simplified scale for masting provides the best lowest denominator allowing all years of recording to be combined with very little bias.

Two canopy caterpillar outbreaks in 2011-12 and 2014 offered the opportunity to assess how loss of foliage to herbivory, and hence photosynthetic activity, could have interfered with tree reproduction.

### Nutrient sampling

#### Previous litter samples providing inter-mast nutrient data

In an earlier-conducted litterfall study between May 1990 and June 1992, monthly bulked samples from twenty 40-cm x 40-cm traps in each of 10 subplots had been divided into two halves: as set A, for separation into fractions (leaves, small wood, etc.), and set B, for separation of just leaves into the commonest 26 species [82]. From this set B, leaves of *M. bisulcata* were selected and later – as for the other species – they were analyzed for N, P, Ca, K and Mg concentrations. Leaflets and rachises were kept together. As this species is a leaf-exchanger, it has a very well-defined period of leaf litterfall each year, starting with onset of the dry season [11]. Five subplots were in low (LEM) and five in high (HEM) ectomycorrhizal stands (each 40 x 80 m) along the transect P in Korup [72, 81]. Since *M. bisulcata* occurred in only the HEM subplots, the number of replicate nutrient values per month was five. The 26 months had been divided into seasons according to the median rainfall pattern at Korup (as defined above), with those in the dry season being generally December 1990–February 1991 and December 1991–February 1992, and the other months the longer wet seasons. Only dry seasons are considered here, and nutrient concentrations for them are now averaged across the three constituent months. A mast fruiting occurred in (August-September) 1989, so dry season 1990–1991 is the second dry season after the event, and dry season 1991–1992 the third one. The next masting was in the wet season of 1992, so seen from this perspective the two dry seasons selected were ∼ 6 and 18 months prior to the second event.

In a later unpublished study of litterfall by Schwan [98] paired litter traps [same as those used by 82] were placed at random below the crowns of 20 *M. bisulcata* trees in the main P-plot. The bulk of the litter (∼90%) fell in the 3-week period of 9-30 Dec. 2002. This 2002–2003 dry season came immediately after a masting in 2002, and next one was in 2004 [11]. In this study [98], only leaflets were analyzed for N, P, K and Mg.

#### New litter samples included in the inter-mast data set

In the dry season 1997–1998 four traps of galvanized metal (same as used in 1990–1992) were compared with four similar ones but lined with hessian sacking (used in the 1989–1991 study of Newbery *et al.* [81], each set being placed at random in each of the two quarter plots of six HEM half-plots. Litter was collected over four consecutive fortnights between 21 Dec. 1997 to 15 Feb. 1998. *Microberlinia bisulcata* leaflets and rachises were again sampled separately, dried and analyzed for N and P concentrations. Not all traps had sufficient material and the replication per subplot/trap type/date ranged from 2 to 4. The N and P values were successively averaged for trap type, then subplot, and their weighted 4:1 leaflet-rachis averages found for the twenty-four date x plot combinations. Trap type had no effect on these two elements’ litter concentrations (initially to test for Zn contamination). This litterfall was collected midway between a moderate double masting in wet seasons 1997 and 1998. Chemical analysis followed [82].

An opportunity to sample *M. bisulcata* leaf litter immediately after a later mast fruiting came in the 2010–2011 dry season. A large system of over 400 seed traps had been used to follow seed production and dispersal in the August-September 2010 masting [80]. These traps remained in place until early 2011, collecting leaf litter. The dry season started on 26 Dec. 2010, and samples were taken on 29 Jan. 2011. This interval of ∼1 month would have covered a very large proportion of the leaf fall of this species that year, although the thin tail into February was not included. Thirty traps, from 68 small clusters of the traps within 25 ha of the 82.5-ha permanent plot [75], had sufficient quantities of almost-only *M. bisulcata* leaves, lying in the traps as a well-defined readily collectable upper layer. Leaflets and rachises were separated on this occasion and analyzed for the same five elements as in 1990–1992, later recombining at the whole leaf level. The mean dry mass ratio of leaflets to rachis in *M. bisulcata* leaves was 0.793:0.207, very close to a 4-to-1 ratio.

#### Nutrients in seeds and pods: extending the earlier data set

The 1998 masting event’s seeds and pods, previously analyzed and reported in [95] for N and P, were analyzed further for K, Mg and Ca using standard acid digest methods and spectrometric determination [see e.g., 99]. To find the nutrient masses allocated to reproduction (fruits) and to compare them to the same used in foliage, leaf litterfall values from another similarly strong (1992) mast year were taken from Green and Newbery [95]. In these new calculations, mature leaf concentrations (for N and P) were used from a retranslocation study in Chuyong et al. [82], which involved three dates: viz. December 1991, and April and July 1992. Litter fall dry mass very slightly underestimated mature leaf mass because of element retranslocation during senescence. Essentially, these leaf concentrations were from trees sampled on average 5 months before seed dispersal, and therefore concurrent with pod maturity that year.

### Soil moisture measurement

Soil moisture was recorded across two dry seasons to demonstrate that its variation could underpin soil nutrient processes which were affecting mast fruiting. *Microberlinia bisulcata* trees spread their extensive lateral fine root systems and ectomycorrhizas mainly in that drying soil surface layer [79, 81]. An additional aim was to determine by how much soil moisture content was reduced by rainfall deficit.

On 2 Dec. 2003, 12 soil moisture content sensors (Theta Probes ML2, Delta-T Devices, Cambridge, UK) were installed in a stratified random manner across the eastern 30 ha of the P-plot, one per block of 100 m x 250 m. Each probe was inserted horizontally (at 40 cm depth, excepting three at 20 cm due to access problems) into a hole excavated at the side of a small pit, the latter backfilled and packed down to the original density. A subsample of soil from the excavated side hole was retained to measure its fresh, water-saturated, and later oven-dry weight (105 °C). With a hand-held meter, soil moisture content (SMC, %) was recorded fortnightly until 26 Apr. 2005 (38 times). Readings were repeated between 15 Nov. 2006 and 15 Mar. 2007 (nine times), with one last recording on 29 Jan. 2011. In the years 2004 and 2007 *M. bisulcata* had major mast fruiting, in 2005 and 2011 it had none.

### Statistical analysis

The fitting of masting, as a 0/1 binomial response variable, to the climate variables was first limited to the 24-year period 1989–2012 because all the events were strong and pronounced (‘M’ in Table 1; ESM, Appendix 2, table S1). Logistic regressions were run with dispersion (φ) either fixed at 1.0 or estimated [100]. Whilst the latter gave marginally more significant fits, the former was adopted because the sample size was small and the estimation of φ would be unreliable [101]. The binomial error distribution appeared to be satisfactory: the complementary log-log one did not improve fitting. Simple logistic regression does not account for temporal autocorrelation, however. With records for only *∼*60 trees in common across all events, the sample size was too small to apply Monte Carlo subsampling in order to achieve temporal independence. Because of the non-stationarity, randomization tests will not provide valid inferences [102].

Extending the analysis to the full 29 years (1989–2017) required the ordinal, or proportional-odds, logistic model [100, 103, 104], where ‘m’ (moderate masting) was scored as 0.5 (see ESM, Appendix 2, table S1). Binomial and ordinal logistic models were run with the ‘glm’ and ‘polr’ commands in R [105, 106], using the package ‘MASS’ [107], and in GenStat for comparison where also φ could be estimated [108]. Given the modest length of the time series and data set, different models were fitted mostly with single independent terms; and when with two then the same variable with a 1-year lag, and only additively. No attempt was made to search for a best-fit subset model with interactions; if fact, most such model fits failed to converge when attempted. Because mean daily dry season rainfall and radiation were not correlated, these variables were fitted in separate models (see Results).

Times-series logistic regression – in full, generalized logistic autoregressive moving-average regression, with the command ‘glarma’ [R package glarma, 109] – accounts for temporal autocorrelations in binomial data [102, 110–114] and it was applied to the masting data.

## RESULTS

### Mast fruiting pattern

The extended time series shows that after 1995 there was a double year 1997–1998 with high seed crops, 1998 somewhat weaker than 1997, and then the interval dropped to 1 year (from a 3- to a 2-year cycle), with a masting sequence of 2000, 2002, and 2004, returning to strong ‘M’ masts in 2007 and 2010 with a 2-year interval, after which the control seemingly weakened to result in two pairs of consecutive years, 2013–2014 and 2016–2017 which had moderate ‘m’ mast fruiting (Fig. 1a). The time series as it extended began to show increasing non-stationarity (i.e. change in temporal pattern over time), the 3-year cycling of first period of 1989–1995 returned only once in 2004–2010: the regular cycle was seemingly temporarily disturbed. Nevertheless, ‘on average’ over the 29 years, and taking as markers the first of those pairs of years with partial masting (2013, 2016), there appears to be good evidence for an intrinsic 3-year cycle. Synchrony across trees within masting events was very high.

**Fig. 1.**
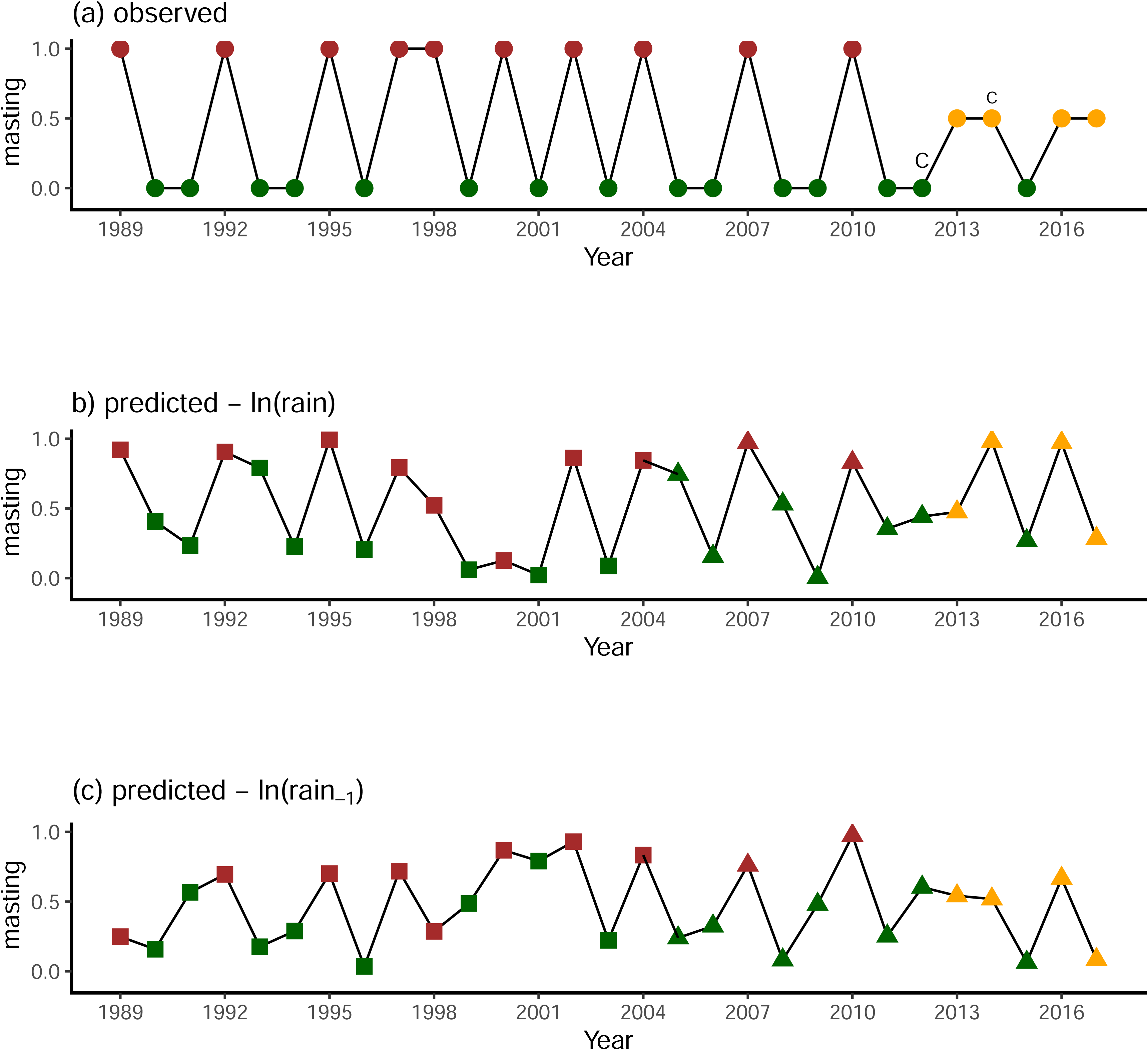
The time series of mast fruiting of *Microberlinia bisulcata* at Korup, 1989-2017. (a) The observed data. Symbol colours: brown, full; orange, partial; and green, no; masting event recorded. ‘C’ marks the year 2012, in the dry season of which the strong caterpillar attack occurred, and ‘c’ the one in 2014 which was a smaller more localized attack. (b, c) The probability of masting as fitted for 1989-2004 (squares), and predicted from 2005-2017 (triangles), using binomial logistic regressions of masting on the logarithm of rainfall in respectively the current and prior year’s dry season. Symbol colours: brown, full; orange, partial; and green, no; masting event recorded.

**Fig. 2.**
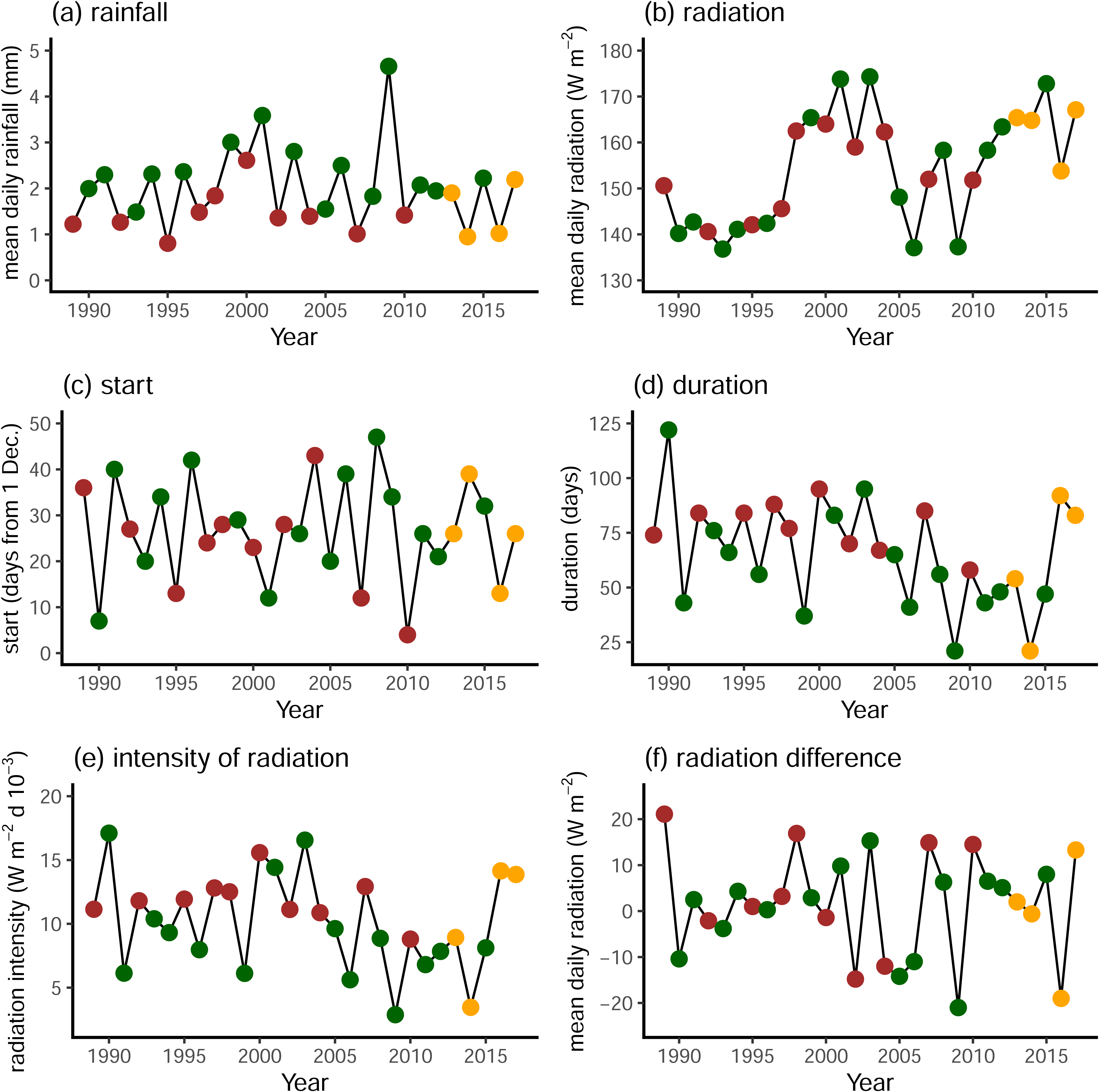
Times series of the dry season climatic variables associated with the masting series. (a) mean daily rainfall; (b) mean daily radiation; (c) start date (from 1 December of year prior) and (d) duration of the season, i.e. that which preceded the masting event; (e) radiation intensity (= radiation x duration·10^−3^); and (f) the difference in radiation (*radi_d_*) between successive years (current minus prior). Symbol colours: brown, full; orange, partial; and green, no; masting event.

### Climate variation in time

Of the four basic climate variables, mean rainfall and radiation per day in the dry season along with the start and length of the season (current year of masting and year prior), only the first required ln-transformation to remove positive skew along the independent variable axis. Dry season rainfall and radiation were not significantly correlated (*r* = 0.110, *P* = 0.57, df = 27) and could be treated as being independent.

Mean daily dry-season rainfall in years of full masting was lower than in those years’ prior with no masting (Fig. 2a) except for 2000. Further, if the averages of the rainfall values for the two double partial masting years are taken these are on a par with the masting ones. The time series for rainfall showed weak temporal autocorrelation (ESM, Appendix 3, fig. S3) and likewise, albeit a little more strongly, for ln(*rain*) at lag 1 (–0.293 vs. –0.221). (Note that the first correlation differs slightly from the rain versus rain_–1_ one based on data which included the lagged 1988-value.) Radiation over time showed a distinctive pattern with a clear central peak between 1996 and 2005, then a rise again, apart from 2009, to those higher levels (Fig. 2b): accordingly, the series evidenced non-stationarity, and as a consequence autocorrelation was positive and significant for lags of 1 and 2 years, then significantly negative (*P* < 0.05) for lags of 6–8 years (ESM, Appendix 3, fig. S3). The partial autocorrelation remained significant for a lag of 1 year only, however.

Start of the dry season varied around the third to fourth week of December with no clear patterns or differences between masting and non-masting years (Fig. 2c), but it was negatively autocorrelated with a lag of 1 year, i.e. early starts tended to be followed by late ones, then early again, in successive years (ESM, Appendix 3, fig. S3). Excepting the two last years, duration of the dry season revealed a general decline (an approximate halving), those years with masting being often longer, and conversely of the six shortest-season years there was no masting, apart from 2014 which was partial (Fig. 2d). Interestingly, autocorrelation of duration was almost significant (*P* < 0.05) at a lag of 3 years (ESM, Appendix 3, fig. S3).

Radiation intensity showed a very similar pattern to duration (since radiation varied much less than duration) (Fig. 2e; and ESM, Appendix 3, fig. S3). Intensity of radiation in the dry season was expressed as mean daily radiation x duration. Lastly, difference in radiation (*radi_d_*) between successive years (one approach to removing trend and non-stationarity) was mixed regarding masting response, although again a weakly significant negative autocorrelation for a lag of 1 year was evident (Fig. 2f; and ESM, Appendix S3, fig. 3).

**Fig. 3.**
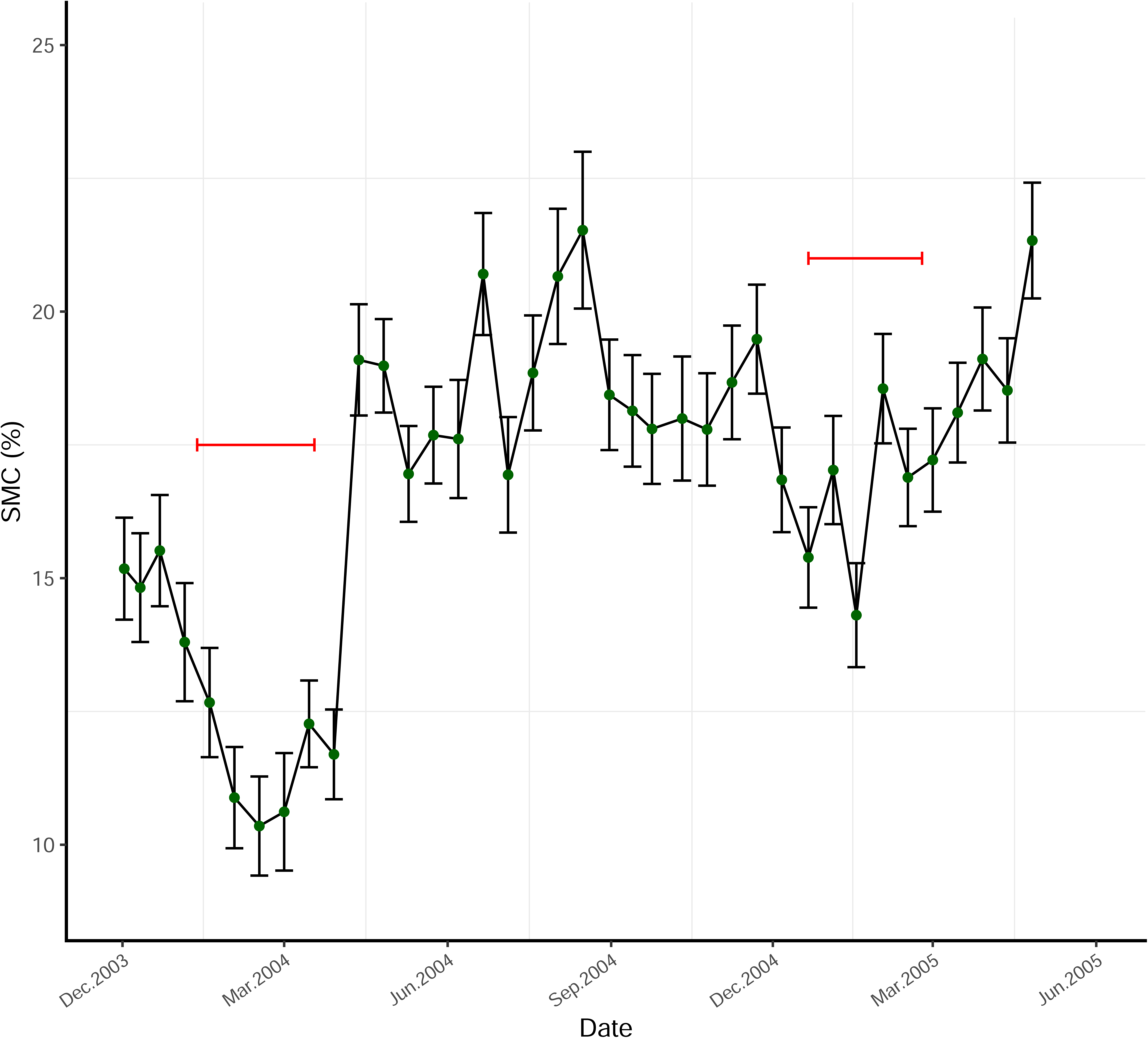
The course of soil moisture content (SMC, %), shown as means and SEs, at (mostly) 40 cm depth, for 2003-2005 at Korup. The horizontal red lines indicate the span of the dry season for each year (see also ESM, Appendix 5, fig. S1).

### Modelling masting and climate

Binomial logistic regressions for 1989–2012 showed a significant negative effect of mean daily rainfall in the current year’s dry season (*P* < 0.05), and a positive effect of mean daily rainfall in the dry season prior (*P* < 0.05), on masting. Using ln(*rain*) and ln(*rain_–1_*) improved the regressions fits with *P* < 0.015 (ESM, Appendix 4, table S1). Mean daily rainfall was only weakly correlated with rainfall the year prior: untransformed values (*r* = – 0.190, *P* = 0.374), transformed (*r* = –0.272, *P* = 0.15). This last result is important because a strong negative temporal autocorrelation, i.e., a year-to-year increase-decrease reversal, was absent. Regressions with both rainfall terms led to reduced fitting because these coefficients were now partial ones adjusting for the correlation (*P* = 0.093 and 0.055). The same regressions with ln-transformed variables also changed the significance levels very little (*P* < 0.10; ESM, Appendix 4, table S1). Ordinal logistic regressions improved the fitting further with the full 1989–2017 data. Concentrating on the models using ln-transformed variables, separate regressions were now more, and highly, significant (*P* < 0.01), having coefficients with the same signs as for the binomial logistic ones (ESM, Appendix 4, table S1) and, whilst significance decreased again using two-term models, the fits were still relatively strongly significant overall (*P* < 0.025).

The mean difference in ln-transformed dry-season mean daily rainfall, ln(*rain*), between the current masting (t_0_) and year prior (t_–1_) was highly significant (*P* < 0.005), but much weaker for the same and penultimate year (t_–2_) (*P* = 0.15). Back-transformed means were 1.37, 2.49 and 1.69 mm/day respectively (see ESM, Appendix 4, table S2). Comparing rain at t_0_ with means of *rain* at t_–1_ and t_–2_ – applying equal or unequal weighting – led to very similar results as those at t_–1_ alone (*P* < 0.005). As a robustness test of the strong vs moderate (‘M’/’m’)-grading of mast fruiting, and how that might have affected the relationship with mean daily rainfall in the dry season where there was some uncertainty in the final grading, the 1989–2017 ordinal logistic regressions were rerun with ‘M’ for 1998 and 2000 replaced by ‘m’, i.e. 1.0 by 0.5. The dependence of masting on ln(*rain*) in the current and prior seasons became slightly stronger (*t* = –3.209, *P* = 0.0036; *t* = 2.913, *P* = 0.0074, respectively).

Mast fruiting showed no significant dependence on mean daily radiation in the drought-defined dry season, for either the 1989–2012, or 1989–2017, time series (using binomial and ordinal regression, respectively), for the current (*radi*; *P* = 0.72 and 0.80) or prior (*radi_–1_*; *P* = 0.55 and 0.62) seasons. Correspondingly, start date was also a poor predictor of masting for the current (*start*; *P* = 0.34 and 0.29) or prior (*start_–1_*; *P* = 0.18 and 0.27) seasons. Duration, although only weakly significant, was better for the current (*dur*; *P* = 0.080 and 0.095) or prior (*dur_–1_*; *P* = 0.071 and 0.138) seasons. It is important to note here that radiation values for the 2010 and 2011 dry seasons had been interpolated (ESM, Appendix 1), so model predictions for the 2010 masting event come with an additional level of uncertainty. Masting was only marginally or non-significantly dependent on masting intensity in the current and previous dry seasons (positively and negatively respectively; binomial logistic, *P* = 0.083 and 0.097; ordinal logistic, *P* = 0.096 and 0.165). Hence there was some indication that high intensity reduces the likelihood of masting in the following year. Differences in mean daily radiation (current-minus-previous, *radi_d_*), on the drought-defined dry-season basis, had no clear effect on masting (binomial logistic, *P* = 0.31; ordinal logistic *P* = 0.84).

Radiation did show a long-term trend from the start of recording until 2017 (ESM, Appendix 3, fig. S2), with evidence of the 11-year solar cycle. This was not related to the change in instrumentation at Bulu (ESM, Appendix 1). Using a LOWESS regression fitting model in R (‘lowess’ command – a set of local smoothing splines), this trend could be removed and the differences between empirical measurements and the detrending curve then used as adjusted local year-to-year changes (ESM, Appendix 3, fig. S2[a]). The best result was with a span parameter of *f* = 0.25, which meant that radiation values > 1 year prior or ahead had some weight in fitting values for any one year, which simple differences between successive years did not. Binomial (1989–2012) and ordinal (1989–2017) logistic regression fits of masting to drought-defined dry-season *radi* and *radi_–1_* were, however, all clearly insignificant (*P* = 0.34 to 1.00).

Lastly, to check whether heavy showers early in the dry season may have damaged flowers or interfered with pollination success, mean daily rainfall was plotted across each year’s season. Of the 14 years with strong (‘M’) or moderate (’m’) mast fruiting, ten had no showers of ≥ 15 mm/day in the first 30 days, whilst in the 15 years without mast fruiting eight had no showers (χ^2^ = 3.55, df = 1, *P* = 0.060), suggesting only a marginal tendency for mast years not to have experienced potentially destructive rainfall events.

Taken together, these regression model fits suggested that only mean daily rainfall in the current and prior dry seasons was strongly affecting the odds of masting. The exponent of the coefficient (e^β^) estimates the fold-change in odds of masting per unit increase in mean daily rainfall. Since the range in mean daily rainfall over the years was 1 to 3 mm, this translates on the ln-scale to *c*. 1 unit, meaning that in the transformed-variable models the change in odds covers the range in rainfall. This outcome provided a sound basis for predicting future masting events.

### Predicting masting from models

Using binomial logistic regressions of masting 1989–2004 on ln(*rain*) in the current and previous years, fitted models were used to predict masting in 2005–2017 (Fig. 1), on the original response scale of 0 to 1 [π; 100]. The series started at 1989, not 1987 as used in Newbery *et al.* [11] and autocorrelation was not incorporated (see ‘Time series analysis’ later though). The model fit for ln(*rain*) was good for the mast years 1989, 1992, 1995 and 1997 (marginal for 1998), and good for 2002 and 2004, but it wrongly indicated a mast in 1993 and missed the mast in 2000 (Fig. 1b; fitted intercept [α] = 3.61 ± 1.80 and slope [β] = –5.77 ± 2.62, *t* = –2.206, *P* = 0.027). Predictions of masting in 2007 and 2010 were very high (π = 0.97 and 0.83 respectively), and that it did not occur in the intervening years. After 2012 though, one each of the pairs of moderate (‘m’) masts was predicted as full (‘M’, π = 0.98 and 0.97). The predicted masting in 2005 (π = 0.75) was incorrect, however.

With the model fit for ln(*rain_–1_*), 1992, 1995, 1997, 2000 [cf. with ln(*rain*)], 2002 and 2004 were all well-fitting as mast years, but 1989 and 1998 were not: 2001 was fitted as a mast year wrongly (Fig. 1c; fitted α = –2.47 ± 1.40 and β = 3.96 ± 2.00, *t* = –1.98, *P* = 0.027). Again, the model predicted exactly the mast years 2007 and 2010 masts (π = 0.76 and 0.97 respectively), and it managed three of the four ‘m’ (moderate) years afterwards as being moderate masts correctly. Therefore, similar to the outcome for cycles and intervals in mast fruiting, the model fits were fair-to-good up to 2012 (the caterpillar-outbreak year; see later subsection) but after that disorder and unpredictability ensued. Binomial logistic regression fits made with the 1989–2012 data gave predictions (not shown) that were very similar to those for 1989–2004, in the main because the masting in 2007 and 2010 matched so well the time series to 2004: yearly π-values (fitted and predicted) from the two runs were in very good agreement.

The median effective level of ln(*rain*), that is the level when π (x) = 0.5 [defined as x = –α/β; 100] was 0.624, which back-transformed to 1.869 mm/day – a value very close to the estimate for the ln(rain_–1_) estimate of 1.867 mm/day. This median level, which might be interpreted as a form of threshold for non-masting and masting, applied well to the events and their rainfall values in most years up to 2011 (Table 1, Fig. 2a), although ln(*rain*) and ln(*rain_–1_*) differed in a few years as being better or worse fitting and predicted. Years 2012, 2013 and 2014 had moderately high and quite similar mean rainfall in their dry seasons (1.90–2.07 mm/day; Table 1), so aside from the caterpillar intervention, the rain threshold theory would not have been expected to work too well – a form of counterfactual evidence. Afterwards, in 2014–2017, the return to low–high alternations in rainfall should, *ceteris paribus*, have led to ‘M’ (strong) masting in 2014 and 2017 (as predicted by the binomial logistic regression for ln(*rain*) (Fig. 1b), but it still did not achieve that outcome.

### Time series approach

Application here supported the binomial logistic regression results for 1987–2012 even more strongly (*P* < 0.01 and *P* < 0.05 for ln(*rain*) and ln(*rain_–1_*) respectively, for both lags p = 1 and 2, (ESM, Appendix 4, table S1). Regressions using the time-series logistic regression model for *radi*, *radi_–1_*, *start* and *start_–1_* either failed to converge, or the fits were insignificant (*P* = 0.17 to 0.73). Whilst there was a weak positive effect on masting for duration in the current year (β = 0.041 ± 0.025, *z* = 1.64, *P* = 0.10), an even stronger negative effect of duration was evidenced for the year prior (duration_–1_; β = –0.099 ± 0.038, *z* = –2.62, *P* = 0.0087) (ESM, Appendix 4, fig. S2). The last relationship analyzed before with a binomial logistic model was only marginally significant (*P* = 0.073), so the evidence for the effect improved.

The dependence of mast fruiting on dry season rainfall appears to be limited to the current and previous (t_–1_) years, because repeat runs replacing *rain_–1_* with *rain_–2_*, the mean daily rainfall in the second year prior to the current one (t_–2_), were all clearly insignificant (*P* = 0.39 and 0.40) for 1989–2012 binomial logistic regressions, rainfall ln-transformed, single and two-term models respectively; and they were correspondingly insignificant (*P* = 0.35 and 0.76) for the 1989–2017 series with ordinal logistic regression. Moreover, dry-season rainfall at t_–2_ was poorly correlated with the same in the current year (*r* = 0.027 and 0.110), but slightly better with that at t_–1_ (*r* = –0.187 and –0.257). Time-series logistic regression fits did not improve fitting to intensity of radiation (current or prior), differences in dry season radiation in successive years, or smoothed radiation regression residuals (current or prior) (*P* = 0.094, 0.062, 0.25, 0.74 and 0.26; respectively).

### Extending models for radiation

Whilst a rainfall deficit delimitation will define the drought period in terms of limited water availability, it may not necessarily correspond precisely to times of heightened radiation. Returning to the modal definition of dry season, namely the calendar months December through February (DJF), or first quarter of the phenologically-referenced year, mean daily radiation can likewise be found (ESM, Appendix 3, fig. S1). Masting was not strongly related using binomial, ordinal or time-series logistic regressions, to this quarter’s radiation however, in either the current or prior year (*P* ≥ 0.18; ESM, Appendix 4, table S3[a]), and also not for the other three quarters (ESM, Appendix 3, fig. S1; *P* > 0.4; model fits not shown). By contrast, and surprisingly, masting was strongly dependent on the *difference* in radiation, both in absolute (*radi_d_*) and percent change (*radi_d%_*) terms, from prior to the current year, and with all three regression models (*P* = 0.007 to 0.024; ESM, Appendix 4, table S3(a) and fig. S3; and ESM, Appendix 3, fig. S1). Using again LOWESS smoothing regressions to remove trend in radiation (ESM, Appendix 3, fig. S2[b]), with binomial logistic regression (1989–2012) masting weakly positively depended on differences in *radi* (*P* = 0.051) and negatively, though more marginally, for *radi_–1_* (*P* < 0.1) (ESM, Appendix 4, fig. S3 and table S3[b]). Time-series logistic regressions improved the fits especially for *radi* (*P* < 0.02), although only slightly for *radi_–1_* (*P* = 0.08). Fits to detrended differences (1989–2017) with ordinal logistic regression were intermediate in significance to those with binomial and time-series logistic ones (ESM, Appendix 4, table S3[b]). Overall, there emerged moderate to strong evidence that the larger the increase in radiation between successive years, or the more positive the difference from the trend, the more likely a mast fruiting. This difference effect was acting as a booster to fruiting. It was not obvious in the earlier analyses of Newbery *et al.* [11].

An interesting feature of the LOWESS fits (ESM, Appendix 3, fig. S2), when taking radiation according to the two definitions of dry season, is how the outliers changed in relation to the smoothed curves. In the drought-defined case, 2001 and 2003 (both 1 year before masting) were very high, and 2006 and 2009 (also both 1 year before masting) were very low. Changing to the DJF-defined case brought 2001 and 2009 radiation much closer to the curve, and the binomial and time-series regression fits were improvements. Further, the autocorrelation was notably stronger, positive and significant, at lags of 1 and 2 years for this DJF-defined dry season than the drought-defined one, shifting to a significant negative correlation at an 8-year lag (ESM, Appendix 3, fig. S4). The partial correlation, again, remained significant for just the lag of 1 year. In the third (JJA) and fourth (SON) quarters a similar but slightly weaker pattern for the function occurred, the negative lag now at 10–12 years (ESM, Appendix 3, fig. S4). More interesting, however, was the almost unstructured weak pattern of the function for the second (MAM) quarter, which means no temporal autocorrelation over the early wet seasons (ESM, Appendix 3, fig. S4). Difference in radiation was significant for a lag of 3 years, and remained so for the partial correlation, hinting at the 3-year cycle.

### Start and duration of the dry season

To shed some light on the apparent paradox of the different definitions of dry season determining the relationship between masting and radiation, regression models were rerun with an extended dry season of the months of November through to March. All binomial logistic model fits, 1989–2012, using radiation in the current and prior dry seasons on this new basis were clearly insignificant (*P* = 0.27 and 0.69). However, dependence on masting on *radi_d_* was marginally significant at *P* = 0.071 (*radi_d%_*, *P* = 0.052), and correspondingly so was it for the time-series logistic model with p = 2 (*P* = 0.056 and 0.043). Compared with the fits for the models using DJF as dry season (ESM, Appendix 4, table S2[a]), these results for the 5-month season were weaker, suggesting that the influence of radiation was being diluted by inclusion of the additional 2 months; and this was reflecting the early wet season autocorrelation. Comparing radiation defined by DJF and that defined by the drought dry season graphically, both for differences between successive years and as residuals from the smoother curve over time (ESM, Appendix 4, fig. S4), indicated that a majority of the years lying above the 1:1 reference line were ones with masting and most of those below were non-masting. This confirmed that raised radiation differences and deviations were associated with mast fruiting.

The placing of start and end dates of the dry season was critical to the analysis outcome because the degree of overlap between the two types of dry season were very variable from year to year (Table 1, Fig. 1a). Mean, standard deviations and ranges in absolute differences were larger and wider for the drought- than DJF-defined dry periods (1.41 ± 11.14 [–21.0 - 21.1] and 0.72 ± 9.62 [–15.0 - 16.2] respectively). Even so, the reasons for the considerable overlap could be various because start and end dates of the drought-defined dry season were often very different from those of the DJF one.

When the mean daily radiation of the non-overlapping weeks with DJF increased or decreased, or vice versa, the *radi_d%_* altered between successive seasons. Since masting fitted to *radi_d_* much better for the DJF- than drought-defined seasons, this would suggest that (a) radiation levels outside of the DJF months were most likely important, and (b) DJF-defined seasons allowed a more consistent year-to-year comparison than the drought-defined ones. That rain may have been lacking, which defined the duration of the drought-defined season, did not also imply radiation was changing proportionally. Radiation could have been sometimes high in the day and rain fell at night: conversely radiation could have been low when there was cloud cover and yet no rain, as was the case in some years at the transition to the wet season. Differences in successive years’ dry season radiation were more, respectively less, pronounced in the first and second half of the time series, for the DJF- than the drought-defined dry seasons.

### Soil moisture content in dry season

On graphing the time series for 2003–2005, three of the 12 sensors had three anomalous and coinciding high peaks (∼ 40 - <60%) in the 2004 and 2005 early-to-mid wet seasons suggesting that their locations were prone to flooding. These sensors’ data for the whole period were accordingly omitted. Two of them were destroyed later by hunters. The time series of mean SMC is shown in Fig. 3: it included the two dry seasons of 2004 and 2005, defined as before as when *rft* fell < 100 mm. Mean SMC over the 1.4 years was 16.80 ± 0.48%. Repeated measures analysis of variance for six times within each of the two dry seasons (Fig. 3), using all 12 sensors’ values, indicated that differences in mean SMC at the 20-and 40-cm depths were very small and insignificant: 10.98 vs 10.69% in 2003–2004 and 16.20 vs 15.89% in 2004–2005, *P* = 0.88 and 0.89 respectively, df = 1, 10). In a similar-length period in the late wet season 2004, starting soon after the peaked occurrences that year, a small difference was evident but again it was clearly not significant (20.20 vs 18.13%, df = 1, 10; *P* = 0.39). Interactions between depth and time were all insignificant for all three short periods (*P* = 0.19 to 0.36).

After the strong dry season of 2003–2004, during which SMC reached a minimum of 10.4%, its values remained constant at 18.24 ± 0.31%. The dry season of 2004–2005 had much less influence on SMC (Fig. 3). The lowest SMC values in 2004 coincided with the lowest *rft* values (down to 12 mm then 0 mm over 4 weeks); in 2005 the SMC either did not go so low or remain low for long, and the match with SMC was weaker. At the start of the 2003–2005 series, the means (± SE) of SMC recorded and those found by gravimetric analysis were 15.09 ± 0.52 and 15.50 ± 0.82% respectively. The estimates were positively correlated (*r* = 0.677, df = 10, *P* = 0.016). Saturated water holding capacity was on average 38.15 ± 1.16%, less than the SMC at the peak times, confirming that some locations were not always free draining.

Between 2 Dec. 2003 and 26 Apr. 2005, SMC was related to 30-day running rainfall total as follows: SMC = 14.4 + 0.0070·rft (*R^2^*_adj_ = 40.5 %, *F*_1,36_ = 26.18, *P* < 0.001, *n* = 38), inferring 14.4 % at 0 mm, and 21.4% at 1000 mm – the last *rft*-value being close to the June 2004 maximum of 995 mm. The difference in SMC between the very driest and wettest times of the mast year was therefore only 7% – for those locations assumed free-draining. SMC before the dry season of 2006–2007 was 20.7 ± 0.10% and declined sharply and linearly to 14.0 ± 0.67% (*n* = 9 sensors), a difference of 6.7%. With the dry season that year lying within the span of SMC recording dates, this last value was likely very close, if not at, the 2007 SMC minimum. The mean value on 29 January 2011, a date also well into that year’s dry season was 16.2 ± 0.8% (*n* = 8, one sensor lost because its cable had been chewed), a value close to that in the dry season of 2005 (Fig. 3).

The 30-day running rainfall total, *rft*, used to define the dry season in terms of drought when this total goes below 100 mm, does not indicate either how much is the deficit, or how steep is the decline into, or incline out of, the dry season. ESM, Appendix 5, fig. S1 shows the course of *rft* as the daily rainfall events occurred. The intense dry season of 2003–2004 (2004 was a masting year) resulted of *rft* going to zero, or very close to zero, for about 6 weeks (ESM, Appendix 5, fig. S1[b]), while in 2004–2005 (2005 was a non-mast year) it was near zero for perhaps just 2 weeks, and shorter overall (ESM, Appendix 5, fig. S1[c]). In 2002–2003, for comparison, the also long dry season *rft* reached zero for a couple of days, and whilst many days had no rain the season was broken by three high rainfall events keeping *rft* just below 100 mm (ESM, Appendix 5, fig. S1[a]). The small decrease in mean daily rainfall between 2005 and 2004 (Fig. 2) translated into a surprisingly large difference towards the lowered SMC and suggested a non-linear effect, which means less rain led to proportionally much more soil drying when rainfall was low-to-average.

### Caterpillar outbreaks

In the latter half of December 2011, and again in the first half of March 2012, an outbreak of a black caterpillar was observed that fed on young foliage of *M. bisulcata* trees recorded for phenology in the P-plot. Although not quantitatively measured, the attacks were extensive, as evidenced by the substantial amounts of falling frass, numerous caterpillars hanging below the crowns on threads, and litterfall consisting of soft, green leaf tissues (G. A. Neba and S. Njibili, pers. obs.). The same caterpillar was also recorded at another *M. bisulcata* grove 4.2 km due south, in March 2012 [115]. However, no visit was made to that site in December 2011. Fallen caterpillars were also observed feeding on young leaves of *Oubanguia alata*, a common understorey tree growing below *M. bisulcata* crowns at Korup.

Also in the latter half of December, but in 2008, an outbreak of most likely the same caterpillar species (an *Achaea* sp., probably *A. catocaloides*) had been recorded on *T. korupensis* trees in the same P-plot [115]. At that time, feeding upon *M. bisulcata* foliage was limited to low levels on just those crowns closely adjacent to the more heavily attacked ones of *T. korupensis*. Furthermore, and separately, at the end of January 2014 a morphologically different and larger species of caterpillar, yellow-green and behaving very differently from the smaller black one, was seen on five of the 65 monitored *M. bisulcata* trees (ESM, Appendix 6, text and fig. S1). Most probably it was another *Achaea* species. Intriguingly, this caterpillar species was feeding on *M. bisulcata* flowers, not young foliage. How influential the new insect herbivore was for *M. bisulcata* phenology that year is uncertain because its distribution was localized to a 2–3-ha patch where the *M. bisulcata* basal area was high.

### Tree phenology and masting intensity

Tree phenology of *M. bisulcata* was compared between the heavy mast fruiting year of 2010, and the year of 2012 in which fruiting was very low (Fig. 4; ESM, Appendix 2, table S1). Both time series covered the drought-defined dry season of that year and followed changes in leaf-flush and leaf-fall (leaf exchange), and flowering, pod formation (green) and pod maturity (brown). Attaining continuous fortnightly scoring was not always feasible and each series has two gaps. Leaf flush began a little before the start of the time series recorded for 2010, i.e., earlier than December 2009 (Fig. 4a), but was adequately captured for 2012 (Fig. 4b). The bulk of the leaf-flush happened within each year’s dry season, flowering well within it in 2010 but coming at the end in 2012. Even though the sum of the flowering scores was similar in both years, pollination resulted in more green and brown pods in 2010 than 2012 (Fig. 4a, b). Caterpillars were not seen in the other years between 2009 and 2013, notably not in 2010. In 2010, green pods were apparent between 18 January and 3 July, which is for 5.5 months, with a 1-month overlap as they turned brown between 5 June and 28 August, over 2.5 months (Fig. 4a). A similar time course was recorded in 2004, and likewise (for green plus brown) in 1995 and 2000 [Fig. 4b of 11]. Interestingly the overall period of pod formation was much shorter in the two back-back years of masting 1997 and 1998.

**Fig. 4.**
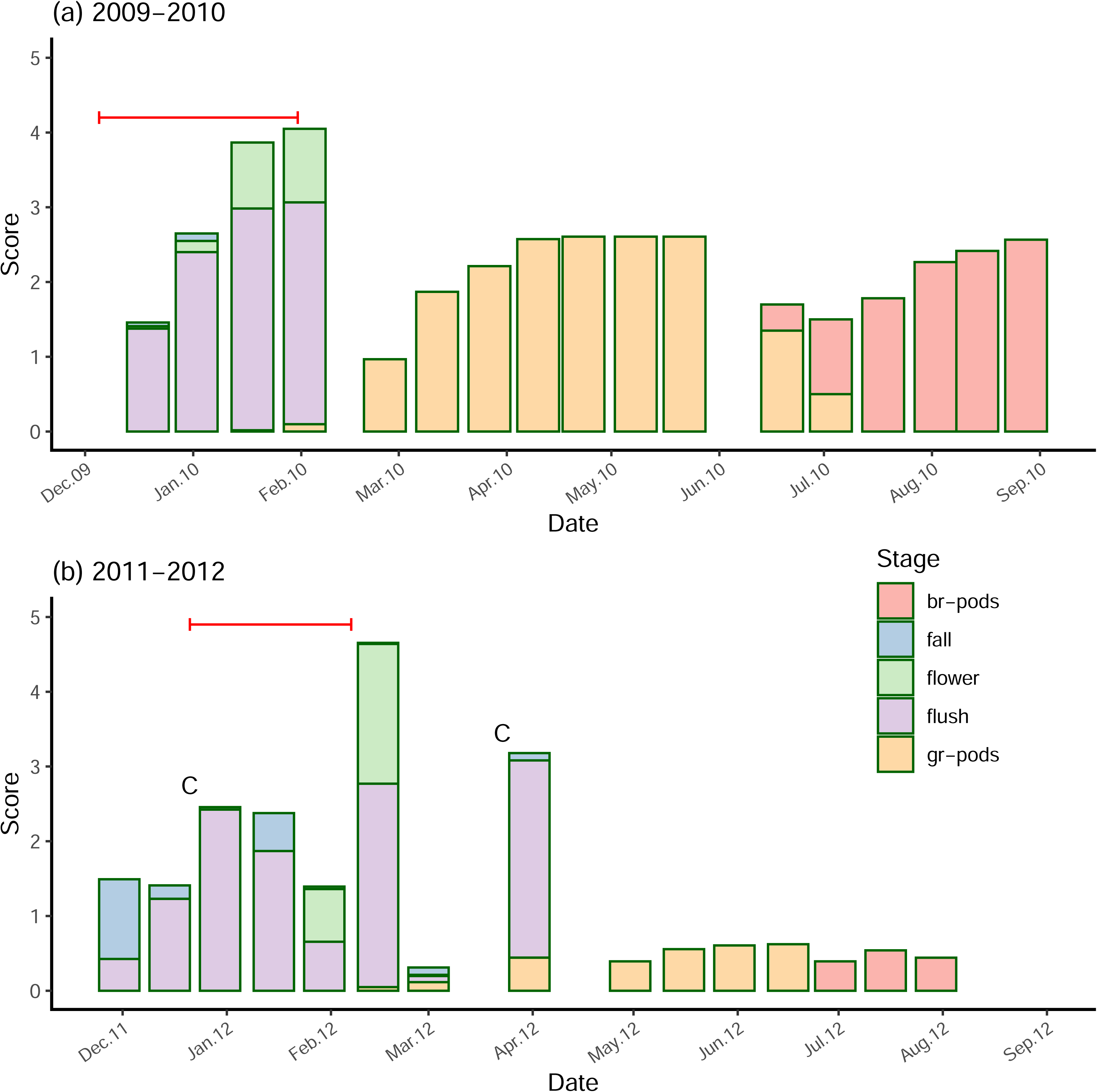
Changes in the mean scores of five phenological stages of *M. bisulcata* trees at Korup in (a) 2009-2010 and (b) 2011-2012. Stages: leaf fall (blue), leaf flush (purple), flowering (green); immature (yellow) and mature (red) pods. The horizontal red lines indicate the span of the dry season for each year (see also ESM, Appendix 5, fig. S2). Dates of caterpillar attacks indicated with ‘C’.

Trees in 2012 suffered unusually in two definite caterpillar attacks by the same black *Achaea* species, both coming just prior to peaks in leaf-flush (marked ‘C’ in Figs 1a and 4). The earlier flush in early January 2012 was part of the annual leaf exchange process, but the later one in early April 2012 was not. The first was in part associated with the leaf fall around 15 Jan. 2012: 21 of the 61 trees lost new leaves then. The third leaf flush in late February 2012 (Fig. 4b) had no associated caterpillar attack. What happened in late March was unfortunately not recorded, but a building up of the re-flush starting then could be surmised if this event followed the same flushing patterns of the earlier and later ones. The sequence of events suggests that leaf flush in early January enabled the caterpillar population to initially increase, and that at the end of February there was a response by the trees to the first attack. The flush at the end of April is more difficult to explain. It could have been either a further delayed response to the first attack, or a direct response to the second attack, by a now larger caterpillar population. The life-cycle dynamics of the herbivore would have allowed, however, only a second cohort of caterpillars to feed on that third and last flush for the year. The most flowering accompanied the middle leaf-flush peak in 2012, which speaks against caterpillars being around then because feeding on leaves might be expected to indirectly reduce flowering.

The presence of the black caterpillar in 2012 appears to have upset the smooth phenological stage-replacement of 2010 and led to a much-reduced reproductive output. The interval from the last year of full masting (1 year) was theoretically suboptimal, so likely the resource levels of P but not N were reduced and still below a threshold. A high N: P ratio may have helped the caterpillar outbreak. In addition, the masting pattern changed after 2011 (Fig. 1a), to the two moderate double-year form, a carryover of the disruption of the feeding in 2012, and a destabilization of the proposed inherent 3-year cycle. In the dry season 2010 mean daily rainfall was 1.42 mm, yet the year prior it was 4.66 mm; correspondingly the rainfall for 2012 were 1.95 and 2.07 mm: prior-to-current ratios of 3.3 and 1.1, respectively. Interestingly, the lower caterpillar activity noted in the other grove to the south was associated with average green-pod score of 1.7 in May 2013 (*n* = 33 trees; ESM, Appendix 2, table S1), showing that masting was more intense there than in the P-plot that year.

The second herbivore attack in January 2014 of the other unknown caterpillar species was before the second of the moderate ‘m’ mast fruiting of the double-years 2013–2014. Much less is known about its effects particularly because detailed phenological recording did not continue to end 2014. However, given that this species eats flowers, as well as young leaves (ESM, Appendix 6) it most likely reduced the number of trees able to fruit that year, even if the outbreak was contained to a far smaller known area than the successive outbreaks of the black caterpillar in late December 2011 and early March 2012.

It is instructive, once again, to look at the course of rainfall events and *rft* during and immediately after the dry seasons of 2009–2010 and 2011–2012 to find clues for the second, early March, caterpillar attack. This last came during a second short dry period when there was that unusual third leaf flush (cf. Fig. 4b with ESM, Appendix 5, fig. S2[c]). With an interval of ca. 14 days between phenology census recordings, it is difficult to say which came first; but the most likely explanation is the further dry period caused flushing, and this promoted renewed feeding, possibly from a next cohort of caterpillars that year. The dry season 2009–2010 had also several dry days at a similar time but was broken by a large wetting event (and there was no further flush after the dry season one). For comparison, the pattern of 2010–2012 shows the shorter, later-starting and not so intense, dry season for a typical non-masting year (ESM, Appendix 5, fig. S2[b])

### Tree size-fruiting relationship

Consistency in the intensity of masting over time at the individual tree level was tested in the following way. Scores for brown pods in 2007 and 2010 were matched with those for an equally strong masting in 1995 on a tree-by-tree basis. Score categories ‘0’ and ‘1’ were combined. Fruiting was significantly positively associated in both cases (χ^2^ = 10.61, df = 1, *P* = 0.031 and χ^2^ = 10.28, df = 1, *P* = 0.036, respectively). There is good evidence that high- and low-scored trees tended to remain as such over a 12–15-year period, although considerable between-trees variability existed.

Ordinal logistic regressions of brown pod scores per tree versus tree stem diameter (at last census in 2005), in 2010, 2012 and 2013, or as fallen pods in 2014, were weakly and insignificantly related though (*t* = 1.40, 0.45, –0.68 and 0.42, e.d.f. = 58 to 61, *P* = 0.17 to 0.60). As the phenology score was recording the *relative* proportion of a crown fruiting, this meant that large trees did not have proportionally more pods than smaller ones.

### Oil palm yield comparison

Access to fruit yield data for oil palm (*Elaeis guineensis*) growing in the estates of PAMOL Plc on the eastern border of the KNP (Bulu-Mundemba, Ndian) allowed a comparison with *M. bisulcata* mast fruiting under the same local climate conditions. An analysis of the monthly yields, for 2003 to 2016, showed no year-to-year differences in means attributable to changes in either rainfall or radiation, although there was a very clear repeating pattern for each of the three variables within each year (ESM, Appendix 7, text and fig. S1). Moreover, mast fruiting of *M. bisulcata* in this period (see Fig. 1a again) did not correspond to peaks or lows in this proxy measure of productivity, for either the current or prior year. This suggested that radiation and C fixation were not the primary drivers of mast fruiting.

The poor relationship with radiation also means that yield cannot be used in an inverse estimation to confirm the interpolated missing months of radiation at Bulu in 2009–2011 (ESM, Appendix 1). The Ndian estates have the same substrate and soil type as southern Korup. After initial trials in the 1960s through to the late 1970s, phosphate fertilization was discontinued because little beneficial effect on palm fruit production was found (C. E. Etta, unpubl. data). It is unlikely that much of the added P was still available in the soil by 2003.

### Stem increment growth and masting

Annual radial increments (*incr*, mm) in stem growth of 20 cored *M. bisulcata* trees ≥ 50 cm diameter in the P-plot from Newbery *et al.* [75] were matched to occurrences of masting over the 15 years of 1989–2003 (ESM, Appendix 8, fig. S1[a]). The end date was the last year which had a reliably determined annual band for all trees. Standard deviation increased linearly with the mean over years (slope [CV] = 1.14, adj. *R^2^* = 50%, *P* = 0.002), indicating a considerable variability in increment between trees in any one year (ESM, Appendix 8, fig. S1[b]). No masting was noted for 1988 [81] but that is not fully certain.

Linear mixed-effects models were run using ln(*incr*) as the response variable, trees as subjects (or ‘blocks’) and ± ‘M’ as the factor. A binomial logistic regression fit of masting versus increment was not feasible as the power, with only seven events in the period, was low. There were no significant differences for increment in either the current year (*F* = 0.79, df = 1, 279; *P* = 0.38), or in the year prior (*F* = 0.12, df = 1, 279, *P* = 0.72: increments available for 1988), or in the year after (*F* = 1.90, df = 1, 259, *P* = 0.17). Thus, larger respectively smaller increments were not associated with current mast fruiting, not (by inference) caused by it beforehand, or conversely masting did not cause a change in increment afterwards. Graphs of the increment series for each of 20 trees individually (not shown) had no common correspondences with ‘M’ years, and the differences in mean individual tree increment for fruiting type were highly inconsistent (+M: mean 2.49 ± SD 1.97, –M: 2.38 ± 1.98; differences 0.11 ± 0.77, range = –2.02 to 1.67). Mast fruiting appeared little connected to stem growth in *M. bisulcata*.

Increment was positively correlated, however, with mean daily radiation in the typically modal dry season period (using all records, i.e. 1984–2003), this ‘quarter 1’ being defined by months DJF – (*r* = 0.507, df =18, *P* = 0.023), but not in the early and mid-wet season quarters 2 and 3 (*P* = 0.32 and 0.41), although again in late wet season ‘quarter 4’ it was (*r* = 0.479, *P* = 0.033). Fruiting would have been 50% over then, by mid-September. For the year as a whole the correlation was also reasonably strong (*r* = 0.503, *P* = 0.024). Increment was not strongly correlated with rainfall in any quarter, nor overall (*P* = 0.12 to 0.70), and not with radiation in the year prior to masting (*P* = 0.10 to 0.73).

Binomial logistic regression fits of masting for 1989–2003 to mean annual core increment with either *radi* or *radi_sr_* (current or prior year for both variables) were all insignificant for the increment term (current: *P* = 0.49 to 0.63; prior: *P* = 0.46 to 0.95). The same approach applied using increment of the year prior brought little improvement in fitting (*P* = 0.18 to 0.33; *P* = 0.18 to 0.62, respectively). Thus, whilst increment and dry-season radiation remained significantly positively correlated (*r* = 0.532, df = 13, *P* = 0.041), partialling out a radiation effect was of little avail in unravelling a fruiting-stem growth interaction. These new analyses on the stem increment data provided further important opposing evidence for C not being the limiting factor for mast fruiting.

### Nutrients during the masting cycle

The newly reported nutrient data, coming from the approaches described in the ‘Methods’ section above, are combined here with other nutrient data already published (see citations within the footnotes to Tables 2 and 3). To begin with, for 1997 and 1998, and for 2011, the leaflet-to-rachis ratio of N was 1.568 and 1.524, and correspondingly for P it was 1.344 and 1.274 respectively, with means of 1.546 and 1.309. For 2011 only, the L/R ratios for K and Mg were 0.540 and 1.686, respectively. It was striking how much higher the K concentration was in the rachises compared to leaflets.

**Table 2.**
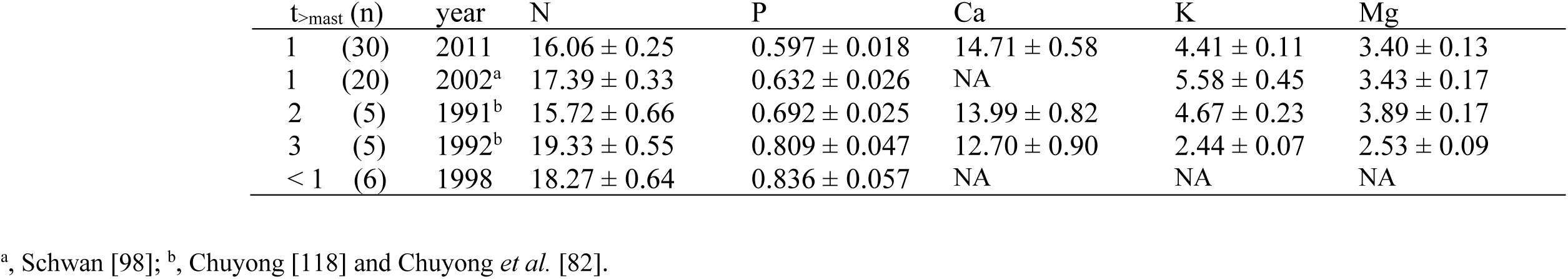
Mean (± se) concentrations (mg g^−1^) of the macro-elements in leaf litterfall of *Microberlinia bisulcata*. Collections were made in the dry season immediately following the mast fruiting of 2010, and in the second and third seasons following the earlier masting of 1989 (t). NA = no data available. Values from collection years with small superscripted letters (see footnotes) are taken from the earlier studies cited, otherwise the values are newly reported (see Methods).

**Table 3.** Concentrations of the five macro-elements in fallen mature seeds and pods (= fruits) of *M. bisulcata*. Collections were made during the 1998 masting event, together with concentrations of the same in mature leaves of this species collected in the same year of a previous (1992) event. Prominent values are shown in boldface. Values with small superscripted letters a – c (see footnotes) are from the earlier studies cited, otherwise the values are newly reported (see Methods).

|  | Concentration (mg g<sup>-1</sup>) |  |  | Amount (kg ha<sup>-1</sup> yr<sup>-1</sup>)<sup>c</sup> |  |  |  | Ratio<br>[fruit: leaf] | Reproductive<br>allocation (%)<sup>d</sup> |
| --- | --- | --- | --- | --- | --- | --- | --- | --- | --- |
|  | Seed | Pod | Leaf<sup>b</sup> | Seed | Pod | Fruit | Leaf |  |  |
| N | 14.5<sup>a</sup> | 3.80<sup>a</sup> | 22.3 | 0.770 | 3.728 | 4.498 | 41.924 | 0.107 | 9.69 |
| P | 1.80<sup>a</sup> | 0.24<sup>a</sup> | 1.18 | 0.096 | 0.235 | 0.331 | 2.218 | 0.149 | 12.99 |
| K | 7.40 | 6.13 | 5.91 | 0.393 | 6.014 | 6.407 | 11.111 | **0.577** | **36.57** |
| Mg | 3.01 | 1.17 | 3.77 | 0.160 | 1.148 | 1.308 | 7.088 | 0.185 | 15.58 |
| Ca | 3.77 | 2.17 | 13.2 | 0.200 | 2.129 | 2.329 | 24.816 | 0.094 | 8.58 |

<sup>a</sup> as reported in Table 1 of Green and Newbery [95]; <sup>b</sup> mature, green leaves, from Chuyong et al. [82]; <sup>c</sup> using estimates of oven-dry mass per ha from Table 1 of [95]: seed, pods and fallen litter: 53.1, 981.1, 1880 (kg ha<sup>-1</sup>) respectively; <sup>d</sup> RA = fruit density/(fruit + leaf amounts).

In the 1990–1992 litter sampling the bulked samples each represented 0.320 ha of forest. In 2011 sampling the trap clusters represented on average 0.368 ha. This means that each nutrient value is similarly based on ∼ 1/3 of a hectare, and the respective samples sizes of 5-and-5 versus 30 can be retained in statistical comparisons without needing to group the 2011 ones. Likewise, the 1997–1998 study used some of the same half-plots, and accordingly had the same scale as in 1990–1992. Working with the typical masting cycle of 3 years shown by *M. bisulcata* at Korup over the period 1989–2010 [11, 68], the three sampling dates together allow a series of immediately-after (2010–2011), and 1 (1990–1991) and 2 (1991–1992) years after (and thus also immediately before the next), masting events to be constructed. The two consecutive years are likely temporally correlated, but the time difference between them and 2011 is large. The further data from 1997–1998 present an interesting comparison in that masting in 1997 had not prevented a repeat even the following year. Over the successive four fortnights, concentrations of N were 18.69, 17.20, 18.58 and 18.63 mg g^−1^, and those of P were 0.802, 0.799, 0.787, 0.956 mg g^−1^.

The concentrations across the three weekly litter samples in December 2002 [98], were averaged per trap. To obtain whole-leaf concentrations, the averaged values per tree were first divided by the 1997–1998/2011 leaflet-to-rachis ratios to have indirect rachis estimates, and then the 4:1 leaflet-to-rachis mass ratio was applied. This assumed that the ratios from the 1997–1998 and 2011 studies could be used here for 2002 (which lay between them). For the complete five-element data sets and direct estimation of leaf concentrations in Table 2 (viz. 1991–1992 and 2011), years 1-3 after masting, N, P, K and Mg all differed strongly and significantly (*F* = 13.4, 12.2, 27.2 and 6.01, all *P* < 0.01) but not Ca (*F* = 1.0, *P* > 0.05).

The results show that in the inter-mast period, N was low for the first two dry seasons and then rose the year before the next masting, P increased from low to high approximately linearly in the period, Ca barely changed, K and Mg were relatively high for the first two seasons but fell in the third before masting (Table 2). This suggested that N and especially P concentrations were depleted by the last mast fruiting and were re-instated before the next event. Both K and Mg increased earlier in the interval but were presumably deployed elsewhere before masting. Despite the different sampling approaches, standard errors of means, across years per element, were quite similar. The concentration within the double-masting period 1997–1998 intriguingly lay between the 1991 and 1992 ones for N, but were higher than both for P. That P was accumulating before masting is clearly demonstrated. The rise in P at the end of the litter fall series, but not of N, in 1998 is also remarkable (Table 2). The 1998 litter was collected immediately before a masting like the 1992 one, so they could be considered comparable in this respect even though the previous mast fruiting events were 1 (unusually) and 3 years prior respectively. Averaging where possible values in Table 2 for 2011 and 2002, and separately those for 1992 and 1998, gives composite estimates of leaf concentration respectively immediately post and prior to masting as: N: 16.73 and18.80 (+12.4%); P: 0.615 and 0.825 (+33.9%); K, 5.00 and 2.44 (–51.1%); Mg: 3.42 and 2.53 (–26.0%). These inter-mast changes suggested that N and P were depleted by fruiting and restored for the next event – proportionally three-fold more for P than N, but the opposite occurred for K and Mg.

Reproductive allocation of N, P, Mg and Ca was in the order of 8% to 16% (Table 3), but for K it was considerably higher at 37%. This last value can be traced back to the relatively high concentration of K in *M. bisulcata* pods, slightly higher than in leaves. The fruit to leaf mass ratio reached almost 0.6 for K compared with < 0.2 for the other elements. These calculations do not include root allocation, however, because estimates for them are lacking, in terms of both root growth rate per year and root concentrations. Fruiting in *M. bisulcata* is much costlier in terms of K than of P. *Microberlinia bisulcata* leaf litterfall rates in the 12-month July-to-June periods of 1990–1991 (non-masting in 1991) and 1991–92 (masting in 1992) – found by integrating the areas under curves of litter fall versus time in Fig. 2d of [82] – were practically the same (1710 and 1670 kg ha^−1^ yr^−1^, respectively). Multiplying these values by their corresponding P concentrations (Table 2), led to a difference of 0.168 kg P ha^−1^ yr^−1^, which is very close to 50% of the investment in masting in 1998 (Table 3), assuming that the mast fruiting that year was similar in intensity to the one in 1992.

### Minimum temperature and mast fruiting

Over the extended time series of 29 years, masting was very weakly negatively, non-significantly, related to either the mean, or the absolute minimum of, minimum temperature (minT) in the current year’s dry season (drought-defined or DJF-quarter-year-defined, for 1989–2012, and 1989–2017; ESM, Appendix 4, table S4; Appendix 9, fig, S1). Furthermore, it was also unrelated to minT in the preceding end wet season months (SON quarter-year; ESM, Appendix 4, table S4). Graphs of daily minT across each DJF quarter year suggested no consistent differences between mast and non-mast years in when, and by how much, minT fell to its dry seasonal minimum. Drops in minT below 20 °C were infrequent and usually small (ESM, Appendix 9, text, figs S2-S5, table S1). Minimum temperatures are available from Etta *et al.* (2022). Mean daily radiation and rainfall during the drought-defined dry seasons 1989-2017 were weakly and insignificantly correlated with minT (*r* = 0.253 and 0.258 respectively, df = 27, *P* ≥ 0.18).

A brief synopsis of the Results is shown in Box 1.

## DISCUSSION

### Mast fruiting as a non-stationary process

If masting is driven by the availability of resources, and these are partly stochastic and partly trending over time, then a range of interval lengths might be accommodated for *M. bisulcata*. In reality this range is limited to 1 to 3 years. The extended time series showed that between 1989 and 1995 there was a 3-year cycle, but after 1995, as radiation increased, there came a double year 1997-98 with high fruiting – 1998 somewhat weaker than 1997, and then the cycle dropped to 2 years with a masting sequence 2000, 2002, and 2004, returning to very strong masting in 2007 and 2010 on a 3-year cycle, and finally after that the control seemingly weakened and resulted in two double-years, 2013-14 and 2016-17, which showed only moderate mast fruiting. Apart from being quite short at 29 years, the time series was non-stationary. There would be little reason to assume that it would *a priori* be stationary, especially, given the late transient cycle stage of the stand [75] and the varying climatic variables on all time scales ranging from years to decades and centuries [116, 117]. The inconstancy might be best understood in a novel way with a two- or three-factor threshold model, where C, P and K reach potentially different thresholds, simultaneously or sequentially. Thresholds themselves may not even be constant over time but relate to age, climate trends, and other tree-internal demands. When one of the resources reaches its threshold early, its effectiveness could be limited by the slower accumulation of another resource. The first resource will need to be stored or allocated elsewhere in the tree, even to then support the acquisition of the lagging resource. In time the second resource accumulates to a level that crosses a pre-set threshold, and this allows the first resource to come into operation.

The allocations of C to mast fruiting in *M. bisulcata* are substantial. In the earlier mast fruiting event of 1995, 28 out of 30 trees recorded had fruits. The average number of seeds per tree was ∼26K (maximum 92K). Using estimates of mean dry masses of seeds and pods, and mean numbers of seeds per pod, the total dry mass investment was 17 and 307 kg per tree respectively, summing to 324 kg per tree [95]. These per-tree masses translate to 53 plus 981 = 1034 kg/ha, ∼55% the mass of average annual leaf-fall [95]. Applying the individual seed and pod masses to counts made for the 2007 and 2010 events, for trees that had pods [80], gave 181 kg per tree in 2007 and 122 kg per tree in 2010. (Excluding the two zero-producers, output values for 1995 were 346 kg/tree.) On this basis the investments per masting tree in 2007 and 2010 were 52% and 35% respectively of that in 1995, illustrating an up to almost two-fold variation in masting intensity across these full fruiting events. Other more semi-quantitative scorings of masting across the rest of the time series suggest a similar range in reproductive output.

Flowering is a precondition to fruiting, but the dependence is not exact. Heavy flowering was recorded in 1992, a mast year; but in 1991 there was also especially heavy flowering but no mast fruiting [68, 118]. In 1990 there was some low fruiting (flowering unmonitored), this coming though in the year after a main mast fruiting in 1989: further, 1993 had very little flowering and consequently almost no fruiting, and 1994 relatively high flowering but only scattered individuals fruiting [90, 95]. From the later, more detailed, phenology recordings, 1995, 1997, 1998 and 2000 had strong flowering and were masting years, whilst 1996 and 1999 flowering was practically absent and had no masting [11]. Whilst valvate bracteoles protect flower buds before anthesis, it is possible that the intense occasional shower could damage flowers enough to interfere with pollination. This is the surmise for 1991, two years after the 1989 masting: fruiting was probably checked and masting was delayed to 1992. In 2004, the dry season started 43 days after 1 December 2003, on 12 January 2004 [11]: recall that the dry season for a given year starts in December the year prior. Flowering peaked over the week before and after that date. But in 2012, the season started 21 days in, on 22 December 2011, which was 1 month before flowering started in 2012, and ended another month later. (In both 2004 and 2012 there were no heavy showers that could have disturbed pollination in the first halves of those dry seasons.) These facets indicate that, firstly, occasional external and unpredictable influences can alter the processes behind a masting cycle; and secondly, the timing of flowering (also linked to the timing of leaf-exchange) can change by up to a month with respect to the start of the dry season. If internal tree nutrient levels do, via deterministic physiological mechanisms, trigger the flowering – which leads most often to fruiting at some level of strength [11], masting remains also subject to climatic stochasticity causing interference to pollination (may be once in 10–15 years) and variability in dry season timing and intensity (every year).

Abortion of small immature green pods falling to the ground was not seen in the 29-year period. It remains possible that outside of the three more intensively recorded periods this phenomenon was missed. If in occasional years there was strong flowering, but no mast fruiting followed, then usually those trees that did develop any pods carried them through to maturity. The control of masting therefore most likely happens before flowering (see [11] for further discussion). After that seed failure is simply a chance unpredictable cost within the evolved life history schedule.

### Nutrient resource interdependencies

Analysis of the seed and pod nutrient concentrations and amounts in relation to those in mature leaves indicated ratios of between ∼ 0.1 and 0.2 for N, P, Mg and Ca, but substantially higher at 0.6 for K. This difference is very largely due to the low concentrations of the first four elements, and a much higher one for K in the pods. Pods collected were recently fallen. Detarioid pods are hard and rather water-repellent for several weeks, so it is very unlikely they adsorbed much K coming down in canopy throughfall (G. Neba, D. M. Newbery and G. B. Chuyong, in prep.), or were they leached to any extent. Had that been a cause of an increase, then Mg should have increased too, yet it did not. Presently it is difficult though to explain why so much K was being invested in pods. The comparative leaf K values, coming admittedly from a retranslocation study where replication of bulked samples per date was small, do closely match those of mature leaves of plantation-grown *M. bisulcata* trees (averaging ∼ 5.8 mg g^−1;^ at ages of 13 to 18 years; D. M. Newbery unpubl. data). Whilst ectomycorrhizas are strongly linked to P acquisition and uptake in the P-poor soils, K is also very low in availability at Korup [72], and the surface-soil root mat (with dense mycorrhizal networks) is an efficient collector of K (and Mg) coming down in throughfall (G. A. Neba, D. M. Newbery and G. B. Chuyong, unpubl. data). In plots at the same site, Chuyong [118] found N, P, K, Mg and Ca concentrations of 13.8, 2.09, 6.41, 2.22 and 3.50 mg g^−1^ respectively in *M. bisulcata* seeds that fell in 1992. Pods were not separated in the 1992 general collections but included within ‘all reproductive parts’ in the HEM plots, those with high abundance of *M. bisulcata* had a mean K concentration of 6.04 mg g^−1^. Although other elements in fruits were albeit a little higher than for *M. bisulcata* alone in 1998, the earlier data confirm the K values of interest well. Investing P in seeds would presumably be important to the survival of establishing seedlings up until the time they could become infected with ectomycorrhizas and link into the mycelial network. Could it be that also K, not P alone – as for some time thought [73, 85], is a second key controlling element of masting in the Korup groves? Several indications suggest that P and K might act synergistically.

The main pattern explaining the masting time series was the sequence of relatively wet dry season followed by relatively dry one. There was no evidence of water limitation to large trees in the dry seasons because of their ability to root deeply [79, 81], so this points to a soil or nutrient factor governed by soil moisture content in the upper soil layer, where fine roots and ectomycorrhizas form a root-mat and are most abundant and active in nutrient cycling. What is it about relative dryness and wetness of the dry months that leads to these relationships? Possibly dryness affected the fine roots and ectomycorrhizas. The drier the dry season, then likely the more restricted the root activity under limiting surface soil water conditions, which would possibly signal stored P in the ectomycorrhizas to be moved to intermediate locations ready for fruiting, and less would be used in maintaining new root growth. In a relatively wetter dry season this root activity would continue longer and more effectively, so that any additional P stored in the roots would stay there and not be signaled away to fruiting. Nevertheless, the main mineralization and uptake occurs in early weeks of the wet season [83], and this would be similar each year once the soil water content was high enough. This dryness is thought to be a proxy for a mechanism that triggers translocation, but it is one that cannot operate without the previous year acquiring more P and setting up a form of ‘relay’. It is interesting that moderate ‘m’ events occur in double-years and twice later, mirroring the two strong ‘M’ years 1997–1998.

At Korup in the HEM plots, sand, silt and clay proportions were 77%, 9% and 14% respectively [81]. There exist very few published water content vs water potential curves for sandy loam tropical soils, but at a site in NE Brazil with a very similar physical composition to Korup, Fisher *et al.* [119] found that a change in SMC (at 30 cm depth) from 20% to 12% (cf. the wet-to-dry-season change in Fig. 3) led to a decrease in soil water potential (SMP) from –10 to –100 kPa – which is very much higher than the generally accepted value of – 1500 kPa where wilting starts. Such a translation across sites should however be treated cautiously because any SMC-SMP relationship is finely dependent on the exact physical composition of the soil [120]. Wilting of understorey treelets and shrubs, even in the strongest dry seasons, was very rarely observed at Korup. Nevertheless, the drying of the top surface soil with the root mat might still be expected to affect nutrient uptake capabilities in the dry season. In those dry months, the surface leaf litter is dried out, and in many places the sandy-loam soil just below the organic layer is visible, leaving the root mat exposed. The implied sensitivity of fine roots and ectomycorrhizas of *M. bisulcata* to surface soil water conditions might in part result from this species very pronounced buttressing and an extensive lateral root system lying just below the surface soil mat [79].

A cycle of 3 years would most likely allow a restocking of P, not only enabling growth of other tree parts, but also storage of P in wood rays and roots ready for fruiting (as the model later explains). A 4-year cycle could arise but, *ex hypothesis*, which would need a sequence of two dry dry-seasons: none was recorded, however. The regression modelling gave an indication of year-to-year increases in radiation, and positive deviations from the smoothed trend of radiation with time. Changes were relatively small, radiation varied rather little from year to year, both for the dry seasons and the years as a whole. The additional radiation probably did not contribute to much increase in photosynthesis which would cater completely for the extra demand in years of masting, above the potential competition for C by leaves, stems and roots. Variation also in start and duration of the dry season, with the fairly steady daily radiation inputs, meant that there was probably sufficient radiation supplying C on top of tree storage, to meet the high dry mass investment in fruits. Carbon supply was therefore likely not the limiting factor to masting. One way to resolve this possible contradiction is to propose that, internal to the tree, there was strong temporal division in the allocation of C between rooting (one year, wetter) and fruiting (the next, drier). Further, the lack of a relationship between proportion of crown fruiting and tree size may be further evidence that C is not the predominant limiting factor for mast fruiting – contrary to what was originally suggested by Green and Newbery [95], if it can be assumed that large trees store more NSCs than small ones. The lack of year-to-year differences in oil palm yield relating to radiation is also moderate evidence of C from photosynthesis not being so critical for their yielding either. Palms can store up to ∼20% of their C in stems from year to year [121, 122]. The lack of correspondence between oil palm yield and *M. bisulcata* fruiting suggests no common climatic driver and points indirectly to nutrients as the key factor in the forest, though possibly not P alone since palms did not respond to P fertilizer.

The very few studies on C and nutrients involved in mast fruiting of dipterocarps that have been made largely support the thesis of this paper. Defoliation and girdling experiments on branches of *Dryobalanops aromatica* trees [123] indicated that flowering drew on C stored in local branches and the filling of fruits came from nearby leaves. This confirms the many similar conclusions for temperate trees mentioned in the Introduction. Relative allocation of P to fruits was shown to be higher than of C and N for of another masting dipterocarp, *Dipterocarpus tempehes* [10], again emphasizing the importance of P in reproduction, especially on P-poor tropical sites. During a different masting event for *D. aromatica*, P concentrations in branch, stem and roots steadily decreased during flowering and fruiting (but not in twigs and leaves), N fluctuated slightly (in all parts) and, whilst K decreased during flowering, it increased during fruiting (in all parts) [124]. Of the P needed for fruiting, 68% was estimated to come from branch storage, compared with only 20% for N and very little for K. This was the first time root-stem-branch stored P was shown to be important for masting, and the result corroborated the hypotheses of Henkel *et al.* [9] and Newbery *et al.* [11]. A further study at the forest stand level provided more support for P and K both being key nutrients in mast fruiting. Aoyagi *et al.* [125] followed nutrient fluxes in soil, stems and fine litter in a lowland dipterocarp forest during one such event and for 4 years afterwards. Fluxes of P and K were much higher in total fine litter in the mast year than later years, whilst those of N, Ca and Mg altered little; the higher P and K was coming from the high allocations to fruits. Interestingly, P and K concentrations in coarse roots – but not stems – of the most abundant dipterocarp, *Shorea multiflora*, decreased more in flowering than non-flowering trees. This might indicate root storage of P and K ahead of stem and branch storage, in preparation for fruiting. After flowering in *S. curtisii*, Yeoh *et al.* [126] detected a decline of twig P which took a year to recover, highlighting further the likely important role in mast fruiting in dipterocarps. Kitayama *et al.* [127] have more generally demonstrated in NE Borneo that across a P-use efficiency gradient, forest trees allocate proportionally much more P than C or N to reproduction (in relation to total allocation), even at P-poor sites.

### Masting and carbon supply

The internal distribution of C between potentially competing sinks within each tree has a further interesting aspect. A majority of the *M. bisulcata* trees in the grove are presently of large to very large (≥ 50 cm stem diameter) sizes which means they likely have considerable respiration and maintenance costs. Carbon at least would not be so readily available in an average year to sustain such a heavy fruiting without causing deficits in growth elsewhere in the tree, perhaps in fine root and ectomycorrhizal production. This would lead, hypothetically, to an even stronger competition between rooting and fruiting, once stem growth is partitioned aside. That seedlings and saplings survive very poorly close to adults, led to the proposal of a reverse drain operating towards the old adults especially in mast fruiting years, for P as well as for C [68, 87, 88, 128].

Previous estimates of leaf litterfall were 1880 and 1380 kg ha^−1^ yr^−1^ in non-masting 1990 and masting 1989 respectively (the first estimate is used in Table 3), and came from Green and Newbery [95], who in turn based their calculations on January-to-December, plot-level, leaf litterfall estimates given in Newbery *et al.* [81]. The 1990–1992 data later indicated 39% of that litter was from *M. bisulcata* trees. This 0.74:1 ratio may have been an overestimate, because it used calendar year estimates which would have missed linking November-December litterfall with that in January to April prior to masting, and the newer 1990–1991 species-specific data were more precise. Or it simply reflects variation in litterfall, viz. 1380, 1880, 1710 and 1670 kg/ha over the four years 1989 through 1992, which suggests that C supply was likely again not to have played an important role in determining mast fruiting since in around one masting event there was an implied cost to leafing and around the other none. The fruit-to-leaf amount ratios and reproductive allocations percentages in Table 3 may well be ∼10% lower than the true values, if the 1990–1992 litterfall estimates did apply. The 1990–1992 data also display a small shoulder in May 1992, possibly the result of the last older leaves being exchanged for the new green pods.

Green pods stay on the trees for more than 5 months and therefore will contribute C to their structures and to developing seeds. This is likely an important source, and pods and seeds may indeed be almost autonomous in this respect. Pods are also held up high on the outer canopy edges where their photosynthesis is maximized [80]. Bennett *et al*. [129] reviewed the role of pods in their photosynthetic capacity, and Zhang *et al.* [130] provide a detail analysis of C balance in the pods of *Medicago sativa* to show the considerable C inputs that are possible. This source of C would make the *M. bisulcata* trees at masting even more independent of C stores within the rest of the tree. It could explain the effect of raised radiation in the current fruiting year when compared with the one prior.

That stem increment growth was positively related to current year radiation, but mast fruiting was not always, or at least not so strongly, suggests that C from photosynthesis was being used for stem growth and in mast years it was additionally contributing to mast fruiting. An internal trade-off between root growth plus ectomycorrhizal activity to enable the uptake of nutrients allowing nutrient uptake, and an allocation to fruits and seeds when cued, may well be the basis to the masting cycle. In the year following a masting, relatively more C would be allocated to roots because, if fruiting is ‘switched off’, roots become the main alternative sink; and then if in the second year there is sufficient P to cross the threshold, with some increase in radiation and extra C input, a next masting would occur after 2 years. If P accumulation were limited the tree would wait until the next year, forming a 3-year cycle. Extra C available in that second interval year might then determine the intensity of the masting (‘M’ or ‘m’). A notable difference between P and K is the rate of change in concentration in the leaf litter as the interval between mast fruiting progresses: P increased slowly over 3 years, but K rose rapidly then fell lower than at the start. This hints at faster recycling of K than P, with a stronger withdrawal of K than P at senescence later as storage cells were being refilled prior to masting.

### Rainfall and nutrient uptake

The effect of mean daily rainfall in the dry season is dependent on the pattern of rainfall distribution over time, through the dry season and just into the early wet season weeks, which is up until the 100 mm threshold defining the season is exceeded. The first week’s rainfall governs root growth and ectomycorrhizal physiology, and the later ones connect to the early wet season and the throughfall which primes the start period of optimal leaf litter decomposition [83, 84]. A dry season may well limit root activity, and it also would delay decomposition: one which was short would only affect the former. On the other hand, a relative wet dry season will be expected to restrict rooting and ectomycorrhizal activity much less and encourage the earlier start of decomposition, both processes leading to better nutrient uptake into the trees. Ectomycorrhizal activity doubtless plays a role in P uptake, and the regrowth and maintenance of the root mat itself determines the level of retention of K leached from the canopy and its uptake.

An intriguing aspect in connection with the P-supply for fruiting is how P gets taken up and stored in inter-mast years. In the dry season of 1988–1989 and 1990–1991 there were pronounced peaks of P in fallen leaf litter in the HEM but not LEM plots [81]. This peak was absent in 1989–1990, directly after the heavy mast fruiting. The HEM litter would have been composed of ∼80% *M. bisulcata* leaves [82]. In a relatively dry dry-season with likely more die back of fine roots and ectomycorrhizas than in a wetter one, P might be translocated out of these fine roots and ectomycorrhizas, going into storage for fruiting. Altogether, the integrated aspects of this P nutrient cycle do provide reasonably strong support for P storage and use leading up to, and during, the masting events as proposed by the original PACER hypothesis.

Another result providing important evidence for P-storage is from the comparison between concentrations and amounts in the years across masting intervals which suggests further that the 50% of P used in masting was coming from retranslocation of P out of senescing leaves a few months before. This P was probably being stored in branches. By difference it might be deduced that the other 50% was older and coming from local branch/wood storage built up in the prior 1 or 2 years, still within the interval. That result would fit with the notion of a temporary store for P and not all of the P needed for masting could be supplied in the current year alone. Rosell *et al.* [131] in a study of a range of tropical tree species found the inner bark of branches was especially important for NSC storage and, although not yet tested, this may be where P is also held, ready for fruiting. Jones *et al.* [132] indicate this as a possibility for premontane tropical forests.

If P in leaf litter is relatively low in concentration after allocation to fruiting [81; and new evidence in this paper], then the fast-forward cycling proposed by Chuyong *et al.* [82] could be temporarily slowed because mineralization by saprophytic fungi would have a low-litter resource to decompose. Root activity would either need to become more intense and efficient to take up the now lower soil levels, or at this time the ectomycorrhizal activity would come more to the fore by not only accessing low molecular-weight organic P molecules efficiently but also by effectively reaching low-concentration inorganic P sources further way from roots. This impasse would strongly select for the several co-adapted root mat and ectomycorrhizal traits seen in the field in order to achieve restocking of P. The rise in P accumulated just before masting could lead to higher P in litter falling that year which on one hand would mean fast re-uptake but on the other appears to be a dissipative risk when the perceived strategy is to save P (in the tree) for immediate use in fruiting. However, the ectomycorrhizas being part of the whole tree system, and cycling highly efficient, P is being ‘saved dynamically’ within the nutrient cycle. Also, N and K would in parallel complement C in stores – the latter benefitting from the small increase due to radiation enhanced photosynthesis, all three also needed for seeds and pods. That P was high in falling litter just prior to masting might furthermore indicate that P stores were quite full after a 2-year interval, and some of that ‘extra’ P was also coming from root retranslocation.

The analysis of mast fruiting in 1989–2004 [11] used 150 trees across the whole P-plot, but the later recording 2009–2014 used a subset of 61 trees in the eastern half, an area 100–500 N by 1000–1200 E, on the higher ground with denser stands and larger trees. The masting time series could potentially have been confounded by sampling extent changing over time, because for the earlier half (1989–2004) it was found that masting was significantly stronger on the relatively higher and drier sites of this eastern part of the P-plot compared with the rest of the plot [11]. Allaying these doubts, the full and strong masting in the second half (2007 and 2010) were confirmed by seed trapping across the eastern 25 ha of the plot [80]. The two double years, in the second half of the time series, were only moderate masting despite being at the more favorable topographic location.

The very strong masting achieved by *Dicymbe corymbosa* in Guyana, with 3–4 t dry pod and seed mass per ha, were associated with pronounced El Nino Southern Oscillation (ENSO) peaks in radiation [9, 133]. However, this ectomycorrhizal detarioid, although phylogenetically very close to *M. bisulcata*, is unusual in that it reproduces annually also by sprouting, which must impose a considerable demand on C and nutrients for growth. For this reason, it might be conjectured that C stores are probably being continually used up and hence, to achieve mast fruiting, a dependence on an additional boost of incoming radiation in those ENSO years would be necessary. Again, in contrast to most African detarioids (including *M. bisulcata*), *Gilbertiodendron dewevrei* – which forms monodominant stands in Central Africa – exhibits mast fruiting which is spatially unsynchronized and at irregular intervals [134]. Its individual seeds are very heavy (∼30 g), so the mass allocated to reproduction per tree is probably far greater than for *M. bisulcata*. Compared with SE Asia and C. and northern S. America, the lack of a strong ENSO signal in C. Africa is pertinent [135, 136]. Masting in dipterocarps may well be for different suite of factors from those controlling the detarioids, achieving an evolutionary stable strategy along an alternative channel.

### Interference effect of herbivores

The loss of a 2- or 3-year cycle in masting after 2011 suggests that the rhythm of wet/dry dry-season release of nutrients needed for full masting was disrupted by a combination of lack of an alternating pattern in rainfall plus heavy caterpillar attacks. This latter unforeseen intervention, combined with an unusual climate regularity over 3 years, acted as a ‘quasi-treatment’ in that C gain and storage were likely temporarily reduced. From the above argumentation it was not so much the decrease in C to supporting fruiting that was alone important but also the loss of C input for roots to allow P to be taken up and accumulated even in the relatively wet dry seasons. With some commitment to flowering already, leaf grazing would have nullified any C input from the local branches attacked to fruiting, requiring the tree to draw on stores in the branches. Nevertheless, some fruits resulted, indicating a means by which tolerance to herbivory could have occurred [115]. The caterpillar attacks may have interfered with the accumulation of P and thereby interrupted masting in full, leading to the unusual double ‘m’ years. The evidence of an herbivore effect is strongest for fruiting in 2013–2014 because of the detailed phenology data in 2011–2012. It is tempting to suggest that something similar might have been happening in 2015 before the very similar masting pattern of 2016–2017: alas, herbivore and tree phenology records were not possible during those later 3 years. A further unknown aspect is how herbivory may have induced stress hormones in the trees which, interacting perhaps with K levels, affected cytokinin concentrations.

The pooled field notes and observations suggest two lepidopteran herbivores occur in the crowns of co-dominant *M. bisulcata* and *T. korupensis* in the natural old-growth forest of Korup. Both species have pinnate leaves – *T. bifoliolata* does not – and similar stem densities at ∼ 3.5 per ha for ≥ 50 cm DBH [68]. But *M. bisulcata* trees are the greatest in size with a near-zero mortality rate over recent decades [75]. If caterpillars indeed fed only on expanding leaves (or flowers), then severe damage could ensue quickly, within 1 to 2 weeks, inducing premature abscission (or lost pollination). For example, within 1 week caterpillars defoliated young leaves in crowns of two monodominant leguminous tree species in rain forests of Brazil and Gabon [137, 138]. Such phenomena are quite easy to miss though. This could explain why the black caterpillar was not noticed before at Korup, but it is unlikely the same reason for the yellow caterpillar given the extensive multi-decadal phenological monitoring of *M. bisulcata* [11, 68, 115]. The detailed monitoring between 2009 and 2013, like that between 1995 and 2000, did not note any caterpillar attacks. To have an importance they would surely have been seen before from their feeding effects (droppings, noise of chewing, fallen individuals, hanging threads, frass in traps). Many other periods of intensive work back to 1984, focused on the dry months and early wet season when caterpillars would occur, did not record them. Whether these outbreaks are triggered by favourable climate factors, or palatable leaf chemistry or less parasitoid pressure, either perhaps induced by drier conditions [139] or simply mounting abundance over years [140], is unknown. Herbivore attacks to flowers, occurring stochastically, might be viewed as being similar in end effect to intense rainfall events disrupting pollination: both limit reproduction.

### Possible role of minimum temperature

Dry season minimum temperature (minT) did not show any pattern in marked declines towards minimum minT across the years 1989 to 2017. There was no correspondence between mean or minimum minT and mast fruiting of M. bisulcata in the following wet season which would suggest that minT was acting as a cue for, or effecting, phenological changes. Flowering occurs annually at Korup, except when other factors such as heavy rain showers [68] or caterpillar attacks (Fig. 4) interfere. This is a common evolutionary stable strategy in seasonal tropical forests. As a leaf-exchanger, the new leaves and flowers come at a similar time during the dry season, just after or as the old leaves are shed. The driving factor is drought, its severity, start and duration determining when leaf exchange happens. Concomitant with the declining water supply as rainfall decreases, cloudless skies lead to a high incidence of solar radiation and consequently a lowering of temperature at night. A physiological mechanism by which minimum minT causes anthesis still remains unknown. At least at Korup, for the period studied, decreases in minT appears to be too small and for far too short time to be effective. They give a signal which is most likely too unreliable to allow a selectable response by the trees in such a stochastically variable climate. It remains unshown whether minT was having a latent effect.

MinT was increasing steadily from 1989 to 2017 (ESM, Appendix 9). The two double years of moderate masting in 2013-2017 might have been caused by higher minT. Such a general effect of rising minT has been put forward to explain recent declines in flowering in many tree species across the tropics because of global climate change [141, 142]. However, the two prior mast fruiting events were very strong. In the case of Korup, the moderate fruiting of M. bisulcata might also be attributed to caterpillar attacks, as argued above, which would confound a rising minT effect. Alternatively, caterpillar attacks might have been caused directly by increasing minT, or indirectly by it on new leaf production. Unfortunately, flowering was not recorded for all of the later years (2010-2017) in the present study.

### Refined updated hypothesis for mast fruiting in detarioids

The mast fruiting cycle in *M. bisulcata* could thus be explained in terms of the availability of three principal resources which govern the timing of reproductive output: the limiting resource P, water as a controlling resource, and C as a qualifying resource. If sufficient P accumulates within 2 years to trigger flowering and there is sufficient C coming from storage laid down over the last year or supplemented due to a positive radiation difference between prior to current year, then fruiting is to be expected. But when P is slower at accumulating, the interval is extended by one year and fruiting happens within 3 years. It is highly likely that in the latter case C would still be sufficient either from reserves laid down in the tree in the previous year and/or a small surplus the current year of fruiting. When there is sufficient P accumulated after one year but not quite enough C for a full masting, fruiting may follow anyway at a moderate level; and in doing so it is likely that not all of the accumulated P would have been allocated ‘as planned’, and the part remaining, plus last year’s continuing uptake, will sustain a second moderate fruiting. According to these resource requirements, it is difficult to see how the pattern could be ‘m-M’ or ‘M-m’, and neither of these were observed: in the first, there would be insufficient time after a ‘m’ fruiting to build up P so rapidly to have an ‘M’ one, and in the second, the ‘M’ first would exhaust P levels down below a threshold and no ‘m’ would likely be possible either. The pattern M-M did occur once, however, at the time of an exceptional rise in radiation levels over a 3–4-year period: it can only be hypothesized that there was unusually even more surplus C for extra root growth and P uptake in both the year preceding the first ‘M’ year and during the second one. The main processes leading to mast fruiting in *M. bisulcata* are schematically shown in Fig. 5.

**Fig. 5.**
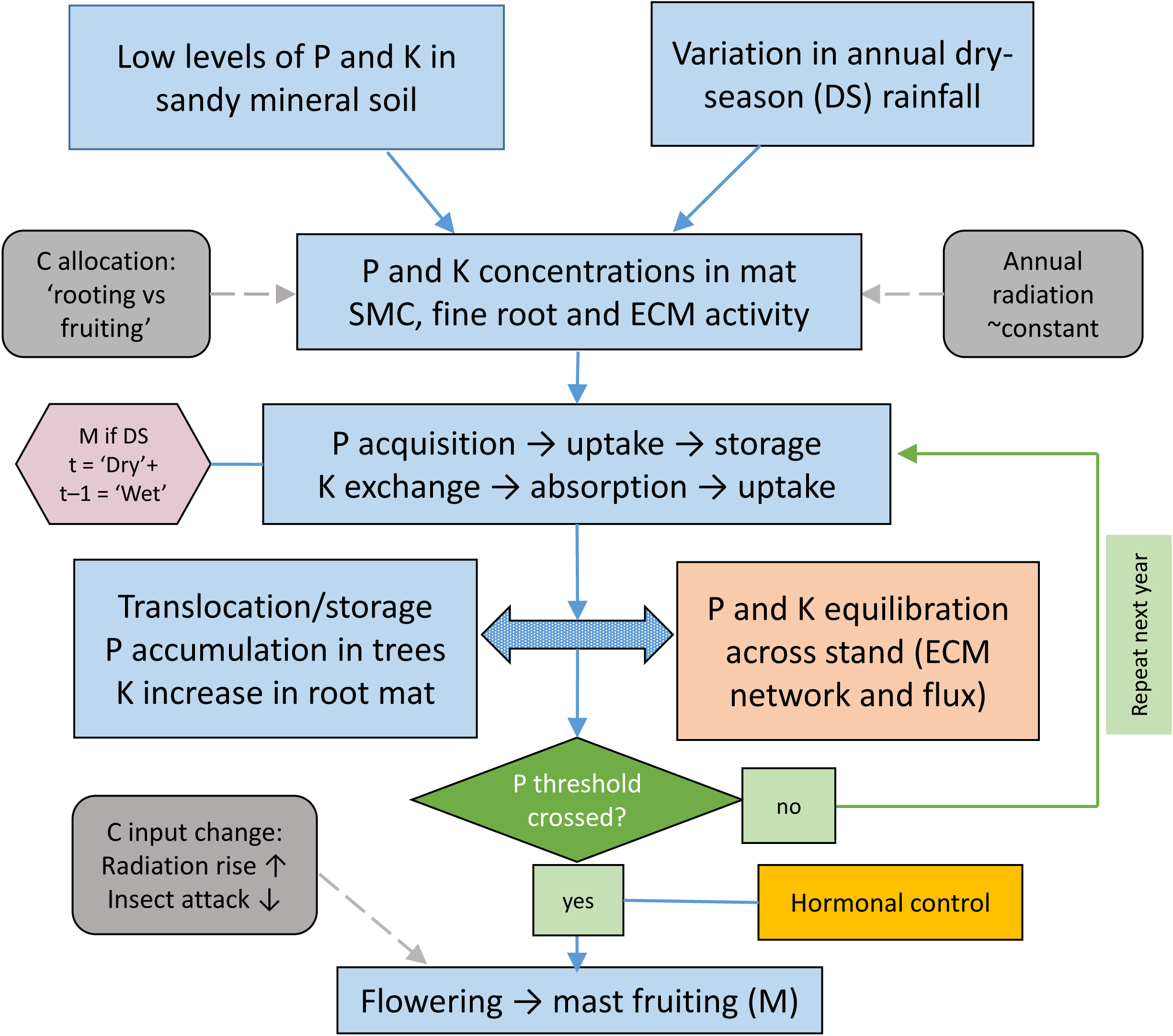
A schematic flow diagram of the interrelated hypothetical processes behind mast fruiting (M) in the grove-forming population of *Microberlinia bisulcata* at Korup. The boxes with light blue shading are either empirically established in the present study, or at least can be strongly inferred from the literature. They involve the deterministic factors soil phosphorus (P) and potassium (K), and the stochastic effects of dry-season (DS) rainfall; SMC means soil moisture content, and ECM stands for ectomycorrhizal. Those in grey concern radiation and carbon (C)-supply, dynamics and allocation. The box in pale brown-orange concerns the likely important role of ectomycorrhizal networks and nutrient fluxes within the root mat, which are hypothetical and require empirical confirmation. The central double-headed arrow represents the interplay between within-tree and between-tree processes. Green shading and lines indicate the proposed threshold triggering flowering and fruiting, and repeated annual uptake and storage over successive years within the masting cycle of typically 2-3 years. Hormonal control of masting, shown in yellow, is strongly suspected: it also requires critical testing. The small polygon in lilac, centre-left, is the primary condition for mast fruiting to occur, as shown by statistical model fitting in this paper (t is time in years). It is the process-link between the start and end of the diagram. Changes in the degree with which flowering results in fruiting, and in the intensity of fruiting itself (e.g., when lowered to ‘m’), are explained in the main text.

An interesting aspect is the feedback control that might operate by way of C going either into fine roots and ectomycorrhizas, which would enhance nutrient uptake, or into wood storage (stem, branch and large root), which could sustain a next fruiting. The 2- or 3-year cycle is simply a consequence of the contingent conditions of these trees growing in low-P soils aided by ectomycorrhizas. Rainfall and radiation are stochastic processes, to some extent inversely related, which overlies the intrinsic deterministic cycle: the important difference between them is that rain, acting primarily on the soil, facilitates P release, acquisition and uptake (needed to accumulate P), and whilst C is needed also for fruiting, it is essential for root production and maintenance of ectomycorrhizas that makes this P uptake possible. That root growth should always precedes fruiting in a masting year, the first in late-dry into early wet season (March–April), the latter in late wet season (September–October) decides the sequence unambiguously: C in the dry season will go to the root sink first and what remains stored, used for stem growth or reserved for fruiting when it happens. The low P supply is therefore the main determinant: C must be sufficient but does not appear to be decisive for the interval length. Although N and K will also be involved the evidence more strongly supports the P thesis. Nitrogen is not limiting at Korup with ample N-fixation and aerial input [81], and whilst K follows P in being intrinsically quite low in supply in Korup soils, the root-mat appears to be highly adapted to catching and retaining canopy-leached K and so enabling its efficient recycling (G. A. Neba, D. M. Newbery and G. B. Chuyong, unpubl. data). The K allocated to pods could, like N, be accommodated by wood ray storage.

Carbon and N will be important for masting, even when P and K are limiting and are the key storage resources, because all elements will be needed in their stoichiometric ratios in order to fill pods with seeds. As in other plant physiological problems, simplified reductionist approaches do not recognize the integrated nature of growth and reproduction. Carbon (sugars) and K are required for transport and storage, while P is bound into C-H-O structures which act as ‘carriers’. It is possible though that a small increase in C from radiation might have a non-linearly disproportional benefit to P transport and storage. Understanding masting physiology requires an integrated and dynamic approach, one of shifting sink strengths over time. After a last masting, roots would likely act as the strongest sink, then the stems (and rays and bark) and branches – part growth, part storage – and finally reproductive parts. No abrupt switches in C, in contrast to a threshold switch for P, but a gradual altering of the components over a 2–3-year cycle occurs. For this to operate it is necessary to hypothesize feedback from the C-depleted branches to the more C-depleted roots. With the fruiting sink removed, C should move from stems to roots. Whether this needs additional hormonal control is an interesting question. It remains a conundrum as to why ‘in the background’ to the rooting-fruiting axis, stem wood growth is positively correlated in a quasi-independent way with radiation. Stems may be working perhaps as the storage stabilizers, being placed centrally between shoots and roots. As they grow, they store and balance the terminal demands. The levelling adjustment of C between the first and second interval years could perhaps explain the lack of clear correlation between fruiting (and inversely rooting) with stem growth increment. The time sequence within the year is also important here. In the second (drier) year of fruiting, C is not, *ex hypothesis* predominantly going to roots in the early wet season but more to stems for storage, and en route there for filling fruits later in the wet season (Fig. 5).

If radiation were to increase during an increasing part of a cycle or as a longer-term trend, and C were alone the limiting resource for masting, then fruiting would come closer together and this would defeat the putative economy-of-scale selection. In this respect a non-stationary time series offer advantages when matched to an also non-constant climate environment. On this basis it would seem also unlikely that C could be the key resource for *M. bisulcata*. Further, C available per tree will decrease with tree age after the point of maximum crown extension because of the higher respiratory loads incurred in maintenance of the whole tree. This may be connected with the observed crown thinning of the largest trees (J. M. Norghauer, unpubl. data). When seed production was substantial, in proportion to the high flowering per crown then there would be a deficit of C to fill pods, or pod production would decrease, or another cost incurred (e.g. at the roots or wood production). But the evidence was that in *M. bisulcata* the relative crown-cover with pods remained constant with size and, by broad inference, age. As a tree grew to have a larger root system with ectomycorrhizas to maintain, to acquire P and retain K, when absolute pod and seed output rose then the demand for P and K would have also. As with C and N, we can expect that P and K are stored in ‘quick release’ forms in wood rays and bark of the stem and the main branches. A phenomenon often remarked is of one part of a large crown flowering and fruiting much more than the others, which could be explained by a correspondingly more active root system of the tree, and a matching storage of nutrients in the wood and branches, on that side: see [79] for the strong crown asymmetry in *M. bisulcata*.

The arguments going from nutrient concentrations in fallen litter through to the possibly the trees building up and storing P and K before masting are tenuous because of the many uncertainties about the processes involved. If some translocation occurs out of leaves before they are shed, then why are the P levels so high just before masting? If P was being actively stored for fruiting, it might be expected that it would be lower in concentration in the fallen leaves. However, the high concentration reflects that all the trees were well stocked with P by that time, the stores were full, or even over-full, and there was still plenty of P to fall. From earlier work, it was established that ectomycorrhizal species, especially *M. bisulcata*, drop P-richer leaf litter than non-ectomycorrhizal ones [81] partly because the retranslocation was less [82], and it was argued then that this allows the decomposition process to operate without P limitation, the ‘fast forward’ strategy. However, the concentration of K (to a lesser extent that of Mg also) reached a peak mid-interval and then fell just before masting. This could mean that K was being translocated into storage before fruiting. In general, the export of K and Mg from leaves of trees is usually low compared with that of N and P. Understanding the differences in dynamics of P and K in this context is highly challenging, and also it explains why N showed no clear mast-interval trends. Whilst K was likely building up on soil colloids in wetter dry seasons, mineralized from fallen leaves, dead roots and old pods during decomposition, and that this process was more limited in the drier dry seasons, suggests that K must also have been transported and actively stored in the tree between the prior and mast years. To recycle more K via litterfall would involve a potentially high loss due to leaching when the cation exchange capacity of the soil has already been reached, compared with the seemingly tighter and ‘safer’ direct access by ectomycorrhizas to organic P. This reliability aspect therefore favours P over K as the better controlling factor in masting (Fig. 5).

### Phosphorus storage and control

What could the likely form of storage of P in trees be? Since Bieleski [143] the most likely candidate is inositol hexa*kis* phosphate (= phytic acid; C_6_H_18_O_24_P_6_). It has been shown many times to be the main storage form in seeds and has been detected in stems and roots. It forms salts with Ca, K, Mg, Mn and Zn, and in soils occurs commonly bound to Fe. It occurs in 19 hexa-structures which are interconvertible [144]. It is viewed as the most frequent P-storage form when P is in excess, that is in passive storage; the P is readily retrievable and mobilizable. Other molecules can have seven or eight atoms of P but seem less common [144]. Inositol hexa*kis* phosphate is involved in cell metabolism, in ATP regeneration, RNA export and the repair of DNA. It is prominent in cell storage biochemistry, which may make it a candidate for *active* storage. Yet very little has been researched in this area, and accounts for trees are very sparse. A notable exception is the study of Kurita *et al.* [145], showing P storage in *Populus alba*, followed by release in spring, and indicating perhaps a common function in many deciduous tree species. It is conceivable that the same mechanism could operate in seasonal tropical trees and even be a basis to P storage for mast fruiting species. How inositol hexa*kis* phosphate is linked to polyphosphate formation in roots and mycorrhizas and comes under hormonal control remains unclear.

In an experiment with *Eucalyptus grandis* seedlings, comparing low and high P media, Mulligan [146] detected stored P in the form of esters and residual P (mostly phosphorus proteins), especially in roots, and more so the low P medium. The idea is that these stores buffer the plant (maybe actively stored), from there P can be mobilized and sent to other stores in the shoots. In a similar vein, Netzer *et al.* [147], comparing mature *Fagus sylvatica* stands – one on high and one on very-low P available soils – found that in the latter P built up in roots in late summer, and from there was moved on to stem and branches, suggestive of at least two storage sites in a series enabling adaptation to the poor site. However, hardly anything is known about how P stores are internally controlled, or how hormones signal transfers of P [148], which could give even a lead as to how P might be built up in stores in masting intervals, and once a threshold level is met how the switch to flowering is signaled.

Much evidence suggests that senescence in leaves, and possibly pod formation, may also involve cytokinins [149]. Chiou and Lin [150] have proposed different hypothetical models for inorganic P being regulated by cytokinins and sugar levels. How these promising models can be extended to storage and release of P has yet to be worked out. The strongest candidate for the elusive ‘florigen’ is now widely held to be a flowering locus T [FT-] protein, which again appears to be controlled by a cytokinin [151]. Cytokinins act at long distances throughout the plant and are bidirectional in their effects, both of which might hypothetically implicate them in inositol hexa*kis* phosphate metabolism and P storage. It remains likely that floral initiation in *M. bisulcata* takes place at the end of the wet season, before leaf exchange occurs in the dry season [see discussion in 11]. Earlier results on cultivated apple (*Malus*) trees suggested already that a two-way process of hormonal control could be operating [47], and the idea that sugars and other NSCs destined for the usually stronger root sink in mandarin (*Citrus*) [152], and in prune (*Prunus*) on K-deficient soils [153], might be diverted to the flowering meristem sinks under the control of gibberellins [154]. How exactly the ‘switch’ occurs remains elusive [48], and still to this day. The precise consistency of alternate bearing in pistachio (*Pistacea vera*) orchards has been questioned though, on statistical grounds [155].

### Active storage hypothesis

Storage of C as reserves in trees (stem rays, bark and branch wood) was widely seen in the past as being passive. The C was being made available for regular annual growth processes in ways that optimized the economy of the tree [156]. Extending this idea to nutrients, Chapin *et al.* [157] produced a systematic table for N and P, reserves of the latter mainly featuring as inorganic phosphate, polyphosphates, or phospholipids, leading them to conclude that dependence on mineral nutrients is likely much more pronounced than it is on C for reproduction. Sala *et al.* [158] argued that the build-up of NSCs in stores in trees might be acting as a buffer against stochastic variation in C inputs due to environmental stresses (e.g. drought, herbivore attacks), which as a bet-hedging strategy would insure against temporary collapses in maintenance and growth. The storage would need to be active, rather than passive, otherwise in the latter case it would be used up immediately in extra current growth. Extending this idea to mineral nutrients, Dietze *et al.* [159] hypothesized that N, P and K might also be accumulated and stored to guard against periods of reduced nutrient uptake, stabilizing the internal nutrient cycles. Hormones might therefore control source-sink processes actively. The empirical evidence for active vs. passive control of C storage in trees still remains weak and unresolved though [160]. But where P and K especially are limiting growth, evolution of such an active mechanism could be of considerable evolutionary advantage. Mast fruiting imposes a quasi-unpredictable (partially stochastic) demand on mineral nutrients supplies, and therefore for P and K active storage would serve it well, particularly in sites where uptake patterns themselves are subject to environmental stochasticity (the intensity of the dry season at Korup, for example).

### Nutrients in alternate bearing of tree crops

Studies of alternate-bearing pistachio trees have provided several valuable insights into how tree nutrient balance changes between ‘on’ and ‘off’ years. Whilst fruiting depresses root growth in fruiting years, uptake rates from the soil remained similar. If N, P and K moved from uptake mainly to leaves and shoots in ‘off’ years, and to fruits in ‘on’ ones at 3–6-fold the investment compared with ‘off’-year uptake, this implies that stores were being drawn upon [50]. Trees stored 7- and 14-fold more N and P respectively in ‘off’ than ‘on’ years, mostly in branches, which also suggests active storage [51]. Of what went into fruits in an ‘on’ year, 70% of N and P came from storage, and 30% from retranslocation. For N and P, storage forms are well known (e.g. phosphoproteins), but for K they are not; it would need to be intensively taken up in the ‘on’ years (study cited here was made on well-fertilized plots, however), despite the slower root growth, transported and used immediately in the fruits [161]. Interestingly, K concentrations in pistachio are inversely related to NSC concentrations, because K is needed for sugar transport to the apices where fruits are being formed [149]. If this last process applied to *M. bisulcata* at Korup, then K would be taken up in the mast years and, assuming also it cannot be stored in roots or branches, the only other feasible (temporary) store would be outside the plant, this as exchangeable forms held on soil colloids, built up in the wetter non-masting years (G. A. Neba, D. M. Newbery and G. B. Chuyong, unpubl. data). How far K might be transported laterally in the mat by mass flow is unknown.

Picchioni *et al.* [162]further showed, for pistachio again, that N and P were translocated out of the mature pericarp into the seeds of the same fruits, although K was not, which left the pericarp relative rich in K on fruit fall (harvesting). This could have been happening in *M. bisulcata*, and in this way explains the K-rich pods. Possibly as pods browned they too moved N and P to the seeds. With K having achieved its role in aiding sugar transport (under the hypothetical hormonal control suggested), it would then be redundant as regards reproductive investment *per se*: the fallen pods would be leached much later by the end wet season rains, even some remaining until next year, and in this way returning the extra K to the forest floor and soil colloid sites. Returning to the question of possible CHO storage in ‘off’ years, direct estimates showed that very little of the stores laid down in those ‘off’ years (8%) contributed to the ‘on’-year yield [163] – confirming again the general finding mentioned above in the *Introduction*. Overall, the main evidence is in favour of nutrients plus hormones being the controllers of alternate bearing and, in extension, masting in forest tree species where nutrients are limiting.

### Soil nutrient status

Nitrogen supply has been shown to be important for fruiting in *Nothofagus* [164]. In either N-fertilized stands, or unfertilized ones when rainfall was high (allowing higher N mineralization), masting was enhanced, subject to a temperature cue also operating. Likewise, Leeper *et al.* [165] showed in N-poor sites, that masting of spruce was less frequent and less intense than at N-rich sites. Cone production was positively correlated with soil N in the year prior, not the current one. Both studies found lagged N-effects, implying that N was being stored. Nevertheless, unravelling the causes is difficult because the better, wetter, conditions one year for N supply are not conducive to floral initiation, which depends more on warm drier ones affording higher sugar transport to apices. An inherent inverse correlation often exists which leads to an unstable alternation: a similar pattern is proposed for *M. bisulcata* at Korup with the relatively wetter and drier dry seasons.

Evidence for depletion of nutrients during masting, needing then restocking for the next event, is unfortunately limited. For white-bark pines, P and N declined in branches after masting – cones having especially high P concentrations [166]; and this could be linked to the species being ectomycorrhizal. Depletion spread over other plant stores according to the amount needed for masting. Not only do Sala *et al.* [166] provide evidence of resource switching (from storage to reproduction), but weather seems to be cuing for masting by synchronized nutrient uptake being made possible. This supports the PACER hypothesis and the main thesis of the current paper on *M. bisulcata*.

Climate factors which affect nutrient resource availability driving masting was a central argument of the review by Allen *et al.* [45]. This underlying process is how economies of scale become operational: the predisposition of trees to avoid seed predation or increasing their flowering efficacy is enhanced by synchronization. Allen *et al.* [45] made strong recognition of the role of ectomycorrhizas, building on Henkel *et al.* [9] and Newbery *et al.* [11]. That either N or P (rarely both) are often limiting where ectomycorrhizas tree species are successfully dominant in forests, must place a high demand on these elements for seed production. A set of resource acquisition and storage traits that would benefit growth and defense in these nutrient-poor habitats generally would be expected to be channel-selected through evolution to support an optimal life history strategy [45]. Different storage routes could be postulated to work in temperate vs. tropical systems where respectively N and P are more limiting. The Korup work posits that resource synchronization might, in part, be achieved via a shared mycorrhizal network. It is a working hypothesis that requires testing in the field at Korup. Given the lack of support for C storage being important for mast fruiting, the C budget model of Isagi *et al.* [19] needs to be rethought in terms P uptake, storage and usage.

Fernandez-Martinez *et al.* [15] recognized that many mast fruiting species grow in N-or P-poor soils, or at least on sites where the demand for these elements is high. The authors relied on foliar N and P concentration data. There are two problems with this approach: firstly, from among the masting species some samples may be from mast years, others from inter-mast ones pulling the differences between masting and non-masting species apart; and secondly, many species well adapted physiologically to nutrient poor soils do have nutrient levels similar to those on richer soils but display different nutrient dynamics. Again, the obvious association with, and role of, ectomycorrhizas was overlooked. Contrary to the claim of Kelly [17], nutrients can be hypothesized to determine masting when both their availability fluctuates, and their tree nutrient concentrations are synchronized via the mycorrhizal network. Stronger competition between masting and non-masting neighbours need not be invoked, as was proposed by Fernandez-Martinez *et al.* [16]. It follows from ecophysiological adaption of the masting species to their nutrient-poor conditions. From the nutrient-supply model the other economy of scale factors can operate; in the case of *M. bisulcata* this would probably be mainly to satiate seed predators [167].

Janzen [168] asked why the intervals between mast fruiting in dipterocarps are so large and irregular. That would not fit with resources like P accumulating to given common levels. Resources could doubtless be stored like NSCs, but not stored only for fruiting, and when not so used would be moved to other sinks such as roots. Dipterocarps form diverse mixtures of species in many forests, with their individuals spaced out among those of other ectomycorrhizal and non-ectomycorrhizal species. Thus, the functioning of a shared below­ground network seems less feasible, especially when the fungal species differ across tree species. The interestingly idea was raised, though, that masting may have originated in species when they were much more clumped in the evolutionary past than they are today, growing as patches of near single-species dominating stands[168]. If nutrient resources were more commonly distributed, shared and equilibrated within such patches in the manner postulated for extant *M. bisulcata*, and over time the tendency to clump by some species degenerated as they locally formed mixtures with other tree species, then the nutrient-threshold mechanism behind masting may have lost its effectiveness.

It is intriguing that for several dipterocarp species in Peninsular Malaysia hill forests, especially in the more seasonal N/NW states, mast fruiting occurred with an interval of 2-3 years (sometimes up to 5 years), showing some synchrony across individuals although often with no more than half the adult tree population participating in any one event [169]. Drought conditions almost always preceded masting. This is a similar situation to *M. bisulcata* at Korup but with a less pronounced pattern. Could the same hypothesis/model of P acquisition, storage and allocation in essence also explain mast fruiting of dipterocarps in these habitats [124, 127], with drought and possibly less strong nutrient limitation operating [64, 170]?

### Tree-tree root connections

Root grafts could potentially provide another means of nutrient transfer between *M. bisulcata* trees within groves. Soil surface grafts have been observed at Korup although none anatomically investigated and mapped so far. Early field experiments on *Pinus strobilus* in North America by Bormann [171] point to this as a real possibility. It was suggested that the process would mostly operate between large dominant trees of similar size to enable an equilibration of resources across trees. Many temperate and tropical tree species form root grafts [67, 172], often close to stems and among large roots, but there is also evidence for grafts between roots at greater distances when the species has long surface roots, e.g. for *Dacryodes excelsa* (Burseraceae) in Puerto Rica [173], or among smaller roots well below the surface, e.g. in oaks [174]. The extensively spreading root systems of *M. bisulcata*, connected to large buttresses [79], make the crossing of roots at 15–20 m away from stems quite feasible. Because grafts form as a result of physical pressing together of roots, the restricted vertical space of the root mat would likely also encourage grafts between medium to small roots, and in this way create a large multi-nodal root network. Apart from possibly providing increased resistance to wind throw – which would not be important for *M. bisulcata* given its strong buttressing – and temporarily supporting suppressed, damaged or dying neighbours, other advantages of root grafts have not been so obvious [175]. However, synchronizing nutrient supplies between, and storage within, trees across whole clusters would clearly be beneficial and strengthen masting. Root grafts would moreover provide a strong ‘platform’ for the ectomycorrhizal network and faster local transfers. A role for root grafts enabling masting has recently been suggested by Noble *et al.* [176] to partly explain alternate bearing in pistachio in a year when pollination synchrony (the Moran effect) was relatively weak. The focus in that plantation study was on C as the putative resource being transferred.

More directly relevant to *M. bisulcata* was the 1953 work of Yli-Vakkuri, cited by [171] showing that P was moving across root grafts of *Pinus sylvestris*: the author made no connection to ectomycorrhizal status of pines or association of ectomycorrhizas with mast fruiting [see 64]. Conceivably, ectomycorrhizal and grafted root-networks might operate together within *M. bisulcata* tree clusters, either contemporaneously or in successive phases around the end of the dry into the early wet season. If hormones are also transportable across grafts, an added level of synchronizing control is afforded. The role of tree root interconnections (ectomycorrhizas and grafts) becomes an essential auxiliary to the main masting hypothesis. Bormann [171] advanced the thesis that root grafts were established first by alignment of xylem tissues which rerouted water flow, and connecting phloem tissues permitted organic and nutrient transfer later. Between the mid- and late-ontogeny stages, *M. bisulcata* trees growing in dense patches become large adults, as described by the transient dominance model [75], and root grafting may have started when water was at a premium in the dry season. It then changed to having a more organic-flux transfer role once deep roots had secured all-year-round water supplies, the root mat had fully formed, the litter nutrient cycle was stable, and P (and possibly K) were by then in greater demand for growth and reproduction. To know whether or not codominants *Tetraberlinia bifoliolata* and *T. korupensis* graft below ground onto *M. bisulcata* would be interesting. That the three species do not usually mast synchronously suggests, though, that they probably do not form interspecific connections as predicted by Graham and Bormann [67].

### Role of environmental stochasticity

Dynamics simulation models involving storage and thresholds, resource-switching in a randomly varying environment, yet with no economy of scale operating, are able to generate masting cycles when either the variation is strong (i.e. very good vs poor years occur) or, when the variation is not so strong, there is an increasing non-linear response to the positive variations [177]. Even then masting can be diluted over successive years and is not quite bimodal. Under weaker variation in the environment an economy of scale is needed to achieve a well-defined masting cycle [69, 178]. Such large differences between very good and poor years in terms of resources (assuming linear responses) are ecologically rather unrealistic though for medium-length time series covering fairly regular 2- or 3-year masting cycles. This in turn suggests that without an economy of scale a non-linear response is more likely to be involved. An essential requirement of such masting would be to avoid degeneration of the time series into chaos – as shown by the models of Rees *et al.* [179].

The systems view would postulate feedback control to avoid such chaos, but it could allow for an unpredictable mix of 1-, 2- and 3-year cycles over time to the same effect in thwarting predator tracking. For a limited resource like P, such a system means regulation of internal fluctuations across the tree parts within a masting cycle, so that some equilibrium in the P dynamics is maintained medium-term. An economy of scale which itself involves a mechanism for P distribution would facilitate masting when based on storage and thresholds of that same resource. In this respect, ectomycorrhizal networks, which enable P acquisition and uptake (and possibly P storage), could bring about an equilibration of P across the connected roots of neighbouring trees, which would allow a contemporaneous storing and triggering of mast fruiting, as P levels rose in concert across all trees in the population (Fig. 5). In the case of *M. bisulcata* in this study, the key climate variables vary in a somewhat normal way, and it is unlikely they can cue masting in a simple directly linear manner. However, if wetter dry seasons lead to non-linear increases, or pulses, of P-and/or K-uptake, coordinated across the forest, synchrony is also possible. The network idea would also explain why one or both *Tetraberlinia* species often mast fruit along with *M. bisulcata*, because they largely share the same mycorrhizal species [73, 86]. The non-stationary aspect of the main hypothesis suggests a finely tuned evolved ecophysiological system in *M. bisulcata* allowing it to respond to variation of the climate to maximize its fitness.

## CONCLUSIONS

The results in this paper confirm, update and extend the findings based on those for 1989–2004 presented in Newbery *et al.* [11] with greater statistical detail and confidence. It would have been valuable to know whether the 3-year cycle returned after 2017. Post-dispersal predation is currently important, especially in well-lighted gaps although not in the shaded understorey or below *M. bisulcata* crowns. It could have been much more common in the past when either *M. bisulcata* individuals were scattered across the landscape [74], or when grove dynamics was at the rebuilding stage [75]. Since, seeds and seedlings need P to establish and reach to the network of ectomycorrhizas, it would be evolutionarily advantageous for an ectomycorrhizal species to mast fruit and increase its chances of establishment. It is interesting that masting is achieved in the most clustered detarioids and was not apparent for the more spatially dispersed species in southern Korup.

Most of the C dynamics models for trees to date [e.g. 180, 181-183] do not cater adequately, if at all, for environmental stochasticity or non-constant reproductive allocation schedules. Source-sink models for herbaceous plants, let alone for more complicated trees, have made little progress in the last three decades since the work of Farrar and others [184–188], apart from raising an unresolved debate over passive versus active storage. They usually assume steady-state equilibrium conditions, are often teleonomic [e.g. 189], and are strongly focussed on stem diameter growth and maximizing tree yield. They are, therefore, still too far away from an understanding of trees in natural forest ecosystems [190] to explain even reproductive schedules like masting. Source-sink relationships, within trees especially, turns out to be very complicated: a more integrated systems approach is probably required. A considerable challenge is working out how P, K and N interact with C dynamics over the complete life-history schedule of a tree. A very interesting hypothesis coming from this extended phenological analysis is that both K and P levels might synergistically control mast fruiting of *M. bisulcata*, mediated by the physical structure and functioning of the surface root mat for nutrient cycling. To test this hypothesis further requires much more detailed recording of nutrient processes at the tree-level and within the system, replicated over several mast fruiting events.

The literature review of the likely mechanisms lying behind mast fruiting in *M. bisulcata* gives valuable pointers to detailed aspects that need investigation. The most critical data for the nutrient resource limitation hypothesis would be phosphorus concentrations in roots, branches, leaves and fruits of a large sample of trees over several masting cycles. Detection of phytic acid in branch wood, prior to fruiting especially, could confirm that P was being stored. Lack of predicted changes in P over time would falsify the hypothesis. In addition, to test the related economy-of-scale mechanism for equilibration of P between adult trees at masting would require showing everywhere a flux of P through the mycelia of the shared ectomycorrhizal network, i.e. between neighbouring large trees.

The fundamental driving variable behind the deterministic mechanically intricate system operating to cause mast fruiting in *M. bisulcata*, is the stochastic variation in rainfall (Fig. 5). The vicissitudes of rainfall in the dry season over successive years could generate the non-linear differences needed to enable the storage process. Stochasticity becomes the ultimate ‘causal’ factor over evolutionary time. This can operate at Korup because P as a resource is strongly limiting the system, there is seasonality, and rainfall alters access, availability and uptake of P. The hypothesis argues for a fruiting-rooting trade-off as the basis to alternate bearing and masting. For a common nutrient threshold to operate at its strongest trees must be spatially close and connected via their roots and mycorrhizas. The new evidence and analyses here, showing that both climate and ectomycorrhizas might operate respectively on and within the forest nutrient cycle, makes for a major step forward in refining the original PACER hypothesis [81]. The life history strategy of a tree species, such as *M. bisulcata*, can only be understood within the frame of all the other processes operating in the ecosystem where it occurs.

## Supporting information

Suppementary Electronic Materials

## ACKNOWLEDGEMENTS

The Ministries of Forests and Wildlife (MINFOF) and Scientific Research and Innovation (MINRESI) granted permission to conduct field research, and the Conservator’s Office of Korup National Park approved access to the field site. We thank S. Ngibili (phenological recording) and M. Zimmermann (chemical analysis of plant and soil materials) for their valuable technical assistance, X. M. van der Burgt for help with the soil moisture readings, and reviewer and editor for their valuable comments and suggestions.

## Supplementary materials

Suppl. material 1. Radiometry at Bulu, Ndian: calibration and interpretation.

Suppl. material 2. Evidence table for the levels of mast fruiting per year.

Suppl. material 3. Time series analysis of dry season climate variables.

Suppl. material 4. Regression model fits of masting on rainfall and radiation.

Suppl. material 5. Rainfall and 30-day rainfall totals defining dry seasons.

Suppl. material 6. Description and feeding of the yellow-green caterpillar.

Suppl. material 7. Analysis of Ndian PAMOL oil-palm plantation yields.

Suppl. material 8. Stem increments from tree stem cores.

Suppl. material 9. Minimum temperature and mast fruiting.

## Notes

https://doi.org/10.5061/dryad.r7sqv9shz

### Competing Interest Statement

The authors have declared no competing interest.

### Summary of Updates

1. A new analysis of the effect of dry season minimum temperature on mast fruiting has been incorporated. This involves new paragraphs in each on the main sections, plus a new Appendix 9 (ESM). 2. A text box "Synopsis of the Results" is now included.

### Funding

- Swiss National Science Foundation: 3100-066655

## REFERENCES

1 Maynard Smith, J. 1972 On evolution. Edinburgh, UK: Edinburgh University Press.

2 Charlesworth, B. 1980 Evolution in age-structured populations. Cambridge, UK: Cambridge University Press.

3 Stearns, S. C. 1992 The evolution of life-history strategies. Oxford, UK: Oxford University Press.

4 Norton, D. A., Kelly, D. 1988 Mast seeding over 33 years by *Dacrydium cupressinum* Lamb. (rimu) (Podocarpaceae) in New Zealand: the importance of economies of scale. Functional Ecology. 2, 399–408. (DOI10.2307/2389413)

5 Kelly, D. 1994 The evolutionary ecology of mast seeding. Trends in Ecology & Evolution. 9, 465–470. (10.1016/0169-5347(94)90310-7)

6 Kelly, D., Sork, V. L. 2002 Mast seeding in perennial plants: Why, how, where? Annual Review of Ecology and Systematics. 33, 427–447. (10.1146/annurev.ecolsys.33.020602.095433)

7 Crone, E. E., Rapp, J. M. 2014 Resource depletion, pollen coupling, and the ecology of mast seeding. Annals of the New York Academy of Sciences. 1322, 21–34. (10.1111/nyas.12465)

8 Pearse, I. S., Koenig, W. D., Kelly, D. 2016 Mechanisms of mast seeding: resources, weather, cues, and selection. New Phytologist. 212, 546–562. (10.1111/nph.14114)

9 Henkel, T. W., Mayor, J. R., Woolley, L. P. 2005 Mast fruiting and seedling survival of the ectomycorrhizal, monodominant *Dicymbe corymbosa* (Caesalpiniaceae) in Guyana. New Phytologist. 167, 543–556. (10.1111/j.1469-8137.2005.01431.x)

10 Ichie, T., Kenta, T., Nakagawa, M., Sato, K., Nakashizuka, T. 2005a Resource allocation to reproductive organs during masting in the tropical emergent tree, *Dipterocarpus tempehes*. Journal of Tropical Ecology. 21, 237–241. (10.1017/s0266467404002214)

11 Newbery, D. M., Chuyong, G. B., Zimmermann, L. 2006a Mast fruiting of large ectomycorrhizal African rain forest trees: importance of dry season intensity, and the resource-limitation hypothesis. New Phytologist. 170, 561–579. (10.1111/j.1469-8137.2006.01691.x)

12 Norden, N., Chave, J., Belbenoit, P., Caubère, A., Châtelet, P., Forget, P.-M., Thébaud, C. 2007 Mast fruiting is a frequent strategy in woody species of eastern South America. PlosOne. 10, 1–9 (e1079). (10.1371/journal.pone.0001079)

13 Laland, K. N., Sterelny, K., Odling-Smee, J., Hoppitt, W., Uller, T. 2011 Cause and effect in biology revisited: Is Mayr’s proximate-ultimate dichotomy still useful? Science. 334, 1512–1516. (10.1126/science.1210879)

14 Corning, P. A. 2019 Teleonomy and the proximate-ultimate distinction revisited. Biological Journal of the Linnean Society. 127, 912–916.

15 Fernandez-Martinez, M., Pearse, I., Sardans, J., Sayol, F., Koenig, W. D., LaMontagne, J. M., Bogdziewicz, M., Collalti, A., Hacket-Pain, A., Vacchiano, G., et al. 2019 Nutrient scarcity as a selective pressure for mast seeding. Nature Plants. 5, 1222–1227. (10.1038/s41477-019-0549-y)

16 Fernandez-Martinez, M., Sardans, J., Sayol, F., LaMontagne, J. M., Bogdziewicz, M., Collalti, A., Hacket-Pain, A., Vacchiano, G., Espelta, J. M., Penuelas, J., et al. 2020 Reply to: Nutrient scarcity cannot cause mast seeding. Nature Plants. 6, 763–765. (10.1038/s41477-020-0703-6)

17 Kelly, D. 2020 Nutrient scarcity cannot cause mast seeding. Nature Plants. 6, 760–762. (10.1038/s41477-020-0702-7)

18 Lloyd, D. G. 1980 Sexual strategies in plants. I. An hypothesis of serial adjustment of maternal investment during one reproductive season. New Phytologist. 86, 69–79. (10.1111/j.1469-8137.1980.tb00780.x)

19 Isagi, Y., Sugimura, K., Sumida, A., Ito, H. 1997 How does masting happen and synchronize? Journal of Theoretical Biology. 187, 231–239. (10.1006/jtbi.1997.0442)

20 Abe, T., Tachiki, Y., Kon, H., Nagasaka, A., Onodera, K., Minamino, K., Han, Q. M., Satake, A. 2016 Parameterisation and validation of a resource budget model for masting using spatiotemporal flowering data of individual trees. Ecology Letters. 19, 1129–1139. (10.1111/ele.12651)

21 Satake, A., Bjornstad, O. N. 2008 A resource budget model to explain intraspecific variation in mast reproductive dynamics. Ecological Research. 23, 3–10. (10.1007/s11284-007-0397-5)

22 Vacchiano, G., Ascoli, D., Berzaghi, F., Lucas-Borja, M. E., Caignard, T., Collalti, A., Mairota, P., Palaghianu, C., Reyer, C. P. O., Sanders, T. G. M., et al. 2018 Reproducing reproduction: How to simulate mast seeding in forest models. Ecological Modelling. 376, 40–53. (10.1016/j.ecolmodel.2018.03.004)

23 Venner, S., Siberchicot, A., Pelisson, P. F., Schermer, E., Bel-Venner, M. C., Nicolas, M., Debias, F., Miele, V., Sauzet, S., Boulanger, V., et al. 2016 Fruiting strategies of perennial plants: a resource budget model to couple mast seeding to pollination efficiency and resource allocation strategies. American Naturalist. 188, 66–75. (10.1086/686684)

24 Yamauchi, A. 1996 Theory of mast reproduction in plants: Storage-size dependent strategy. Evolution. 50, 1795–1807. (10.2307/2410737)

25 Ye, X. J., Sakai, K. 2016 A new modified resource budget model for nonlinear dynamics in citrus production. Chaos Solitons & Fractals. 87, 51–60. (10.1016/j.chaos.2016.03.016)

26 Marshall, C., Grace, J., eds. 1992 Fruit and seed production: aspects of development, environmental physiology and ecology. Cambridge, UK: Cambridge University Press.

27 Bustan, A., Avni, A., Lavee, S., Zipori, I., Yeselson, Y., Schaffer, A. A., Riov, J., Dag, A. 2011 Role of carbohydrate reserves in yield production of intensively cultivated oil olive (*Olea europaea* L.) trees. Tree Physiology. 31, 519–530. (10.1093/treephys/tpr036)

28 Hoch, G. 2005 Fruit-bearing branchlets are carbon autonomous in mature broad-leaved temperate forest trees. Plant Cell and Environment. 28, 651–659. (10.1111/j.1365-3040.2004.01311.x)

29 Hoch, G., Richter, A., Körner, C. 2003 Non-structural carbon compounds in temperate forest trees. Plant, Cell and Environment. 26, 1067–1081. (10.1046/j.0016-8025.2003.01032.x)

30 Ichie, T., Igarashi, S., Yoshida, S., Kenzo, T., Masaki, T., Tayasu, I. 2013 Are stored carbohydrates necessary for seed production in temperate deciduous trees? Journal of Ecology. 101, 525–531. (10.1111/1365-2745.12038)

31 Igarashi, S., Shibata, M., Masaki, T., Tayasu, I., Ichie, T. 2019 Mass flowering of *Fagus crenata* does not depend on the amount of stored carbohydrates in trees. Trees-Structure and Function. 33, 1399–1408. (10.1007/s00468-019-01867-w)

32 Millard, P., Sommerkorn, M., Grelet, G. A. 2007 Environmental change and carbon limitation in trees: a biochemical, ecophysiological and ecosystem appraisal. New Phytologist. 175, 11–28. (10.1111/j.1469-8137.2007.02079.x)

33 Miyazaki, Y. 2013 Dynamics of internal carbon resources during masting behavior in trees. Ecological Research. 28, 143–150. (10.1007/s11284-011-0892-6)

34 Mund, M., Kutsch, W. L., Wirth, C., Kahl, T., Knohl, A., Skomarkova, M. V., Schulze, E. D. 2010 The influence of climate and fructification on the inter-annual variability of stem growth and net primary productivity in an old-growth, mixed beech forest. Tree Physiology. 30, 689–704. (10.1093/treephys/tpq027)

35 Mund, M., Herbst, M., Knohl, A., Matthäus, B., Schumacher, J., Schall, P., Siebicke, L., Tamrakar, R., Ammer, C. 2020 It is not just a ‘trade-off’: indications for sink- and source-limitation to vegetative and regenerative growth in an old-growth beech forest. New Phytologist. 226, 111–125. (10.1111/nph.16408)

36 Richardson, A. D., Carbone, M. S., Keenan, T. F., Czimczik, C. I., Hollinger, D. Y., Murakami, P., Schaberg, P. G., Xu, X. M. 2013 Seasonal dynamics and age of stemwood nonstructural carbohydrates in temperate forest trees. New Phytologist. 197, 850–861. (10.1111/nph.12042)

37 Sanchez-Humanes, B., Sork, V. L., Espelta, J. M. 2011 Trade-offs between vegetative growth and acorn production in *Quercus lobata* during a mast year: the relevance of crop size and hierarchical level within the canopy. Oecologia. 166, 101–110. (10.1007/s00442-010-1819-6)

38 Wurth, M. K. R., Pelaez-Riedl, S., Wright, S. J., Korner, C. 2005 Non-structural carbohydrate pools in a tropical forest. Oecologia. 143, 11–24. (10.1007/s00442-004-1773-2)

39 Igarashi, S., Yoshida, S., Kenzo, T., Sakai, S., Nagamasu, H., Hyodo, F., Tayasu, I., Mohamad, M., Ichie, T. 2024 No evidence of carbon storage usage for seed production in 18 dipterocarp masting species in a tropical rain forest. Oecologia. (10.1007/s00442-024-05527-w)

40 Bogdziewicz, M., Kelly, D., Ascoli, D., Caignard, T., Chianucci, F., Crone, E. E., Fleurot, E., Foest, J. J., Gratzer, G., Hagiwara, T., et al. 2024 Evolutionary ecology of masting: mechanisms, models, and climate change. Trends in Ecology & Evolution. 39, 851–862. (10.1016/j.tree.2024.05.006)

41 Han, Q. M., Kabeya, D. 2017 Recent developments in understanding mast seeding in relation to dynamics of carbon and nitrogen resources in temperate trees. Ecological Research. 32, 771–778. (10.1007/s11284-017-1494-8)

42 Han, Q. M., Kabeya, D., Iio, A., Inagaki, Y., Kakubari, Y. 2014 Nitrogen storage dynamics are affected by masting events in *Fagus crenata*. Oecologia. 174, 679–687. (10.1007/s00442-013-2824-3)

43 Fernandez-Martinez, M., Vicca, S., Janssens, I. A., Espelta, J. M., Penuelas, J. 2017b The role of nutrients, productivity and climate in determining tree fruit production in European forests. New Phytologist. 213, 669–679. (10.1111/nph.14193)

44 Le Roncé, I., Dardevet, E., Venner, S., Schönbeck, L., Gessler, A., Chuine, I., Limousin, J. M. 2023 Reproduction alternation in trees: testing the resource depletion hypothesis using experimental fruit removal in *Quercus ilex*. Tree Physiology. 43, 952–964. (10.1093/treephys/tpad025)

45 Allen, R. B., Millard, P., Richardson, S. J. 2017 A resource centric view of climate and mast seeding in trees. In Progress in Botany. Vol. 79. (ed.^eds. F. Canovas, U. Lüttige, R. Matyssek), pp. 233–268. Cham, Switzerland: Springer.

46 Crawley, M. J., Long, C. R. 1995 Alternate bearing, predator satiation and seedling recruitment in *Quercus robur* L. Journal of Ecology. 83, 683–696. (10.2307/2261636)

47 Davis, L. D. 1957 Flowering and alternate bearing. American Society for Horticultural Science. 70, 545–556.

48 Monselise, S. P., Goldschmidt, E. E. 1982 Alternate bearing in fruit trees. Horticultural Research. 4, 128–173. (10.1002/9781118060773.ch5)

49 Garcia, G., Re, B., Orians, C., Crone, E. 2021 By wind or wing: pollination syndromes and alternate bearing in horticultural systems. Philosophical Transactions of the Royal Society B-Biological Sciences. 376, 1–9. (10.1098/rstb.2020.0371)

50 Rosecrance, R. C., Weinbaum, S. A., Brown, P. H. 1996 Assessment of nitrogen, phosphorus, and potassium uptake capacity and root growth in mature alternate-bearing pistachio (*Pistacia vera*) trees. Tree Physiology. 16, 949–956. (10.1093/treephys/16.11-12.949)

51 Rosecrance, R. C., Weinbaum, S. A., Brown, P. H. 1998 Alternate bearing affects nitrogen, phosphorus, potassium and starch storage pools in mature pistachio trees. Annals of Botany. 82, 463–470. (10.1006/anbo.1998.0696)

52 Ashton, P. S., Givnish, T. J., Appanah, S. 1988 Staggered flowering in the Dipterocarpaceae: new insights into floral induction and the evolution of mast fruiting in the aseasonal tropics. American Naturalist. 132, 44–66.

53 Sakai, S. 2002 General flowering in lowland mixed dipterocarp forests of South-east Asia. Biological Journal of the Linnean Society. 75, 233–247.

54 Sakai, S., Harrison, R. D., Momose, K., Kuraji, K., Nagamasu, H., Yasunari, T., Chong, L., Nakashizuka, T. 2006 Irregular droughts trigger mass flowering in aseasonal tropical forests in Asia. American Journal of Botany. 93, 1134–1139.

55 Chen, Y. Y., Satake, A., Sun, I. F., Kosugi, Y., Tani, M., Numata, S., Hubbell, S. P., Fletcher, C., Supardi, M. N. N., Wright, S. J. 2018 Species-specific flowering cues among general flowering Shorea species at the Pasoh Research Forest, Malaysia. Journal of Ecology. 106, 586–598. (10.1111/1365-2745.12836)

56 Chechina, M., Hamann, A. 2019 Climatic drivers of dipterocarp mass-flowering in South-East Asia. Journal of Tropical Ecology. 35, 108–117. (10.1017/s0266467419000087)

57 Brearley, F. Q., Proctor, J., Surinata, Nagy, L., Dalrymple, G., Voysey, B. C. 2007 Reproductive phenology over a 10-year period in a lowland evergreen rain forest of central Borneo. Journal of Ecology. 95, 828–839.

58 Kurten, E. L., Bunyavejchewin, S., Davies, S. J. 2018 Phenology of a dipterocarp forest with seasonal drought: Insights into the origin of general flowering. Journal of Ecology. 106, 126–136. (10.1111/1365-2745.12858)

59 Tutin, C. E. G., Fernandez, M. 1993 Relationships between minimum temperature and fruit production in some tropical forest trees in Gabon. Journal of Tropical Ecology. 9, 241–248.

60 Chapman, C. A., Wrangham, R. W., Chapman, L. J., Kennard, D. K., Zanne, A. E. 1999 Fruit and flower phenology at two sites in Kibale National Park, Uganda. Journal of Tropical Ecology. 15, 189–211.

61 Alexander, I. 1989a Mycorrhizas in tropical forests. In Mineral nutrients in tropical forest and savanna ecosystems. (ed.^eds. J. Proctor), pp. 169–188. Oxford: Blackwell Scientific Publications.

62 Alexander, I. J. 1989b Systematics and ecology of ectomycorrhizal legumes. Monographs in Systematic Botany of the Missouri Botanical Garden. 29, 607–624.

63 Brearley, F. Q. 2012 Ectomycorrhizal associations of the Dipterocarpaceae. Biotropica. 44, 637–648. (10.1111/j.1744-7429.2012.00862.x)

64 Newbery, D. M. 2005 Ectomycorrhizas and mast fruiting in trees: linked by climate-driven tree resources? New Phytologist. 167, 324–326. (10.1111/j.1469-8137.2005.01468.x)

65 Corrales, A., Henkel, T. W., Smith, M. E. 2018 Ectomycorrhizal associations in the tropics - biogeography, diversity patterns and ecosystem roles. New Phytologist. 220, 1076–1091. (10.1111/nph.15151)

66 Simard, S. W., Perry, D. A., Jones, M. D., Myrold, D. D., Durall, D. M., Molina, R. 1997 Net transfer of carbon between ectomycorrhizal tree species in the field. Nature. 388, 579–582.

67 Graham, B. F., Bormann, F. H. 1966 Natural root grafts. Botanical Review. 32, 255–292. (10.1007/bf02858662)

68 Newbery, D. M., Songwe, N. S., Chuyong, G. B. 1998 Phenology and dynamics of an African rainforest at Korup, Cameroon. In Dynamics of tropical communities. (ed.^eds. D. M. Newbery, H. H. T. Prins, N. D. Brown), pp. 267–308: Blackwell Science.

69 Kerkhoff, A. J. 2004 Expectation, explanation and masting. Evolutionary Ecology Research. 6, 1003–1020.

70 Letouzey, R. 1968 Etude Phytogéographique du Cameroun. Paris: P. LeChevalier.

71 Gartlan, J. S. 1992 Cameroon. In The Conservation Atlas of Tropical Forests: Africa. (ed.^eds. J. A. Sayer, C. S. Harcourt, N. M. Collins), pp. 110–118. London, UK: Macmillan Publishers.

72 Gartlan, J. S., Newbery, D. M., Thomas, D. W., Waterman, P. G. 1986 The influence of topography and soil phosphorus on the vegetation of Korup Forest Reserve, Cameroun. Vegetatio. 65, 131–148. (10.1007/bf00044814)

73 Newbery, D. M., Alexander, I. J., Thomas, D. W., Gartlan, J. S. 1988 Ectomycorrhizal rain-forest legumes and soil phosphorus in Korup National Park, Cameroon. New Phytologist. 109, 433–450. (10.1111/j.1469-8137.1988.tb03719.x)

74 Newbery, D. M., Gartlan, J. S. 1996 Structural analysis of the rain forest at Korup and Douala Edea, Cameroon. Proceedings of the Royal Society of Edinburgh B. 104, 177–224. (10.1017/S0269727000006138)

75 Newbery, D. M., van der Burgt, X. M., Worbes, M., Chuyong, G. B. 2013 Transient dominance in a central African rain forest. Ecological Monographs. 83, 339–382. (10.1890/12-1699.1)

76 Newbery, D. M., van der Burgt, X. M., Moravie, M. A. 2004 Structure and inferred dynamics of a large grove of *Microberlinia bisulcata* trees in central African rain forest: the possible role of periods of multiple disturbance events. Journal of Tropical Ecology. 20, 131–143. (10.1017/s0266467403001111)

77 LPWG. 2017 A new subfamily classification of the Leguminosae based on a taxonomically comprehensive phylogeny. Taxon. 66, 44–77. (10.12705/661.3)

78 de la Estrella, M., Forest, F., Klitgard, B., Lewis, G. P., Mackinder, B. A., de Queiroz, L. P., Wieringa, J. J., Bruneau, A. 2018 A new phylogeny-based tribal classification of subfamily Detarioideae, an early branching clade of florally diverse tropical arborescent legumes. Scientific Reports. 8, 1–14. (10.1038/s41598-018-24687-3)

79 Newbery, D. M., Schwan, S., Chuyong, G. B., van der Burgt, X. M. 2009 Buttress form of the central African rain forest tree *Microberlinia bisulcata*, and its possible role in nutrient acquisition. Trees-Structure and Function. 23, 219–234. (10.1007/s00468-008-0270-3)

80 Norghauer, J. M., Newbery, D. M. 2015 Tree size and fecundity influence ballistic seed dispersal of two dominant mast-fruiting species in a tropical rain forest. Forest Ecology and Management. 338, 100–113. (10.1016/j.foreco.2014.11.005)

81 Newbery, D. M., Alexander, I. J., Rother, J. A. 1997 Phosphorus dynamics in a lowland African rain forest: The influence of ectomycorrhizal trees. Ecological Monographs. 67, 367–409. (10.1890/0012-9615(1997)067[0367:Pdiala]2.0.Co;2)

82 Chuyong, G. B., Newbery, D. M., Songwe, N. C. 2000 Litter nutrients and retranslocation in a central African rain forest dominated by ectomycorrhizal trees. New Phytologist. 148, 493–510. (10.1046/j.1469-8137.2000.00774.x)

83 Chuyong, G. B., Newbery, D. M., Songwe, N. C. 2002 Litter breakdown and mineralization in a central African rain forest dominated by ectomycorrhizal trees. Biogeochemistry. 61, 73–94. (10.1023/a:1020276430119)

84 Chuyong, G. B., Newbery, D. M., Songwe, N. C. 2004 Rainfall input, throughfall and stemflow of nutrients in a central African rain forest dominated by ectomycorrhizal trees. Biogeochemistry. 67, 73–91. (10.1023/B:BIOG.0000015316.90198.cf)

85 Newbery, D. M., Chuyong, G. B., Green, J. J., Songwe, N. C., Tchuenteu, F., Zimmermann, L. 2002 Does low phosphorus supply limit seedling establishment and tree growth in groves of ectomycorrhizal trees in a central African rainforest? New Phytologist. 156, 297–311. (10.1046/j.1469-8137.2002.00505.x)

86 Newbery, D. M., Alexander, I. J., Rother, J. A. 2000 Does proximity to conspecific adults influence the establishment of ectomycorrhizal trees in rain forest? New Phytologist. 147, 401–409. (10.1046/j.1469-8137.2000.00698.x)

87 Newbery, D. M., Chuyong, G. B., Zimmermann, L., Praz, C. 2006b Seedling survival and growth of three ectomycorrhizal caesalpiniaceous tree species in a Central African rain forest. Journal of Tropical Ecology. 22, 499–511. (10.1017/s0266467406003427)

88 Newbery, D. M., Praz, C. J., van der Burgt, X. M., Norghauer, J. M., Chuyong, G. B. 2010 Recruitment dynamics of the grove-dominant tree *Microberlinia bisulcata* in African rain forest: extending the light response versus adult longevity trade-off concept. Plant Ecology. 206, 151–172. (10.1007/s11258-009-9631-2)

89 Norghauer, J. M., Newbery, D. M. 2010 Recruitment limitation after mast-seeding in two African rain forest trees. Ecology. 91, 2303–2312. (10.1890/09-0071.1)

90 Green, J. J., Newbery, D. M. 2001a Light and seed size affect establishment of grove-forming ectomycorrhizal rain forest tree species. New Phytologist. 151, 271–289. (10.1046/j.1469-8137.2001.00156.x)

91 Green, J. J., Newbery, D. M. 2001b Shade and leaf loss affect establishment of grove-forming ectomycorrhizal rain forest tree species. New Phytologist. 151, 291–309. (10.1046/j.1469-8137.2001.00157.x)

92 Simard, S. W., Beiler, K. J., Bingham, M. A., Deslippe, J. R., Philip, L. J., Teste, F. P. 2012 Mycorrhizal networks: mechanisms, ecology and modelling. Fungal Biology Reviews. 26, 39–60.

93 Simard, S. W., Durall, D. M. 2004 Mycorrhizal networks: a review of their extent, function, and importance. Canadian Journal of Botany. 82, 1140–1165. (10.1139/b04-116)

94 Karst, J., Jones, M. D., Hoeksema, J. D. 2023 Positive citation bias and overinterpreted results lead to misinformation on common mycorrhizal networks in forests. Nature Ecology & Evolution. 7, 501-+. (10.1038/s41559-023-01986-1)

95 Green, J. J., Newbery, D. M. 2002 Reproductive investment and seedling survival of the mast-fruiting rain forest tree, Microberlinia bisulcata A. Chev. Plant Ecology. 162, 169–183. (10.1023/a:1020304212118)

96 Zimmermann, L. 2000 A comparative study of growth and mortality of trees in a caesalp dominated lowland African rainforest at Korup, Cameroon: Diploma (MSc) thesis: University of Bern, Switzerland.

97 Etta, C. E., Chuyong, G. B., Newbery, D. M. 2022 Climate records for Bulu, Ndian Division, SW Cameroon. [Data set]. https://zenodo.org/records/7220319.

98 Schwan, S. 2003 Phenology, resource conservation and tree architecture of large ectomycorrhizal trees in a lowland African rain forest at Korup, Cameroon. Diploma (MSc) thesis: University of Bern, Switzerland.

99 Newbery, D. M., Neba, G. A. 2019 Micronutrients may influence the efficacy of ectomycorrhizas to support tree seedlings in a lowland African rain forest. Ecosphere. 10, 1–28. (10.1002/ecs2.2686)

100 Agresti, A. 2007 An introduction to categorical data analysis. 2nd ed. New Jersey, USA: J. Wiley & Sons.

101 McCullagh, P., Nelder, J. A. 1989 Generalized linear models. Second ed. London, UK: Chapman & Hall.

102 Cox, D. R. 1981 Statistical analysis of time series: some recent developments. Scandinavian Journal of Statistics. 8, 93–115.

103 Fox, J. 2008 Applied regression analysis and generalized linear models. 2nd ed. Los Angeles, USA: Sage Publications.

104 Hosmer, D. W., Lemeshow, S., Sturdivant, R. X. 2013 Applied logistic regression. New Jersey, USA: J. Wiley & Sons.

105 Fox, J., Weisberg, S. 2011 An R companion to applied regression. 2nd ed. Los Angeles, CA.: Sage Publications.

106 Venables, W. N., Ripley, B. D. 2002 Modern applied statistics with S. 4th ed. Berlin: Springer.

107 R_Core_Team. 2022 R: A language and environment for statistical computing. R Foundation for Statistical Computing, Vienna, Austria. *URL* https://www.R-project.org/.

108 VSN_International. 2022 The guide to the Genstat Command Language (Release 22), Part 1 Syntax. Part 2 Statistics. VSN International, Hemel Hempstead, UK.

109 Dunsmuir, W. T. M., Scott, D. J. 2015 The glarma package for observation-driven time series regression of counts. Journal of Statistical Software. 67, 1–36.

110 Cox, D. R. 1958 The regression analysis of binary sequences. Journal of the Royal Statistical Society Series B-Statistical Methodology. 20, 215–242. (10.1111/j.2517-6161.1958.tb00292.x)

111 Cox, D. R., Snell, E. J. 1989 Analysis of binary data. 2nd. ed: Chapman & Hall/CRC.

112 Dunsmuir, W. T. M. 2016 Generalized linear autoregressive moving average models. In Handbook of discrete-valued time series. (ed.^eds. R. A. David, S. H. Holan, R. Lund, N. Ravishanker), pp. 51–76. Boca Raton, USA: CRC Press.

113 McKenzie, E. 1985 Some simple models for discrete variate time series. Water Resources Bulletin. 21, 645–650. (10.1111/j.1752-1688.1985.tb05379.x)

114 McKenzie, E. 2003 Discrete variate time series. In Handbook of Statistics. (ed.^eds. C. R. Rao, D. N. Shanbhag), pp. 573–606: Elsevier.

115 Norghauer, J. M., Newbery, D. M., Neba, G. A. 2023 Indications of an *Achaea* sp. caterpillar outbreak disrupting fruiting of an ectomycorrhizal tropical tree in Central African rainforest. Plant Ecology and Evolution. 156, 46–58. (10.5091/plecevo.96572)

116 Nicholson, S. E., Funk, C., Fink, A. H. 2018 Rainfall over the African continent from the 19th through the 21st century. Global and Planetary Change. 165, 114–127. (10.1016/j.gloplacha.2017.12.014)

117 Nicholson, S. E., Some, B., Kone, B. 2000 An analysis of recent rainfall conditions in West Africa, including the rainy seasons of the 1997 El Nino and the 1998 La Nina years. Journal of Climate. 13, 2628–2640. (10.1175/1520-0442(2000)013<2628:Aaorrc>2.0.Co;2)

118 Chuyong, G. B. 1994 Nutrient cycling in ectomycorrhizal legume-dominated forest in Korup National Park, Cameroon. PhD thesis: Stirling University, UK.

119 Fisher, R. A., Williams, M., Ruivo, M. D., de Costa, A. L., Meira, P. 2008 Evaluating climatic and soil water controls on evapotranspiration at two Amazonian rainforest sites. Agricultural and Forest Meteorology. 148, 850–861. (10.1016/j.agrformet.2007.12.001)

120 Or, D., Wraith, J. M. 2002 Soil water content and water potential relationships. In Soil physics companion. (ed.^eds. A. W.o Warrick), pp. 49–84. Boca Raton, FL, USA: CRC Press.

121 Corley, R. H. V., Tinker, P. B. 2016 The oil palm. 5th ed: J. Wiley.

122 Legros, S., Mialet-Serra, I., Clement-Vidal, A., Caliman, J. P., Siregar, F. A., Fabre, D., Dingkuhn, M. 2009 Role of transitory carbon reserves during adjustment to climate variability and source-sink imbalances in oil palm (*Elaeis guineensis*). Tree Physiology. 29, 1199–1211. (10.1093/treephys/tpp057)

123 Ichie, T., Kenzo, T., Kitahashi, Y., Koike, T., Nakashizuka, T. 2005b How does *Dryobalanops aromatica* supply carbohydrate resources for reproduction in a masting year? Trees-Structure and Function. 19, 703–710. (10.1007/s00468-005-0434-3)

124 Ichie, T., Nakagawa, M. 2013 Dynamics of mineral nutrient storage for mast reproduction in the tropical emergent tree *Dryobalanops aromatica*. Ecological Research. 28, 151–158. (10.1007/s11284-011-0836-1)

125 Aoyagi, R., Imai, N., Hidaka, A., Samejima, H., Kitayama, K. 2018 Abrupt increase in phosphorus and potassium fluxes during a masting event in a Bornean tropical forest. Ecological Research. 33, 1193–1205. (10.1007/s11284-018-1642-9)

126 Yeoh, S. H., Satake, A., Numata, S., Ichie, T., Lee, S. L., Basherudin, N., Muhammad, N., Kondo, T., Otani, T., Hashim, M., et al. 2017 Unravelling proximate cues of mass flowering in the tropical forests of South-East Asia from gene expression analyses. Molecular Ecology. 26, 5074–5085. (10.1111/mec.14257)

127 Kitayama, K., Tsujii, Y., Aoyagi, R., Aiba, S. 2015 Long-term C, N and P allocation to reproduction in Bornean tropical rain forests. Journal of Ecology. 103, 606–615. (10.1111/1365-2745.12379)

128 Norghauer, J. M., Newbery, D. M. 2016 Density-dependent dynamics of a dominant rain forest tree change with juvenile stage and time of masting. Oecologia. 181, 207–223. (10.1007/s00442-015-3534-9)

129 Bennett, E. J., Roberts, J. A., Wagstaff, C. 2011 The role of the pod in seed development: strategies for manipulating yield. New Phytologist. 190, 838–853. (10.1111/j.1469-8137.2011.03714.x)

130 Zhang, W. X., Mao, P. S., Li, Y., Wang, M. Y., Xia, F. S., Wang, H. 2017 Assessing of the contributions of pod photosynthesis to carbon acquisition of seed in alfalfa (*Medicago sativa* L.). Scientific Reports. 7, 1–7 (e42026). (10.1038/srep42026)

131 Rosell, J. A., Piper, F. I., Jimenez-Vera, C., Vergilio, P. C. B., Marcati, C. R., Castorena, M., Olson, M. E. 2021 Inner bark as a crucial tissue for non-structural carbohydrate storage across three tropical woody plant communities. Plant Cell and Environment. 44, 156–170. (10.1111/pce.13903)

132 Jones, J. M., Heineman, K. D., Dalling, J. W. 2019 Soil and species effects on bark nutrient storage in a premontane tropical forest. Plant and Soil. 438, 347–360. (10.1007/s11104-019-04026-9)

133 Henkel, T. W., Mayor, J. R. 2019 Implications of a long-term mast seeding cycle for climatic entrainment, seedling establishment and persistent monodominance in a Neotropical, ectomycorrhizal canopy tree. Ecological Research. 34, 472–484. (10.1111/1440-1703.12014)

134 Hart, T. B. 1995 Seed, seedling and sub-canopy survival in monodominant and mixed forests of the Ituri Forest, Africa. Journal of Tropical Ecology. 11, 443–459. (10.1017/s0266467400008919)

135 Camberlin, P., Janicot, S., Poccard, I. 2001 Seasonality and atmospheric dynamics of the teleconnection between African rainfall and tropical sea-surface temperature: Atlantic vs. ENSO. International Journal of Climatology. 21, 973–1005. (10.1002/joc.673)

136 Poccard, I., Janicot, S., Camberlin, P. 2000 Comparison of rainfall structures between NCEP/NCAR reanalyses and observed data over tropical Africa. Climate Dynamics. 16, 897–915. (10.1007/s003820000087)

137 Nascimento, M. T., Proctor, J. 1994 Insect defoliation of a monodominant Amazonian rainforest. Journal of Tropical Ecology. 10, 633–636. (10.1017/s0266467400008312)

138 Maisels, F. 2004 Defoliation of a monodominant rain-forest tree by a noctuid moth in Gabon. Journal of Tropical Ecology. 20, 239–241. (10.1017/s0266467403001044)

139 Van Bael, S. A., Aiello, A., Valderrama, A., Medianero, E., Samaniego, M., Wright, S. J. 2004 General herbivore outbreak following an El Nino-related drought in a lowland Panamanian forest. Journal of Tropical Ecology. 20, 625–633. (10.1017/s0266467404001725)

140 Wong, M., Wright, S. J., Hubbell, S. P., Foster, R. B. 1990 The spatial pattern and reproductive consequences of outbreak defoliation in *Quararibea asterolepis*, a tropical tree. Journal of Ecology. 78, 579–588. (10.2307/2260885)

141 Numata, S., Yamaguchi, K., Shimizu, M., Sakurai, G., Morimoto, A., Alias, N., Azman, N. Z. N., Hosaka, T., Satake, A. 2022 Impacts of climate change on reproductive phenology in tropical rainforests of Southeast Asia. Communications Biology. 5, (10.1038/s42003-022-03245-8)

142 Vleminckx, J., Hogan, J. A., Metz, M. R., Comita, L. S., Queenborough, S. A., Wright, S. J., Valencia, R., Zambrano, M., Garwood, N. C. 2024 Flower production decreases with warmer and more humid atmospheric conditions in a western Amazonian forest. New Phytologist. 241, 1035–1046. (10.1111/nph.19388)

143 Bieleski, R. L. 1973 Phosphate pools, phosphate transport, and phosphate availability. Annual Review of Plant Physiology and Plant Molecular Biology. 24, 225–252. (10.1146/annurev.pp.24.060173.001301)

144 Raboy, V. 2003 Myo-Inositol-1,2,3,4,5,6-hexakisphosphate. Phytochemistry. 64, 1033–1043. (10.1016/s0031-9422(03)00446-1)

145 Kurita, Y., Baba, K., Ohnishi, M., Matsubara, R., Kosuge, K., Anegawa, A., Shichijo, C., Ishizaki, K., Kaneko, Y., Hayashi, M., et al. 2017 Inositol hexakis phosphate is the seasonal phosphorus reservoir in the deciduous woody plant *Populus alba* L. Plant and Cell Physiology. 58, 1477–1485. (10.1093/pcp/pcx106)

146 Mulligan, D. R. 1988 Phosphorus concentrations and chemical fractions in *Eucalyptus* seedlings grown for a prolonged period under nutrient- deficient conditions. New Phytologist. 110, 479–486. (10.1111/j.1469-8137.1988.tb00286.x)

147 Netzer, F., Schmid, C., Herschbach, C., Rennenberg, H. 2017 Phosphorus-nutrition of European beech (*Fagus sylvatica* L.) during annual growth depends on tree age and P-availability in the soil. Environmental and Experimental Botany. 137, 194–207. (10.1016/j.envexpbot.2017.02.009)

148 Rennenberg, H., Herschbach, C. 2013 Phosphorus nutrition of woody plants: many questions - few answers. Plant Biology. 15, 785–788. (10.1111/plb.12078)

149 Marschner, P., ed. 2012 Marschner’s mineral nutrition of higher plants. 3rd edition ed. Amsterdam: Elsevier.

150 Chiou, T. J., Lin, S. I. 2011 Signaling network in sensing phosphate availability in plants. Annual Review of Plant Biology. 62, 185–206. (10.1146/annurev-arplant-042110-103849)

151 Turnbull, C. 2011 Long-distance regulation of flowering time. Journal of Experimental Botany. 62, 4399–4413. (10.1093/jxb/err191)

152 Goldschmidt, E. E., Golomb, A. 1982 The carbohydrate balance of alternate-bearing citrus trees and the significance of reserves for flowering and fruiting. Journal of the American Society for Horticultural Science. 107, 206–208. (10.21273/JASHS.107.2.206)

153 Hansen, P., Ryugo, K., Ramos, D. E., Fitch, L. 1982 Influence of cropping on Ca, K, Mg, and carbohydrate status of ‘French’ prune trees grown on potassium limited soils. Journal of the American Society for Horticultural Science. 107, 511–515. (10.21273/JASHS.107.3.511)

154 Sachs, R. M. 1977 Nutrient diversion: an hypothesis to explain the chemical control of flowering. HortScience. 12, 220–222. (10.21273/HORTSCI.12.3.220)

155 Rosenstock, T. S., Rosa, U. A., Plant, R. E., Brown, P. H. 2010 A reevaluation of alternate bearing in pistachio. Scientia Horticulturae. 124, 149–152. (10.1016/j.scienta.2009.12.007)

156 Zimmerman, M. H., Brown, C. L. 1971 Trees: structure and function. New York: Springer.

157 Chapin, F. S., Schulze, E. D., Mooney, H. A. 1990 The ecology and economics of storage in plants. Annual Review of Ecology and Systematics. 21, 423–447. (10.1146/annurev.ecolsys.21.1.423)

158 Sala, A., Hopping, K., McIntire, E. J. B., Delzon, S., Crone, E. E. 2012a Masting in whitebark pine (*Pinus albicaulis*) depletes stored nutrients. New Phytologist. 196, 189–199. (10.1111/j.1469-8137.2012.04257.x)

159 Dietze, M. C., Sala, A., Carbone, M. S., Czimczik, C. I., Mantooth, J. A., Richardson, A. D., Vargas, R. 2014 Nonstructural Carbon in Woody Plants. Annual Review of Plant Biology. 65, 667–687. (10.1146/annurev-arplant-050213-040054)

160 Hartmann, H., Trumbore, S. 2016 Understanding the roles of nonstructural carbohydrates in forest trees - from what we can measure to what we want to know. New Phytologist. 211, 386–403. (10.1111/nph.13955)

161 Brown, P. H., Weinbaum, S. A., Picchioni, G. A. 1995 Alternate bearing influences annual nutrient consumption and the total nutrient content of mature pistachio trees. Trees-Structure and Function. 9, 158–164. (10.1007/bf02418205)

162 Picchioni, G. A., Brown, P. H., Weinbaum, S. A., Muraoka, T. T. 1997 Macronutrient allocation to leaves and fruit of mature, alternate-bearing pistachio trees: Magnitude and seasonal patterns at the whole-canopy level. Journal of the American Society for Horticultural Science. 122, 267–274. (10.21273/jashs.122.2.267)

163 Stevenson, M. T., Shackel, K. A. 1998 Alternate bearing in pistachio as a masting phenomenon: Construction cost of reproduction versus vegetative growth and storage. Journal of the American Society for Horticultural Science. 123, 1069–1075. (10.21273/jashs.123.6.1069)

164 Smaill, S. J., Clinton, P. W., Allen, R. B., Davis, M. R. 2011 Climate cues and resources interact to determine seed production by a masting species. Journal of Ecology. 99, 870–877. (10.1111/j.1365-2745.2011.01803.x)

165 Leeper, A. C., Lawrence, B. A., LaMontagne, J. M. 2020 Plant-available soil nutrients have a limited influence on cone production patterns of individual white spruce trees. Oecologia. 194, 101–111. (10.1007/s00442-020-04759-w)

166 Sala, A., Woodruff, D. R., Meinzer, F. C. 2012b Carbon dynamics in trees: feast or famine? Tree Physiology. 32, 764–775. (10.1093/treephys/tpr143)

167 Norghauer, J. M., Newbery, D. M. 2011 Seed fate and seedling dynamics after masting in two African rain forest trees. Ecological Monographs. 81, 443–468. (10.1890/10-2268.1)

168 Janzen, D. H. 1974 Tropical blackwater rivers, animals and mast fruiting by the Dipterocarpaceae. Biotropica. 6, 69–103. (10.2307/2989823)

169 Burgess, P. F. 1972 Studies on the regeneration of the hill forests of the Malay Peninsular. The phenology of dipterocarps. Malaysian Forester. 35, 103–123.

170 Sakai, S., Kitajima, K. 2019 Tropical phenology: Recent advances and perspectives. Ecological Research. 34, 50–54. (10.1111/1440-1703.1131)

171 Bormann, F. H. 1966 Structure, function, and ecological significance of root grafts in *Pinus strobus* L. Ecological Monographs. 36, 1–26. (10.2307/1948486)

172 Larue, C. D. 1952 Root-grafting in tropical trees. Science. 115, 296–296. (10.1126/science.115.2985.296)

173 Basnet, K., Scatena, F. N., Likens, G. E., Lugo, A. E. 1993 Ecological consequences of root grafting in tabonuco (*Dacryodes excelsa*) trees in the Luquillo Experimental-Forest, Puerto Rico. Biotropica. 25, 28–35. (10.2307/2388976)

174 Saunier, R. E., Wagle, R. F. 1965 Root grafting in *Quercus turbinella* Greene. Ecology. 46, 749–750. (10.2307/1935020)

175 Loehle, C., Jones, R. H. 1990 Adaptive significance of root grafting in trees. Functional Ecology. 4, 268–271.

176 Noble, A. E., Rosenstock, T. S., Brown, P. H., Machta, J., Hastings, A. 2018 Spatial patterns of tree yield explained by endogenous forces through a correspondence between the Ising model and ecology. Proceedings of the National Academy of Sciences of the United States of America. 115, 1825–1830. (10.1073/pnas.1618887115)

177 Fernandez-Martinez, M., Bogdziewicz, M., Espelta, J. M., Penuelas, J. 2017a Nature beyond linearity: meteorological variability and Jensen’s Inequality can explain mast seeding behavior. Frontiers in Ecology and Evolution. 5, 1–8. (10.3389/fevo.2017.00134)

178 Lyles, D., Rosenstock, T. S., Hastings, A., Brown, P. H. 2009 The role of large environmental noise in masting: General model and example from pistachio trees. Journal of Theoretical Biology. 259, 701–713. (10.1016/j.jtbi.2009.04.015)

179 Rees, M., Kelly, D., Bjornstad, O. N. 2002 Snow tussocks, chaos, and the evolution of mast seeding. American Naturalist. 160, 44–59. (10.1086/340603)

180 Cannell, M. G. R., Dewar, R. C. 1994 Carbon allocation in trees: a review of concepts for modeling. In Advances in Ecological Research*, Vol* 25. (ed.^eds. pp. 59–104

181 Lacointe, A. 2000 Carbon allocation among tree organs: A review of basic processes and representation in functional-structural tree models. Annals of Forest Science. 57, 521–533. (10.1051/forest:2000139)

182 Makela, A., Valentine, H. T., Helmisaari, H. S. 2008 Optimal co-allocation of carbon and nitrogen in a forest stand at steady state. New Phytologist. 180, 114–123. (10.1111/j.1469-8137.2008.02558.x)

183 Franklin, O., Johansson, J., Dewar, R. C., Dieckmann, U., McMurtrie, R. E., Brannstrom, A., Dybzinski, R. 2012 Modeling carbon allocation in trees: a search for principles. Tree Physiology. 32, 648–666. (10.1093/treephys/tpr138)

184 Farrar, J. F. 1993 Sink strength: what is it and how do we measure it? A summary. Plant Cell and Environment. 16, 1045–1046. (10.1111/j.1365-3040.1996.tb02061.x)

185 Kozlowski, T. T. 1971 Growth and development of trees. Vol. II. Cambial growth, root growth and reproductive growth. New York: Academic Press.

186 Kozlowski, T. T. 1992 Carbohydrate sources and sinks in woody plants. Botanical Review. 58, 107–222. (10.1007/bf02858600)

187 Minchin, P. E. H., Thorpe, M. R. 1996 What determines carbon partitioning between competing sinks? Journal of Experimental Botany. 47, 1293–1296. (10.1093/jxb/47.Special_Issue.1293)

188 Wardlaw, I. F. 1990 The control of carbon partitioning in plants. New Phytologist. 116, 341–381. (10.1111/j.1469-8137.1990.tb00524.x)

189 Thornley, J. H. M., Parsons, A. J. 2014 Allocation of new growth between shoot, root and mycorrhiza in relation to carbon, nitrogen and phosphate supply: Teleonomy with maximum growth rate. Journal of Theoretical Biology. 342, 1–14. (10.1016/j.jtbi.2013.10.003)

190 Le Roux, X., Lacointe, A., Escobar-Gutierrez, A., Le Dizes, S. 2001 Carbon-based models of individual tree growth: A critical appraisal. Annals of Forest Science. 58, 469–506.

