## Supplementary material for "Mast fruiting in a large tropical African legume tree species and the mineral nutrient limitation hypothesis": Suppementary Electronic Materials

### ELECTRONIC SUPPLEMENTARY MATERIAL

#### Contents

Appendix 1. Radiometry at Bulu, Ndian: calibration and interpretation.

Appendix 2. Evidence table for the levels of mast fruiting per year.

Appendix 3. Time series analysis of dry season climate variables.

Appendix 4. Regression model fits of masting on rainfall and radiation.

Appendix 5. Rainfall and 30-day rainfall totals defining dry seasons.

Appendix 6. Description and feeding of the yellow-green caterpillar.

Appendix 7. Analysis of Ndian PAMOL oil-palm plantation yields.

Appendix 8. Stem increments from tree cores and mast fruiting.

Appendix 9. Minimum temperature and mast fruiting.

**Appendix 1: Radiometry measurements at Bulu, Ndian.**

From the onset in 1984 radiation was recorded at the PAMOL station, at Bulu, Ndian (80 m a.s.l), using a Gunn-Bellani radiometer (Baird and Tatlock, London), in units of ml water evaporated per day (V). Due to a breakage of this (GB1) instrument, data were then missing for 3 April 2009 through 29 May 2011 (which included the dry seasons of 2010 and 2011), until a replacement (GB2) of the same design could be installed[1]. A pyranometer (Skye Instruments, Powys, Wales; model SK1110) recorded at the same station in the periods 1 September 2006 to 9 May 2007 (5 runs) and 9 June 2012 to 21 June 2013 (3 runs), overlapping the services of both GB radiometers. Averaging 10-min interval readings per day, this led to two best-reliable calibration equations relating radiation (R; Watts/$m^2$) to V (ml/day):

GB1: $R = 24.1 + 13.2 \cdot V$ [$n = 220$, $r^2 = 0.898$]

GB2: $R = 21.1 + 14.7 \cdot V$ [$n = 246$, $r^2 = 0.924$]

The Skye recordings were further checked against another pyranometer (Kipp and Zonen, Delft, Neths; Model CM3), 16 December 2009 through 7 January 2010 (again averaging 10-min interval readings), leading to a regression fit:

$R_{SKYE} = -0.52 + 0.935 \cdot R_{KIPP}$ [$n = 23$, $r^2 = 0.961$].

The difference in the Skye-Kipp regression slope from 1.0 was likely due to the slightly differing spectral profiles of the two pyranometers. The two calibration lines for GB1 and GB2 make the radiation data for the two phases fully comparable.

*The climate data for Bulu, 1984 to 2017, are placed in a Zenodo archive (Etta et al. 2022).*

To estimate the missing mean daily radiation for dry seasons 2010 and 2011, daily radiation values (W/$m^2$) across all the other dry seasons were matched to maximum temperature (T, deg. C; 07:00 h) and the best fitting relationship was found as:

$R = 325.3 + 0.813 \cdot Y - 5.85 \cdot T$ [$n = 32$, $r^2 = 0.507$],

Where Y is year number set at 1 for 1984 through to 32 for 2017. Fitting radiation to T alone achieved a weak predictive value ($r^2 = 0.073$). Two other alternative were (a) to take a simple linear interpolation of the radiation values for 2009 and 2012, and (b) use the Holt-Winters filtering method (`HoltWinters()` in R package 'forecast') on the time series of monthly mean radiation between 1984 and 2009, and predict the weighted mean values for closest corresponding months of the dry seasons 2010 and 2011, V = 9.780 and 10.047 ml/day respectively[2].

| W $m^{-2}$ | Regression model | Linear interpolation | Time-series forecast |
|---|---|---|---|
| 2010 | 156.13 | 146.00 | 153.20 |
| 2011 | 163.43 | 154.67 | 156.72 |

[1] Kindly donated by Dr Gerald Stanhill of the Agricultural Research Organization, Volcani Centre, Ben-Dragan, Israel.

[2] For 2010: mean {Dec. 2009, Jan. 2010); for 2011 mean {[Jan. 2011] and mean [Feb. and Mar. 2011]}. Refer to Table 1 main text.

Averages of these three estimates were used to replace the missing radiation values for dry seasons 2010 and 2011 (151.78 and 158.27 W $m^{-2}$). Interpolation of the missing months (`na.interp()` in R package 'forecast') gave slightly higher values of 160.48 and 173.33 W $m^{-2}$ for 2010 and 2011, but were strongly influenced by an unusually high value recorded for May 2011 at the restart of the non-missing series.

The radiation vales presented in the paper here are lower than those in Newbery et al. (2006) which applied the calibration equation of Pereira (1959). That author derived the relationship:

$R' = 130 + 36.5 \cdot V$, where R' was presented in unit g·cal $cm^{-2}day^{-1}$, V as above.

The conversion from g·cal $cm^{-2}day^{-1}$ to W $m^{-2}$ is $R = 0.4846 \cdot R'$. Which led to:

$R = 63.0 + 17.69 \cdot V$, with R in W $m^{-2}$. [as on p. 562, ibid.].

This equation applied to the radiometer data up to 2004 (i.e. readings from GB1), but the new calibrations for Bulu show that radiation had been over-estimated. Nevertheless, the new estimates are linearly related to the Pereira ($R_{Per}$) ones:

$R_{GB1} = -19.4 + 0.69 \cdot R_{Per}$ and $R_{GB2} = -34.2 + 0.84 \cdot R_{Per,}$

And the relative comparisons across the dry seasons remain valid.

Recalibrated radiation for 1989-2004 (W $m^{-2}$) replacing col. 6 in Table 1a of Newbery *et al.* (2006): 150.6, 140.2, 142.7, 140.6, 136.8, 141.1, 142.1, 142.4, 145.6, 162.5, 165.4, 164.0, 173.8, 159.0, 174.3, 162.3.

Coda 1: In contrast to Pereira (1959) at Mugaga, Kenya (2700 m a.s.l.), Davies (1965) derived different calibration lines for coastal Benin and Ibadan, Nigeria, respectively, as:

$R = 9.8 + 9.50 \cdot V$ and $R = 12.5 + 8.9 \cdot V$ [R converted to W $m^{-2}$].

Differences between the sites' calibration values are likely due to elevation and how the radiometer was set up (see Monteith and Szeicz 1960, and McCulloch and Wangati 1967). The value at Ndian fit well for what is expected generally for such lowland western Central African location, 5° N (Linacre 1992, Campbell and Norman 1998).

Coda 2: Solar radiation interpolated for Bulu (8.52 E, 4.75N) and extracted from the WorldClim 2 data base, http://worldclim.org/ (Fick and Hijmans 2017) gave the following values, averaged for the period 1970-2000:

| W $m^{-2}$ | Dec | Jan | Feb | Mar |
|---|---|---|---|---|
| WorldClim 2 | 178.3 | 180.1 | 190.7 | 189.2 |
| Bulu radiometer | 157.3 | 153.4 | 164.1 | 166.1 |

These are 15% higher than those recorded by instruments at the site (1984-2008), although closer to the interpolated estimates, and the most likely reasons are (a) the Sahel atmospheric dust load in the dry season in SW Cameroon caused by ITCZ movement, (b) difference in dates compared when radiation in trending upwards over time at the Ndian site.

Coda 3. A separate attempt was made to estimate the missing months of radiation by accessing the cloud cover interpolated for the 0.5 x 0.5 deg. grid cell centred on 4. 75 N, 8.75 E from the CRU TS4.04 archive (Harris et al. 2020). However, for the months in 1984 to 2016 recorded, radiation (W $m^{-2}$) was poorly correlated with cloud cover (%) (radi = 264 – 1.391·cld%, $R^2$ = 37%). The nearest stations were Tiko, Calabar, Mamfe and Douala, all outside the cell and incomplete or lacking sunshine hours recording). Furthermore, the estimated cloud cover in the dry season was far too high for Korup (ca. 70%) and total annual rainfall ~50% of actuality.  Radiation at Bulu was also poorly related to OLR (W $m^{-2}$) (radi = 84.8 + 0.294·OLR, $R^2$ = 7%). In the dry season months, DJF, these two variables we barely correlated ($r$ = 0.084, $P$ = 0.425). Given the sparse network of stations in western Central Africa, the special topographic effects inland (north) of Mt. Cameroon and such world data base estimates are likely to be highly biased, unreliable and hence erroneous.

References:


Campbell, G. S., and J. M. Norman. 1998. An introduction to environmental biophysics. Springer-Verlag, New York.

Davies, J. A. 1965. The use of a Gunn-Bellani distillator to determine net radiative flux in West Africa. Journal of Applied Meteorology **4**:547-549.

Etta, C. E., G. B. Chuyong, and D. M.  Newbery. 2022. Climate records for Bulu, Ndian Division, SW Cameroon. [Data set]. Zenodo. https://doi.org/10.5281/zenodo.7220319.

Harris, I., Osborn, T.J., Jones, P. *et al.* Version 4 of the CRU TS monthly high-resolution gridded multivariate climate dataset. *Sci Data* **7,** 109 (2020).

Fick, S. E., and R. J. Hijmans. 2017. WorldClim 2: new 1-km spatial resolution climate surfaces for global land areas. International Journal of Climatology **37**:4302-4315.

Linacre, E. 1992. Climate data and resources. Routledge, London, UK.

McCulloch, J. S. G., and F. J. Wangati. 1967. Notes on the use of the Gunn Bellani radiometer. Applied Meteorology **4**:63-70.

Monteith, J. L., and G. Szeicz. 1960. The performance of a Gunn-Bellani radiation integrator. Quarterly Journal of the Royal Meteorological Society **86**:91-94.

Pereira, H. C. 1959. Practical field instruments for estimation of radiation and of evaporation. Quarterly Journal of the Royal Meteorological Society **85**:253-261.

**Appendix 2: Table S1.** Evidence table for the mast fruiting events of *Microberlinia bisulcata* at Korup (2005-2017) [a].

| year | Intensity of masting | Source of evidence |
|---|---|---|
| 2005 | None/v. low | Personal observations [DMN] |
| 2006 | None/v. low | Setting up of the new phenology plot [DMN] |
| 2007 | High | Mb dispersal study (Norghauer & Newbery 2015); phenology recording: 63/65 trees flowering, 64/65 pod fall |
| 2008 | None/v. low | Field observations of JMN and DMN |
| 2009 | Low | From unpublished inter-mast pod predation study (JMN): 12/61 trees mature pods, of which 3 'heavy' (> 5000 per tree estimated: CF. Norghauer & Newbery 2015, Fig. 3a). |
| 2010 | High | Mb dispersal study (Norghauer & Newbery 2015); phenology recording: 52/61 trees mature pods, 51 'heavy'. |
| 2011 | None | Phenology recording |
| 2012 | Low | Phenology recording: good flowering but 15/61 trees with some pods, only 2 'heavy'. |
| 2013 | Moderate | Phenology recording: moderate flowering; 34/61 trees with some pods, none 'heavy'; at IsRd grove 21/33 of which 14 'heavy' |
| 2014 | Moderate | Pods on floor for 20/58 trees, only 2 'heavy'. Many pods around Mb trees in ECM exclusion expt (Newbery and Neba (2019) |
| 2015 | None | No new pods on floor; pers. observations and SN. |
| 2016 | Moderate | 40% trees with some pods, 5% heavy (GN estimate). |
| 2017 | Moderate | No direct records of mature pods, but many with green pods in May-June 2017 [DMN] |

[a] for 1988-2004 refer to Newbery et al. (2006a), cited in main paper.

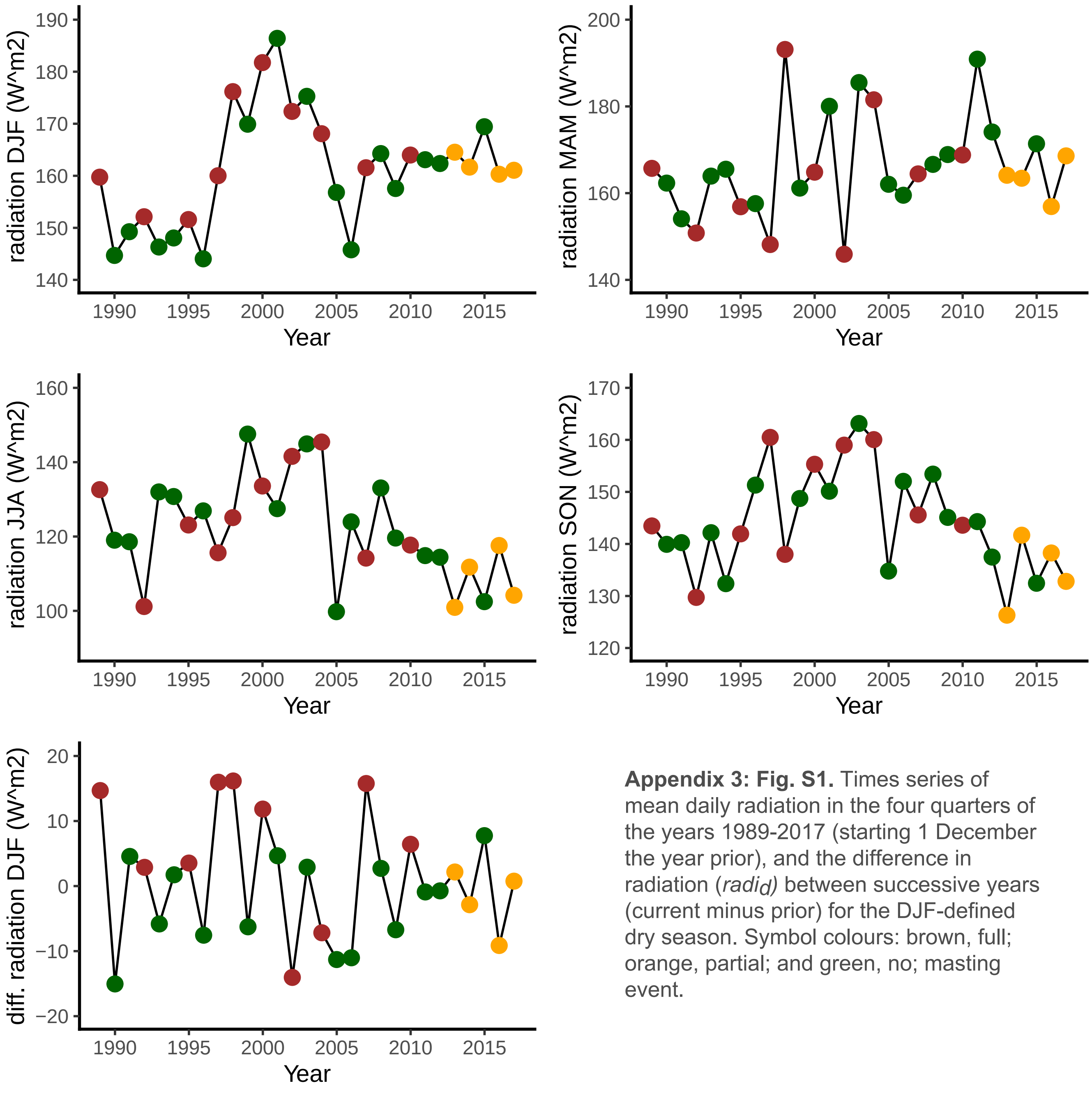


**Appendix 3: Fig. S1.** Times series of mean daily radiation in the four quarters of the years 1989-2017 (starting 1 December the year prior), and the difference in radiation (*radi*$_d$) between successive years (current minus prior) for the DJF-defined dry season. Symbol colours: brown, full; orange, partial; and green, no; masting event.

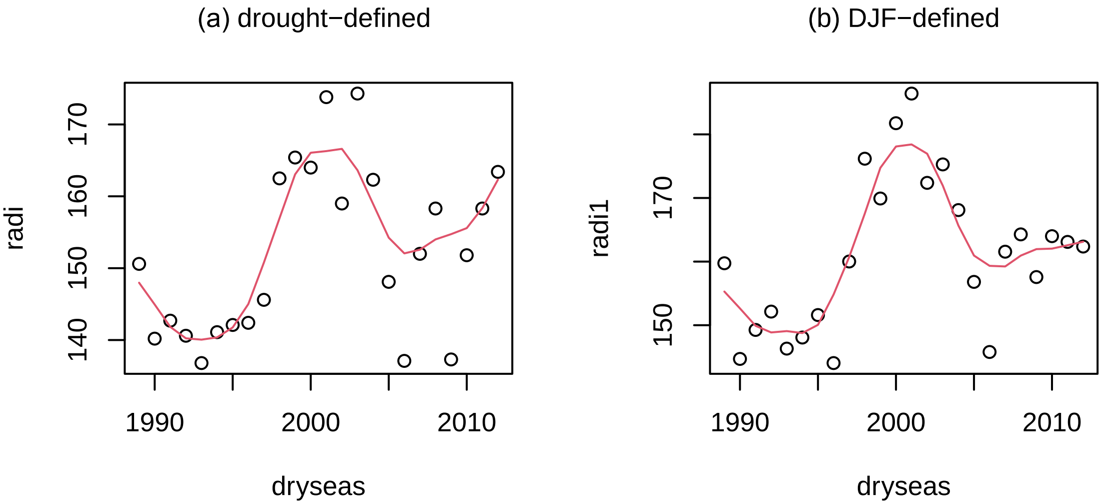


**Appendix 3: Fig. S2.** Time course of mean daily radiation (W $m^{-2}$) in dry seasons 1989-2012 defined by (a) droughted days, and (b) days of the months December to February. Red lines are LOWESS smoothed fits (see main text).

**Appendix 3: Fig. S3.** Autocorrelation (ACF) functions of the times series of the dry season climatic variables associated with the masting series and graphed in Fig. 2 of the main text: mean daily rainfall (rain) and radiation (radi); start date (start) and duration (dur) of the season, radiation intensity (radi.dur); and the difference in radiation (radi.diff) between successive years.

**Appendix 3: Fig. S4.** Autocorrelation (ACF) functions of the times series of mean daily radiation in the four quarters of the years 1989-2017 (starting 1 December the year prior), and the difference in radiation (radi.diff = $radi_d$) between successive years (current minus prior) for the 'DJF-defined' dry season. Series are graphed in Appendix 3: Fig. S1.

Appendix 3: Fig. S3

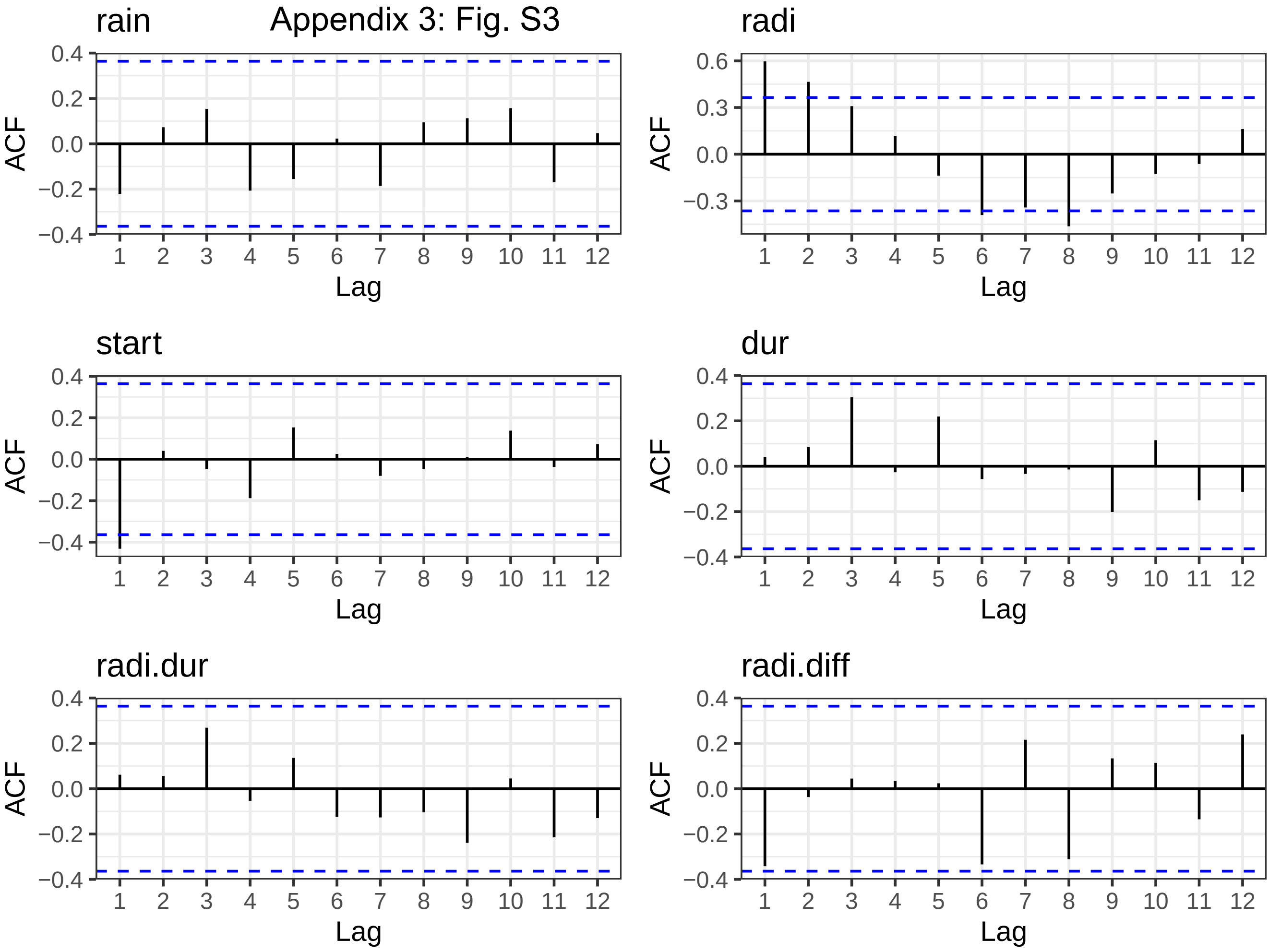

Appendix 3: Fig. S4

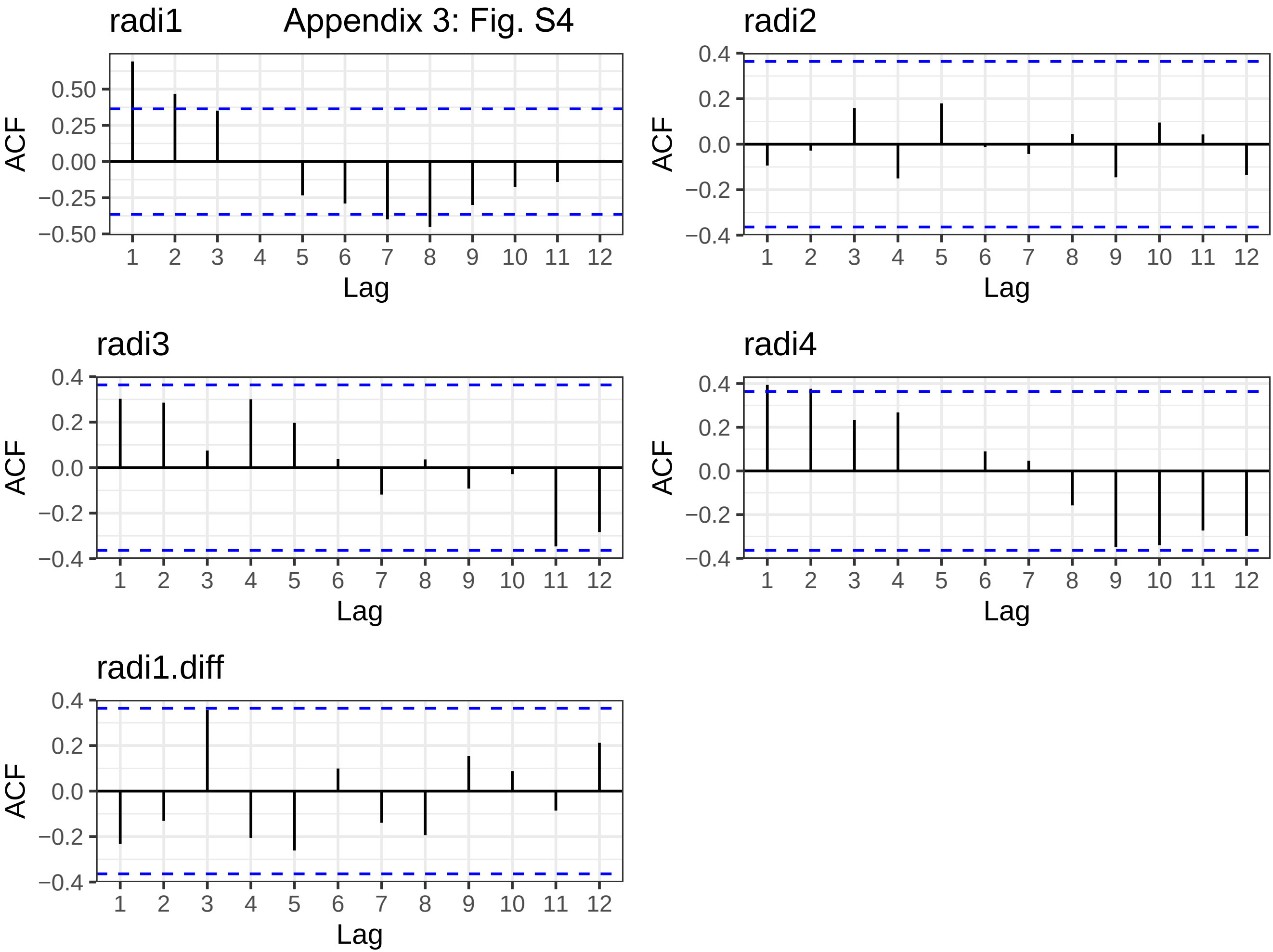

**Appendix 4: Table S1.** Estimates of coefficients from regression fits of masting on mean daily rainfall**,** in the current-year (*rain*) and the year-before (*rain.1*) dry season, ln-transformed, for 1989-2012 by generalized linear models (glm, glarma) and for 1989-2017 by proportional odds (ordinal) logistic regression (polr). Under 'S', coefficients are from two single-term models; under 'D', from one (double) two-term model. The binomial time series model, glarma-1 had p = 1 lag, and glarma-2 p= 2 lags.

| period | model | | β ± SE | *t*-value | P(t) | $e^{\beta}$ |
|---|---|---|---|---|---|---|
| 1989-2012 | glm-S | | | | | |
| | | ln(rain) | −6.38 ± 2.53 | −2.56 | 0.012 | $1.69 \cdot 10^{-3}$ |
| | | ln($rain_{-1}$) | 4.99 ± 2.00 | 2.50 | 0.013 | $1.47 \cdot 10^{2}$ |
| | glm-D | | | | | |
| | | ln(rain) | −9.26 ± 5.15 | −1.80 | 0.072 | $9.55 \cdot 10^{-5}$ |
| | | ln($rain_{-1}$) | 8.49 ± 4.54 | 1.87 | 0.061 | $4.86 \cdot 10^{3}$ |
| | glarma-1 | | | | | |
| | | ln(rain) | −7.38 ± 2.69 | −2.75 | 0.006 | $6.22 \cdot 10^{-4}$ |
| | | ln($rain_{-1}$) | 4.66 ± 2.20 | 2.12 | 0.034 | $1.06 \cdot 10^{2}$ |
| | glarma-2 | | | | | |
| | | ln(rain) | −6.90 ± 2.45 | −2.82 | 0.005 | $1.01 \cdot 10^{-3}$ |
| | | ln($rain_{-1}$) | 4.98 ± 2.08 | 2.40 | 0.016 | $1.46 \cdot 10^{2}$ |
| 1989-2017 | polr-S | | | | | |
| | | ln(rain) | −4.00 ± 1.37 | −2.93 | 0.007 | $1.83 \cdot 10^{-2}$ |
| | | ln($rain_{-1}$) | 4.17 ± 1.47 | 2.83 | 0.009 | $6.47 \cdot 10^{1}$ |
| | polr-D | | | | | |
| | | ln(rain) | −3.20 ± 1.30 | −2.47 | 0.021 | $4.08 \cdot 10^{-2}$ |
| | | ln($rain_{-1}$) | 3.72 ± 1.55 | 2.40 | 0.024 | $4.13 \cdot 10^{1}$ |

**Appendix 4: Table S2.** Comparison of mean (± SE) daily rainfall (*rain*, mm/day) in the dry seasons of current year of masting ($t_0$) and each of the two prior years ($t_{-1}$, $t_{-2}$)—using paired *t*-tests on ln-transformed values, for the ten 'M' masting events of the time series 1989-2012. Further comparisons entailed the means of the two prior years with either equal or unequal weighting.

| | ln(*rain*) | | | |
|---|---|---|---|---|
| Test | $t_0$ | $t_{-1}$ / $t_{-2}$ | *t*-values | P(*t*) |
| $t_0$ vs. $t_{-1}$ | 0.318 ± 0.100 | 0.913 ± 0.115 | −4.10 | 0.003 |
| $t_{-2}$ | | 0.525 ± 0.104 | −1.58 | 0.149 |
| $(t_{-1} + t_{-2})/2$ | | 0.763 ± 0.076 | −3.97 | 0.003 |
| $(t_{-1} \cdot 2 + t_{-2})/3$ | | 0.820 ± 0.186 | −4.18 | 0.002 |

**Appendix 4: Table 3.** Estimates of coefficients from regression fits of masting on mean daily radiation: (a) in the current-year (*radi*) and the year-before ($radi_{-1}$) first quarter of the 12 month phenological year (DJF) dry season, and the difference in radiation between successive years ('current-minus-before', $radi_d$) or corresponding percent change on current year ($radi_{d\%}$), for 1989-2012 by generalized linear models (glm and glarma) and for 1989-2017 by proportional odds (ordinal) logistic regression (polr); (b) similarly for *radi* and $radi_{-1}$, but as differences from fitted LOWESS regressions over time. The binomial time series model, glarma-2 used p = 2 lags.

(a)

| period | model | β ± SE | *t*-value | P(t) | $e^{\beta}$ |
|---|---|---|---|---|---|
| 1989-2012 | glm | | | | |
| | radi | 0.050 ± 0.038 | 1.32 | 0.188 | 1.05 |
| | $radi_{-1}$ | –0.024 ± 0.035 | –0.69 | 0.493 | 0.98 |
| | $radi_d$ | 0.143 ± 0.063 | 2.27 | 0.023 | 1.15 |
| | $radi_{d\%}$ | 0.246 ± 0.106 | 2.33 | 0.020 | 1.28 |
| | glarma-2 | | | | |
| | radi | 0.053 ± 0.039 | 1.34 | 0.181 | 1.05 |
| | $radi_{-1}$ | –0.027 ± 0.037 | –0.74 | 0.459 | 0.97 |
| | $radi_d$ | 0.230 ± 0.085 | 2.71 | 0.007 | 1.26 |
| | $radi_{d\%}$ | 0.381 ± 0.146 | 2.61 | 0.009 | 1.46 |
| 1989-2017 | polr | | | | |
| | radi | 0.046 ± 0.037 | 1.26 | 0.220 | 1.05 |
| | $radi_{-1}$ | –0.024 ± 0.035 | –0.71 | 0.486 | 0.98 |
| | $radi_d$ | 0.124 ± 0.051 | 2.41 | 0.024 | 1.13 |
| | $radi_{d\%}$ | 0.212 ± 0.084 | 2.52 | 0.019 | 1.24 |

(b)

| period | model | β ± SE | *t*-value | P(t) | $e^{\beta}$ |
|---|---|---|---|---|---|
| 1989-2012 | glm | | | | |
| | radi | 0.240 ± 0.123 | 1.95 | 0.051 | 1.27 |
| | $radi_{-1}$ | –0.159 ± 0.095 | –1.68 | 0.093 | 0.85 |
| | glarma-2 | | | | |
| | radi | 0.270 ± 0.115 | 2.35 | 0.019 | 1.31 |
| | $radi_{-1}$ | –0.167 ± 0.095 | –1.75 | 0.080 | 0.85 |
| 1989-2017 | polr | | | | |
| | radi | 0.196 ± 0.098 | 2.00 | 0.057 | 1.22 |
| | $radi_{-1}$ | –0.145 ± 0.082 | –1.77 | 0.089 | 0.87 |

**Appendix 4: Table 4**. Estimates of coefficients from regression fits of masting on minimum daily temperature (minT) as the mean of minT (*mn_*) and the minimum of minT (*min_*), in the current-year's drought-defined (*ds*) and quarter-year-defined (months *DJF*) dry seasons, and in the most recent quarter-year-defined wet season prior to the dry season (months $SON_{-1}$), for 1989-2012 by the generalized linear model (glm) and for 1989-2017 by the proportional odds (ordinal) logistic regression (polr).

| period | model | β ± SE | t-value | P(t) | $e^{\beta}$ |
|---|---|---|---|---|---|
| 1989-2012 | glm | | | | |
| | mn_ds | −1.046 ± 0.696 | −1.50 | 0.133 | 0.35 |
| | min_ds | −0.355 ± 0.268 | −1.33 | 0.185 | 0.70 |
| | mn_DJF | −1.182 ± 0.704 | −1.68 | 0.093 | 0.31 |
| | min_DJF | −0.237 ± 0.295 | −0.80 | 0.422 | 0.77 |
| | mn_$SON_{-1}$ | −0.745 ± 0.722 | −1.03 | 0.302 | 0.47 |
| | min_ $SON_{-1}$ | −0.404 ± 0.469 | −0.86 | 0.389 | 0.67 |
| 1989-2017 | polr | | | | |
| | mn_ds | −0.457 ± 0.422 | −1.08 | 0.289 | 0.63 |
| | min_ds | −0.273 ± 0.226 | −1.21 | 0.237 | 0.76 |
| | mn_DJF | −0.576 ± 0.424 | −1.36 | 0.187 | 0.56 |
| | min_DJF | −0.161 ± 0.238 | −0.68 | 0.505 | 0.85 |
| | mn_ $SON_{-1}$ | −0.266 ± 0.413 | −0.65 | 0.524 | 0.77 |
| | min_ $SON_{-1}$ | −0.212 ± 0.348 | −0.61 | 0.548 | 0.81 |

$e^{\beta}$ is the multiplicative factor by which masting changes for each unit increase in minT.

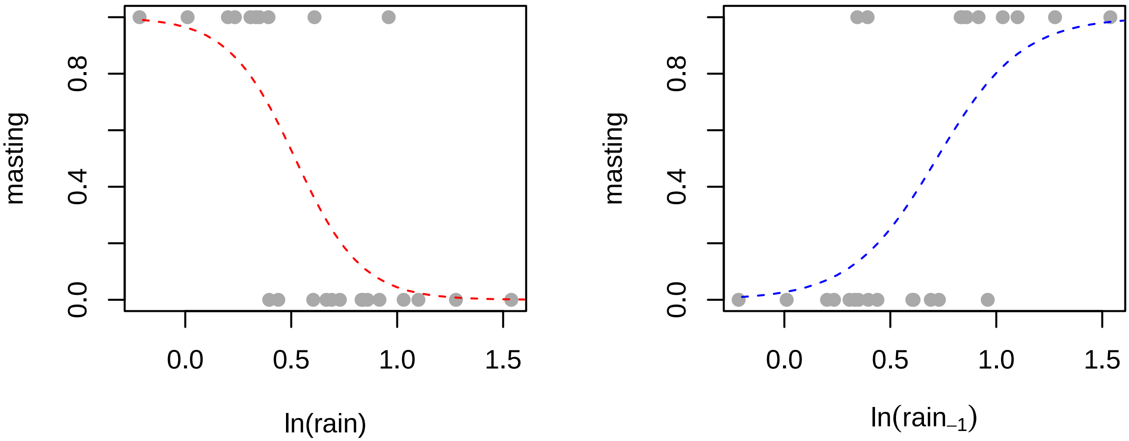


**Appendix 4: Fig. S1.** Dependence of masting in years 1989-2012 on the logarithm of mean daily rainfall, in the current year's drought-defined dry season [ln(rain)] or the same in the dry season in the year prior [ln($rain_{-1}$)].

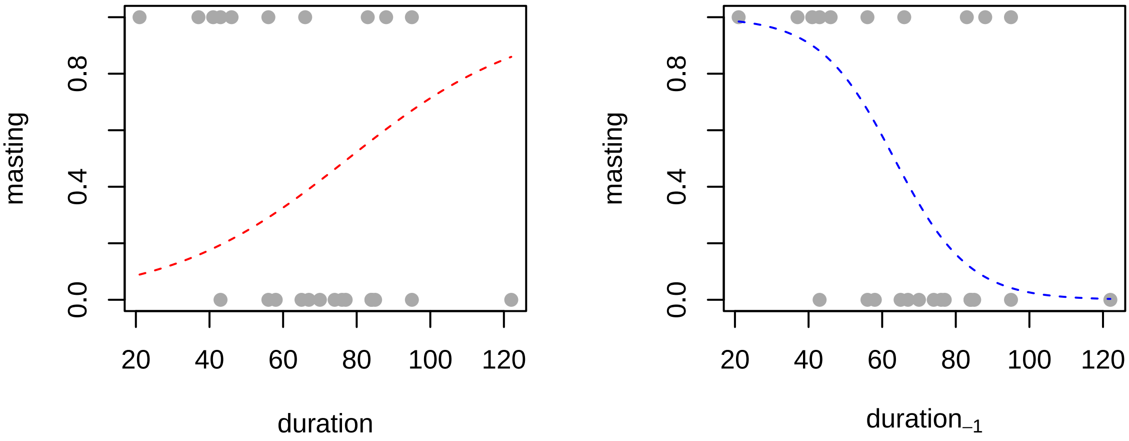


**Appendix 4: Fig. S2.** Dependence of masting in years 1989-2012 on the length of the current year's drought-defined dry season (duration) or the same in the dry season in the year prior ($duration_{-1}$).

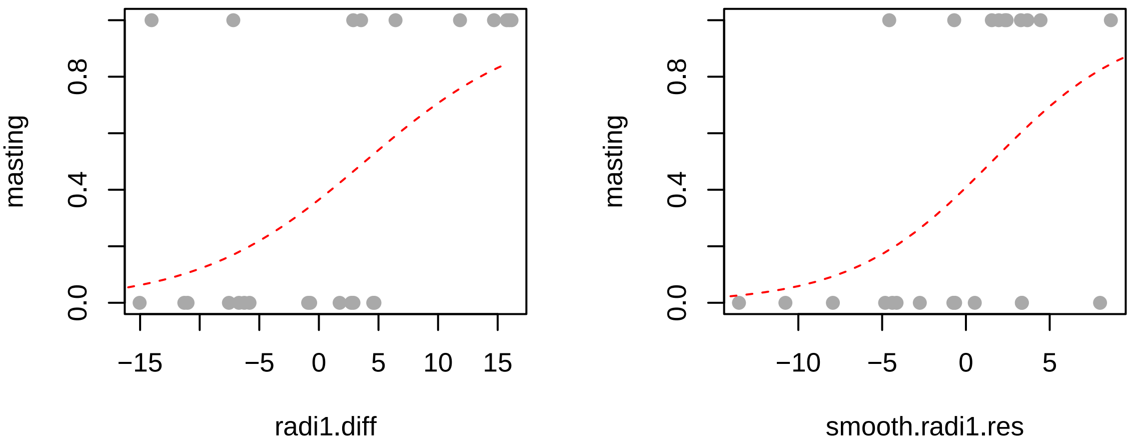


**Appendix 4: Fig. S3.** Dependence of masting in years 1989-2012 on the difference in radiation (current - prior) between DJF-defined dry seasons, and the same on the difference in radiation from a smoothed regression fit to radiation over time.

**Appendix 4: Fig. S4.** Relationships between differences ('_d') in radiation (W $m^{-2}$): (a) between successive years (current – previous), and (b) residuals from smoothed LOWESS curve ('_sr') for the current year, for radiation in the DJF-defined (*radi1*) versus drought-defined dry season (*radi*), for 1989-2012. The dashed lines are 1:1 references.

(a)

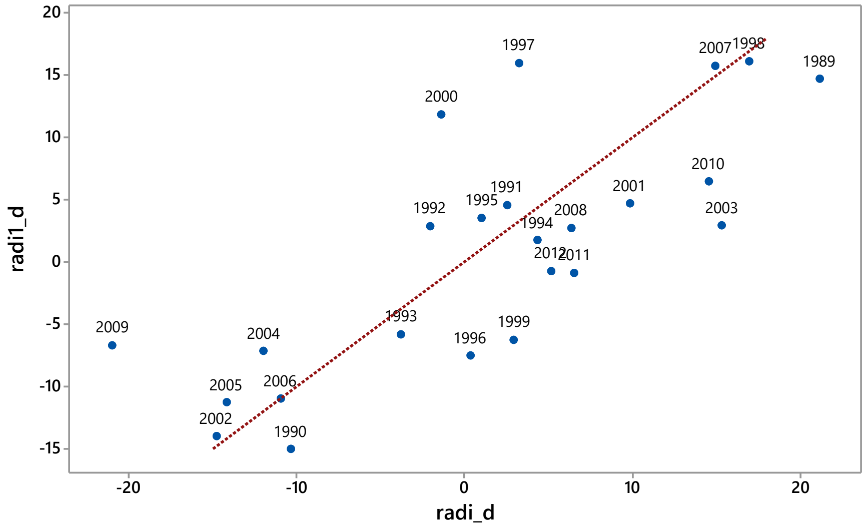


(b)

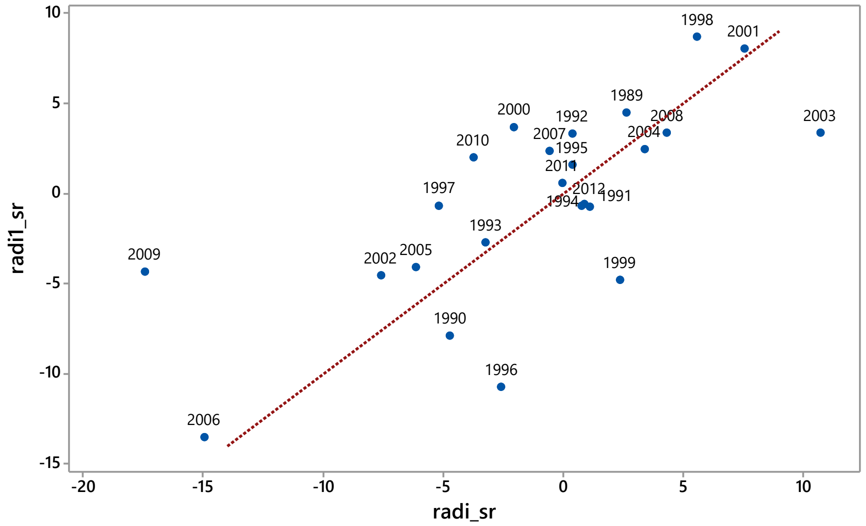

**Appendix 5: Fig. S1.** The course of daily rainfall events between 1 December one year to 30 April the following year (green closed circles; left axis) and the 30-day running rainfall total, Rft (blue dashed line; right axis) for the same period, these dates encompassing the dry season and early following wet season: (a) 2002-2003, (b) 2003-2004, and (c) 2004-2005). The red line defines the dry season, when Rft remains (almost entirely) below 100 mm. In Newbery *et al.* (1996: Table 1) the case was made to consider the 2002-2003 dry season as being unusual in that it was long and made up of three parts.

**Appendix 5: Fig. S2.** As Fig. S1, for (a) 2009-2010, (b) 2010-2011, and (c) 2011-2012, where in (c) the ‘C’s indicate the caterpillar attacks that year.

Appendix 5: Fig. S1

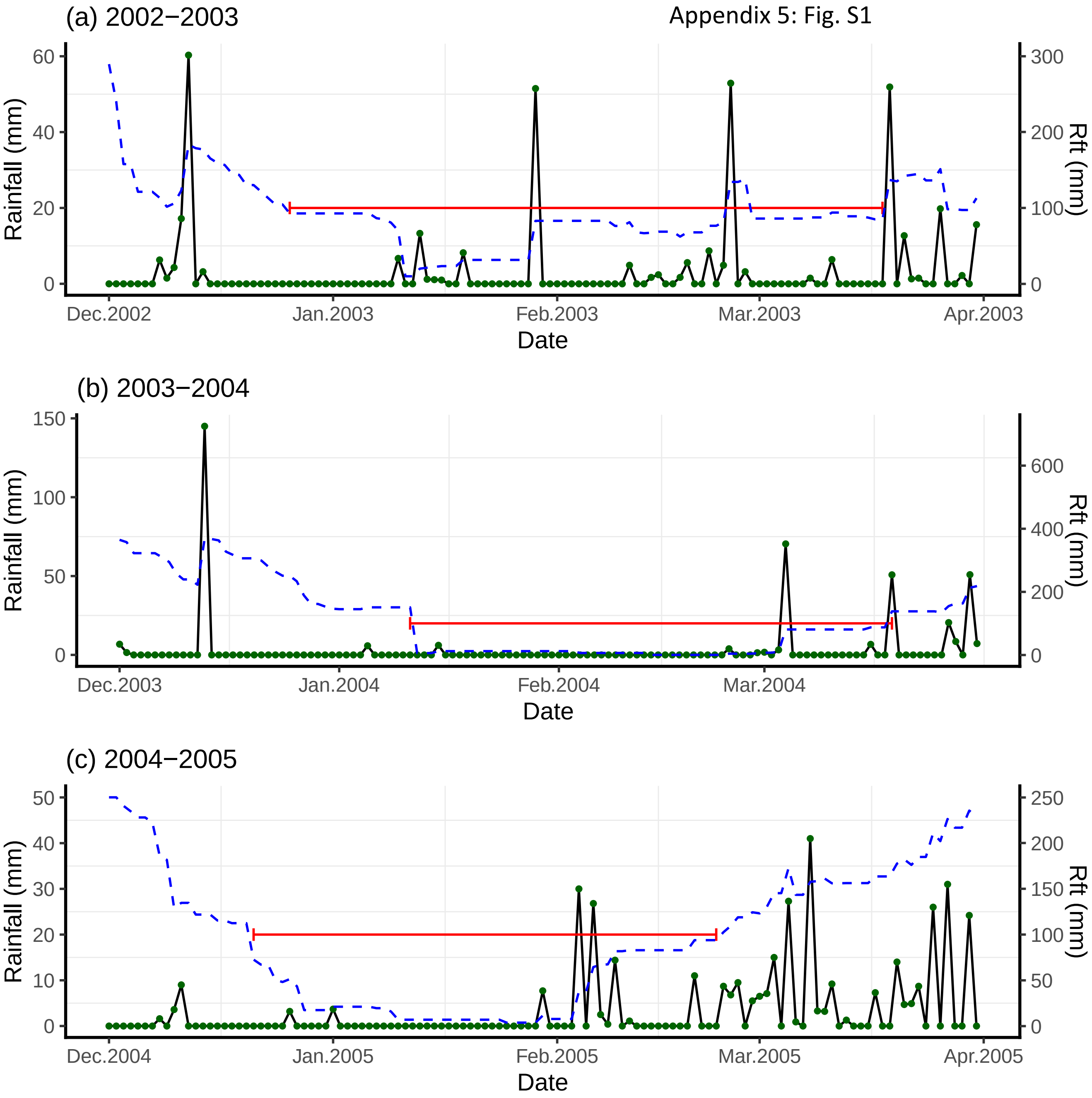

Appendix 5: Fig. S2

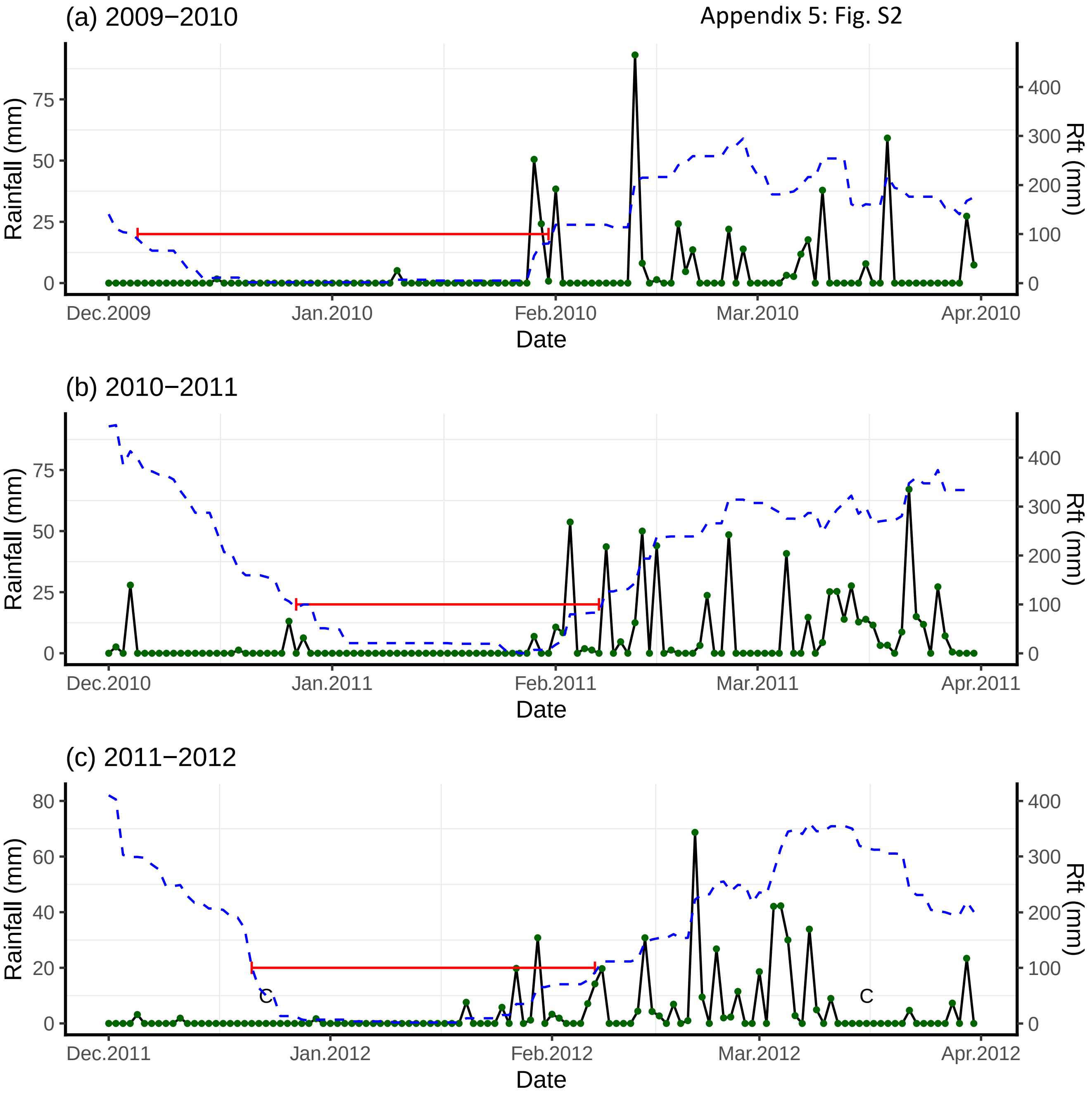

**Appendix 6:** Description of the yellow-green caterpillar feeding in 2014.

On the 27 and 28 January 2014, a second caterpillar, this one yellow-green with thin, black longitudinal stripes (Fig. S1(a)) This caterpillar, however, seems to have eaten buds as well as flowers, as encountered fresh or shrivelled on understory *Cola* leaves (Fig. S1(a) and (b) close-up). They may also have fed, perhaps secondarily to complete instar development, upon recently expanded/mature leaflets of *M. bisulcata* from their new leaf flush in the preceding December. Occasionally the caterpillars fell—perhaps propelled downward by winds or rare rainfall—below the crowns and into the understory, where they did not eat *M. bisulcata* seedlings. No hanging silky lines were seen, but frass, webbing and fallen caterpillars were evident on the ground (inset of Fig. S1(a)). This speculation is based on the pointed sharp-looking protruding mandibles visible in Fig. S1(b). If the photographed caterpillars (27 January 2014) were late instars (nearly ready to pupate), then their herbivorous activity in crowns may have even begun 2–3 weeks prior.

The yellow-green caterpillar is non-hairy, with a pronounced spherical head cinnamon-orange in color and what look like 3 or 4 pairs of usable prolegs (Fig. S1(b)); another smaller ventral pair look degenerate and unsuitable for gripping). Fully extended, its body is likely longer than the black caterpillar described above, and its locomotion is more wave-like than loop-like (Fig. S1(b)). Given its features (and lack of others, such as a forked tail or head), the species is almost certainly a moth, suspected to belong to the Catocalinae or Calpinae subfamily of the Noctuidae or Eribidae family, respectively. Its head/body/striation pattern bear some resemblance to *Avatha discolor* (http://lepidoptera.butterflyhouse.com.au/ereb/discolor.html) yet this and all other species but one in that genus are not found in Africa, strongly pointing instead to another *Achaea* sp. (albeit neither *A. catocaloides* nor *A. lienardi*) given the distinctive red protruding knobs on the rear dorsal end of the caterpillar's body (Fig. S1(b)).

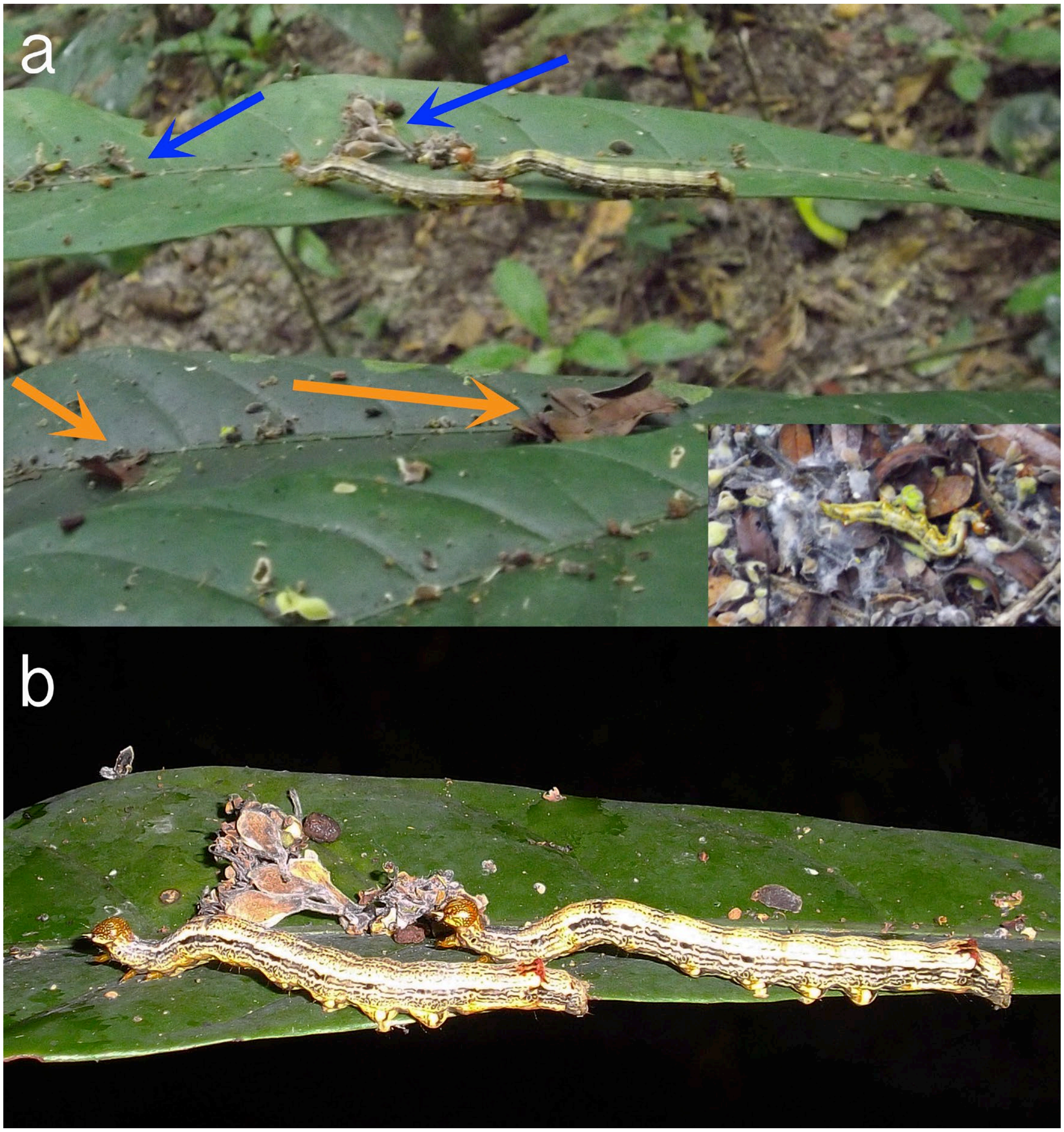


**Fig. S1.** Photographs of unknown caterpillar morphotype active in the crowns of *M. bisulcata* in January 2014 at Korup National Park, Cameroon. In (a), the blue and orange arrows respectively point to flowers and pinnate leaflets from crowns of *M. bisulcata* trees; and the lower-right inset image is of a fallen caterpillar in the litter amid webbing and shed *M. bisulcata* leaflets. Shown in (b) is a close-up of the same two caterpillars (on the same leaf) that was taken just a few seconds later using flash photography. Photo credit: J. M. Norghauer.

#### Appendix 7: PAMOL yield data 2003-2016 analysis

Yield data (oil-palm nuts, as t/ha) were restricted to Jan 2003 to Dec 2016 for the entire Ndian Estate (five divisions, including Mana which borders Korup National Park) and matched up with monthly mean radiation ($W/m^2$) and rainfall recorded at the Bulu station; 2017 was dropped because the climate data are incomplete for that year. The trend in yield over the time span was weakly negative and only marginally significant ($t$ = 1.755, df = 1, 166; $P$ = 0.081). Using a GLS regression with an AR (1) term, significance was lost when autocorrelation ($r$ = 0.719) was taken into account ($t$ = –0.577, df = 1, 166; $P$ = 0.565). Factoring in season (= month) did not improve the fit ($t$ = –0.613, df = 1, 155; $P$ = 0.541).

Radiation declined significantly with time ($t$ = –2.671, df = 1, 166; $P$ = 0.0083), but using a GLS regression with an AR (1) term ($r$ = 0.678) it decreased in significance ($t$ = –1.189, df = 1, 166; $P$ = 0.236). Factoring in month for radiation improved the fit considerably ($t$ = –3.020, df = 1, 155; $P$ = 0.0030). Regressing yield on radiation, with GLS and an ARMA (2,2) term, and factoring in again month, led to an insignificant (positive) effect of radiation ($t$ = 0.519, df = 1, 154; $P$ = 0.605). Therefore, across years, differences in mean radiation did not influence oil-palm yield. Replacing current monthly radiation by radiation lagged by 1 to 6 months in the past led to a positive effect of (marginal) significance for only a lag of 2 months ($t$ = 1.956, df = 1, 154; $P$ = 0.052): for the other lags $P$ was between 0.32 and 0.82. Within each of the phenological quarters of the year (DJF, etc.) yield was insignificantly correlated with radiation in quarters 1, 3 and 4 ($P$ = 0.61 to 0.97), although it was marginally positively correlated in quarter 2 (early wet [MAM], $P$ = 0.087). The time between anthesis to full nut maturity and harvesting is 5-6 months; 2 months before harvest the palm oil starts to build up (Corley and Tinker 2016)[1].

[1] Reference given in main paper.

Average rainfall, in contrast to radiation, did not change significantly with time ($t$ = 0.789, df = 1, 166; $P$ = 0.431), and applying the GLS regression and AR (1) term (autocorrelation $r$ = 0.680) this did not improve the outcome ($t$ = 0.322, df = 1, 166; $P$ = 0.748). Factoring in month for rainfall had little consequence either ($t$ = 1.006, df = 1, 155; $P$ = 0.316). Regressing yield on rainfall, with GLS and the ARMA (2,2) term, and factoring in month, led to an insignificant (negative) effect of rainfall on year-to-year yield differences ($t$ = –0.616, df = 1, 154; $P$ = 0.539). Thus, across the years mean rainfall also did not influence oil-palm yield. Again, replacing current monthly rainfall by rainfall lagged by 1 to 6 months in the past led to a now positive effect of (marginal) significance for a lag of 1 month ($t$ = 1.792, df = 1, 154; $P$ = 0.075), yet a negative effect at a lag of 4 months ($t$ = 1.736, df = 1, 154; $P$ = 0.085): for the other lags $P$ was between 0.10 and 0.47. Monthly yield means exhibited the expected annual trends, with a peak in April and a low in September-October (Appendix 7: Fig. S1). A lag of 4 months is 1 month after anthesis, a lag of 1 month is 1 month before harvesting.

Nevertheless, masting events in Mb (M [= strong] for 2004, 2007 and 2010; m [= moderate] for 2013, 2014 and 2016) showed no correspondence with mean annual oil-palm yields, in the same years of fruiting (ANOVA: 'M', vs 'm', vs 'none'; $F$ = 0.03, df = 2,11; $P$ = 0.97). One proviso is the assumption that across the whole estate age distributions of palms were similar from year to year. This was likely so because the means are taken over all divisions and the ~22-year rotation is aimed at a roughly steady output over time. In the analysis of oil palm yields (adjusted with an age-yield regression), the mean yield was 8.37 t/ha (1976-1995), a value higher than from the present analysis for 2003-2016 of 5.98 t/ha because the latter included young stands (< 6 yr old) which were non-yielding. As before there was no clear correspondence between oil nut production and mast fruiting of *M. bisulcata*.

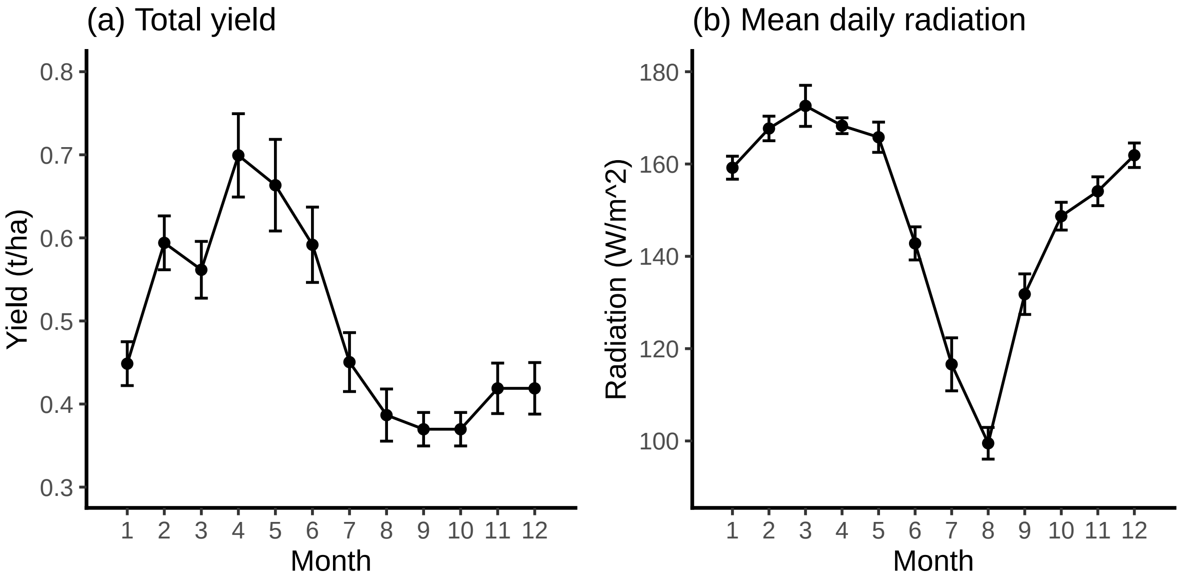


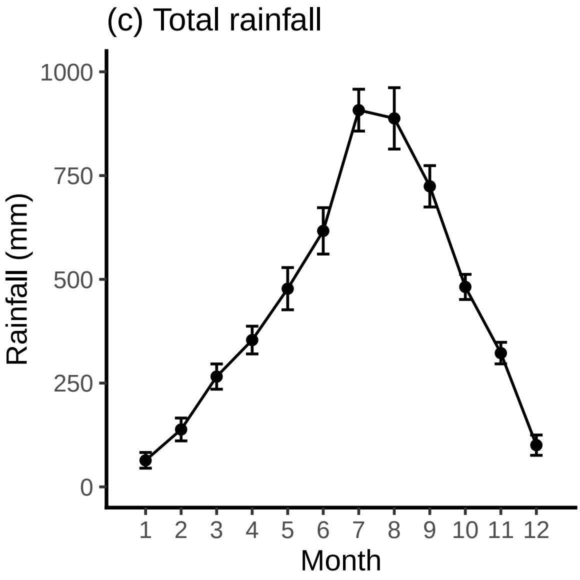


**Appendix 7: Fig. S1.** Annual trends in oil-palm yield at the Ndian Estate, 2003-2016, and in radiation and rainfall recorded at the Bulu station.

The interested reader should contact C. Etta at PAMOL Plc if they wish to have access to the yield data set.

**Appendix 8: Fig. S1.** Mean and SD stem radial increments ($n$ = 20 trees) in relation to mast fruiting of *Microberlinia bisulcata* in the P-plot: (a) time series of mean where 'M' indicates a full masting, (b) SD versus mean annotated by year.

(a)

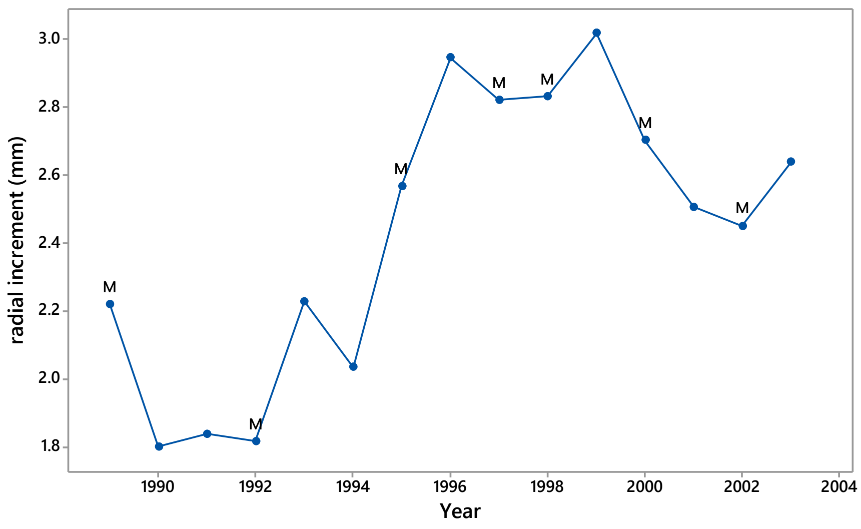


(b)

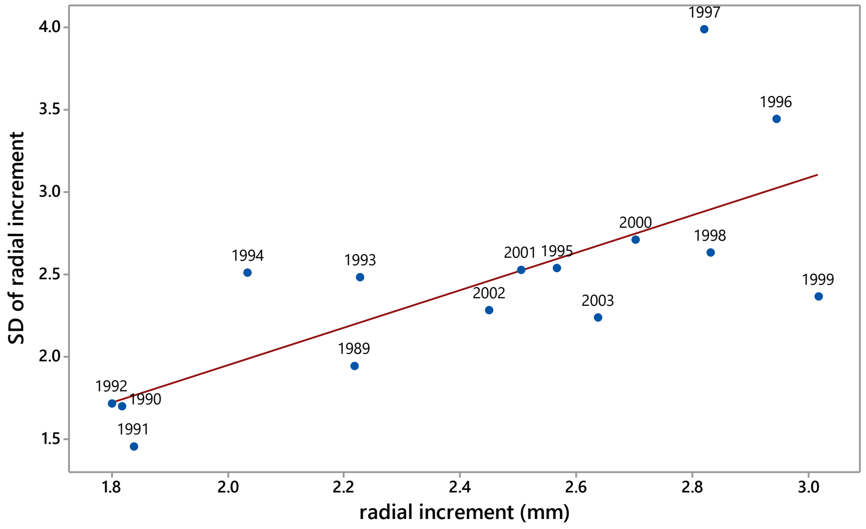

**<u>Appendix 9:</u>** <u>Minimum temperature and mast fruiting</u>

The number of the first day with the absolute minimum value of minT (*fdmin*, days; range 9-62), or run of consecutive equal values (1 day increase in a run allowed) for each quarter-year-defined (DJF) dry season was not significantly correlated with the start day of the drought-defined dry season (both dates variables from 1 December), or its duration ($r$ = 0.245 and −0.121, df = 27, P ≥ 0.12 respectively). Minimum minT showed a strong negative relationship to *fdmin* ($r$ = -0.551, df = 27, P = 0.002; ESM, Appendix 9, fig. S3). The later the drop when minT occurred, the stronger that drop with *fdmin* (value changing on average from c. 22 to 16 deg. C). Outside of the DJF seasons, minT fell to < 20 deg. C on only three isolated days in 29 years. Minimum minT was less strongly negatively correlated, however, with the time of the temperature drop into the drought-defined dry season (*fdmin* minus start day; $r$ = −0.355, df = 27, P = 0.059; ESM, Appendix 9, fig. S4). Importantly, in no year was the identified minimum minT preceded by other drops of < 20 deg. C. Thus, the further into the dry season a significant minT occurred the stronger it was, simply reflecting the intensity of the dry season and dissociating it further from any flowering occurring at the start of the dry season after leaf exchange.

Mean and maximum minT for the moderate ('m') mast fruiting were significantly higher than for the strong ('M') and non-masting years, but this was only slightly so for minimum minT (ESM, Appendix 9, Table S1). Differences between mean and minimum minT were very similar for all masting classes. The difference for the 'm' years arose because they occurred at the end of time series of masting events (2013-2017), when the minT at Korup had being steadily rising over 30 years (1987-2017) by 0.8 deg. C per decade (Etta *et al*. 2022). However, strong mast fruiting occurred in 2007 and 2010, when minimum minT was already high. Day number of minimum minT did not differ significantly between masting classes, nor did the start or duration of the dry season, and hence also not the difference *fdmin*−start (ESM, Appendix 9, Table S1).

**Appendix 9: Fig. S1**. Change in (a) mean (*mn minT*), and (b) minimum (*min minT*), of minimum daily temperature recorded at the Bulu, Ndian, climate station for the drought-defined dry seasons at Korup for the years 1989 to 2017. 'M' and 'm' indicate 'strong' and 'moderate' masting years respectively.

(a)

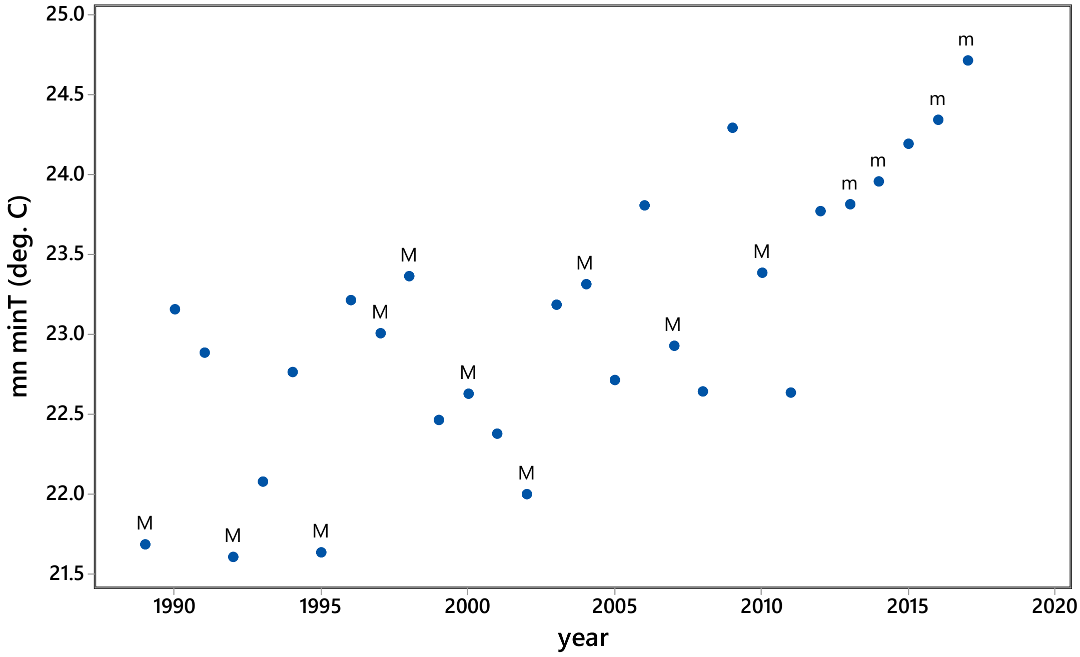


(b)

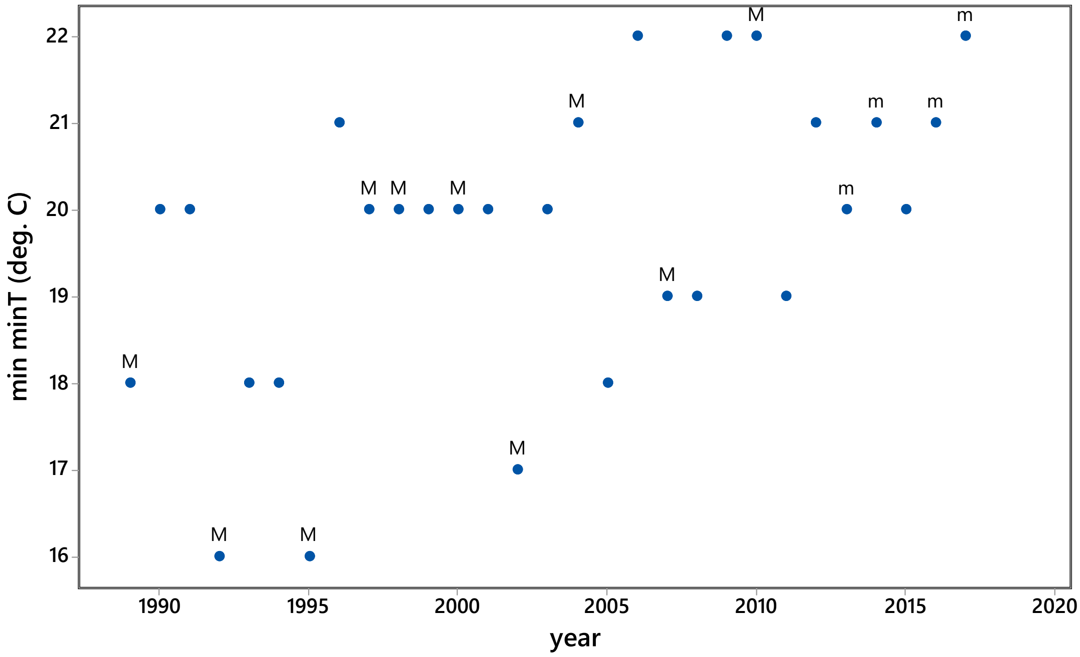

**Appendix 9: Fig. S2.** The course of daily minimum temperature (*minT*, deg. C) in the quarter-year defined dry season (i.e. months December to February) of each phenology year at Korup as recorded at the Bulu, Ndian, climate station for 1989 to 2017. 'M' and 'm' following year date in the panel headings, indicate 'strong' and 'moderate' masting respectively, no lettering means a non-masting year.

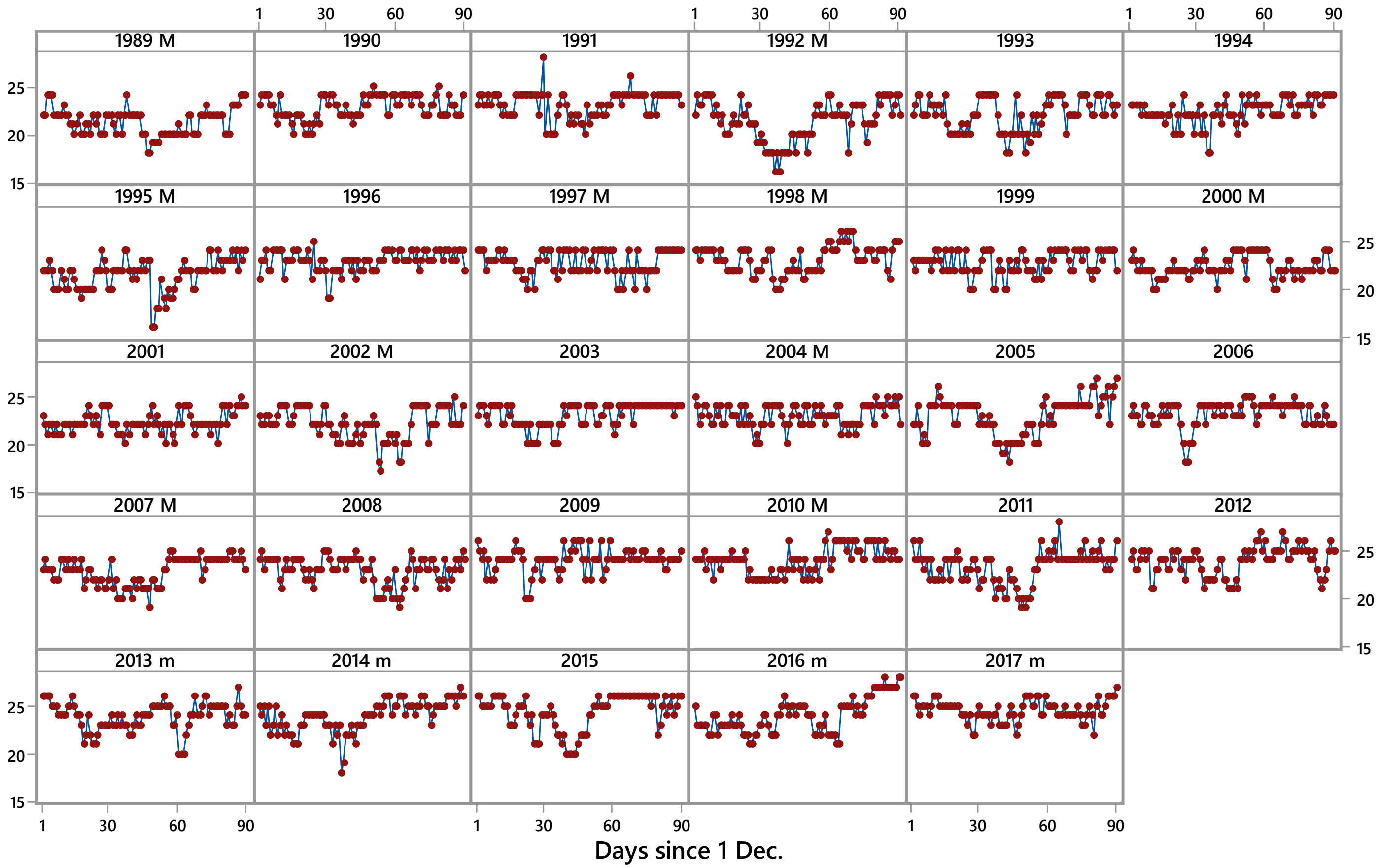

**Appendix 9: Fig. S3**. Relationship between the minimum of minimum daily temperature (*min minT*) recorded at the Bulu, Ndian, climate station and (a) the number of days into the quarter-year-defined dry season at which *min minT* first dropped to its lowest value (*fdmin*) as in Fig. S2, and (b) the difference between *fdmin* and start day of the drought-defined dry season at Korup, for the years 1989 to 2017. 'M' and 'm' indicate 'strong' and 'moderate' masting years respectively.

(a)

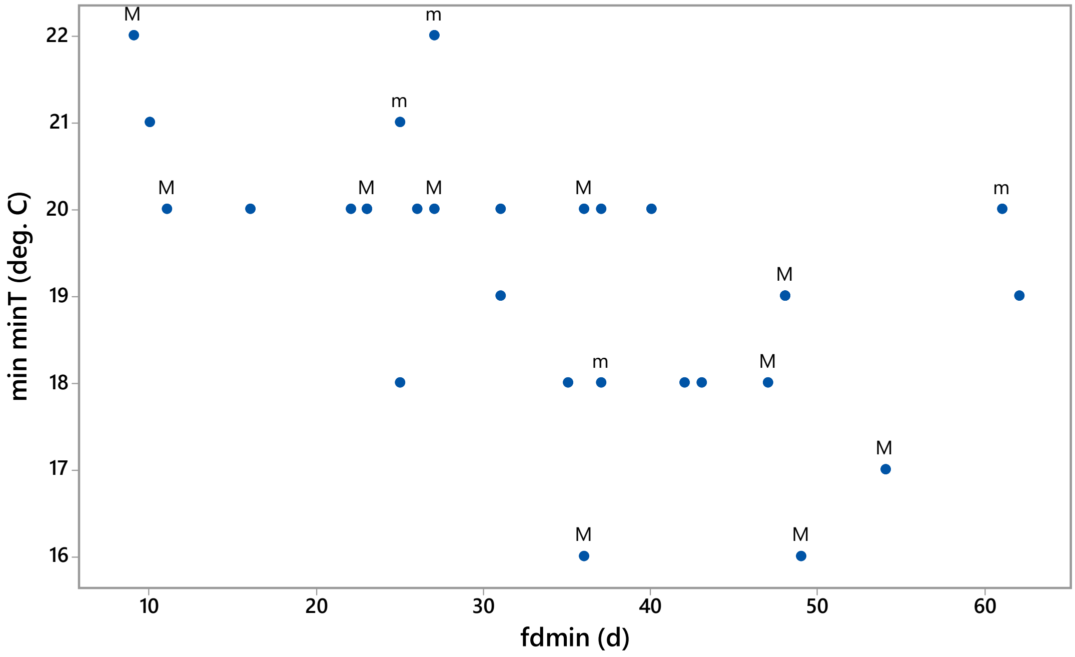


(b)

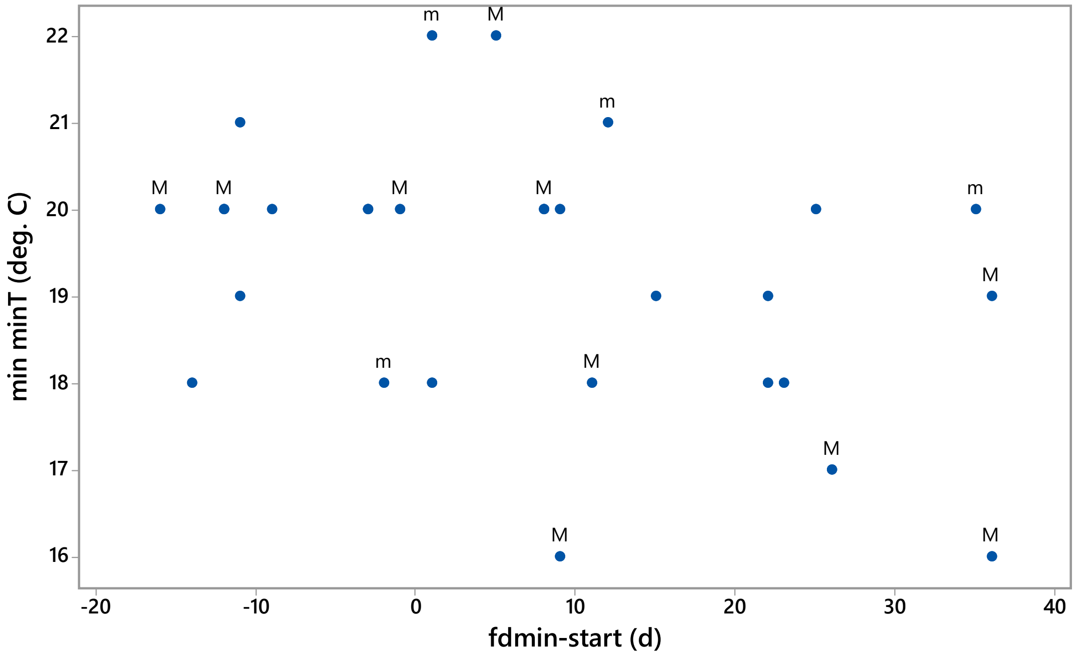

**Appendix 9: Table S1.** Minimum temperatures and their timing in the quarter-year-defined dry season (i.e. months December to February of each phenology year at Korup as recorded at the Bulu, Ndian, climate station, for strong ('M'), moderate ('m') and no masting ('none') years between 1989 and 2017. (a) Mean, minimum and maximum minimum temperatures (mn_, min_ and max_minT) and the mn_– min_mnT difference; (b) Number of days in the quarter-year-defined dry season at which min_minT first dropped to its lowest value (*fdmin*), the start day and the duration of the drought-defined dry season (ds), and the difference between *fdmin* and that start day.

| mast (a) | n | mn_minT | min_minT | max_minT | mn_– min_minT |
|---|---|---|---|---|---|
| none | 15 | 23.05 ± 0.16 | 19.33 ± 0.25 | 25.53 ± 0.36 | 3.72 ± 0.22 |
| m | 4 | 24.09 ± 0.15 | 20.25 ± 0.85 | 27.25 ± 0.25 | 3.84 ± 0.75 |
| M | 10 | 22.42 ± 0.28 | 18.80 ± 0.63 | 24.80 ± 0.33 | 3.62 ± 0.40 |
| F-value (P[F])[1] | | 8.37 (0.002) | 1.38 (0.27) | 5.86 (0.008) | 0.06 (0.94) |

| (b) | | fdmin | start ds | duration ds | fdmin – start ds |
|---|---|---|---|---|---|
| none | 15 | 32.7 ± 3.4 | 28.6 ± 2.9 | 59.9 ± 6.6 | 4.1 ± 3.7 |
| m | 4 | 37.5 ± 8.3 | 26.0 ± 5.3 | 62.5 ± 16.0 | 11.5 ± 8.4 |
| M | 10 | 34.0 ± 5.1 | 23.8 ± 3.7 | 78.2 ± 3.5 | 10.2 ± 5.7 |
| F-value (P[F])[1] | | 0.17 (0.85) | 0.54 (0.59) | 2.01 (0.15) | 0.59 (0.56) |

[1] df = 2,26
